# Dynamic Generalised Additive Models (DGAM) for forecasting discrete ecological time series

**DOI:** 10.1101/2022.02.22.481550

**Authors:** Nicholas J Clark, Konstans Wells

**Author notes:** Corresponding Author: Nicholas J Clark, School of Veterinary Science, The University of Queensland, Gatton QLD 4343 Australia.

## Abstract

1. Generalised Additive Models (GAMs) are increasingly popular for describing smooth nonlinear relationships between predictors and response variables. GAMs are particularly relevant in ecology for representing hierarchical functions for discrete responses that encompass complex features including zero-inflation, bounding and uneven sampling. However, GAMs are less useful for producing forecasts as their smooth functions provide unstable predictions outside the range of training data.
2. We introduce Dynamic Generalised Additive Models (DGAMs), where the GAM linear predictor is jointly estimated with unobserved dynamic components to model time series that evolve as a function of nonlinear predictor associations and latent temporal processes. These models are particularly useful for multivariate series, as they can estimate hierarchical smooths while learning complex temporal associations via dimension-reduced latent factor processes. We implement our models in the *mvgam* R package, which uses the *mgcv* and *rjags* packages to construct smoothing splines and estimate unobserved parameters in a probabilistic framework.
3. Using simulations, we illustrate how our models outperform competing formulations in realistic ecological forecasting tasks while identifying important smooth predictor functions. We use a real-world case study to highlight some of *mvgam’s* key features, which include functions for: calculating correlations among series’ latent trends, performing model selection using rolling window forecasts, online data augmentation via a recursive particle filter, and visualising probabilistic uncertainties for smooths and predictions.
4. Dynamic GAMs (DGAM) offer a solution to the challenge of forecasting discrete time series while estimating ecologically relevant nonlinear predictor associations. Our Bayesian latent factor approach will be particularly useful for exploring competing dynamic ecological models that encompass hierarchical smoothing structures while providing forecasts with robust uncertainties, tasks that are becoming increasingly important in applied ecology.

## INTRODUCTION

Rapidly changing climates and landscape modification are impacting species and ecosystems at all micro- and macroecological levels, incurring substantial economic and environmental costs (World Health Organization 2005, United Nations 2015, Kennedy et al. 2019). There is broad consensus among scientists, parliamentarians and applied decision-makers that anticipating probable future states is vital to mitigate impacts of environmental change on ecosystem functionality and services (Schmidt et al. 2010, Dietze et al. 2018, Intergovernmental Panel on Climate Change 2018).

Two challenges impede the improvement and adoption of common forecasting tools in ecology. First, natural systems are driven by networks of interacting biotic and abiotic processes (Levin 1998, Choler et al. 2001, Massoud et al. 2018). These dynamic natural processes are the products of multiple sources of variation including long-term trends, seasonal and other cyclic oscillations, environmental forcing, temporal dependence or species interactions (Choler et al. 2001, Dietze 2017, Auger-Méthé et al. 2021). Second, ecological time series tend to be integer-valued variables such as observations of species presence or abundance that exhibit complex features including observation error, zero-inflation, over-dispersion, bounds, missing values and uneven sampling frequency (Lindén and Mäntyniemi 2011, Simpson 2018, Warton 2018, Kowal and Canale 2020). Moreover, ecological observations are almost always multivariate when contextual information is considered. These features make it difficult to analyse ecological time series while sufficiently accounting for the various systematic time series components and possible multivariate relationships (Auger-Méthé et al. 2021).

Time-series analyses are often concerned with decomposing temporal variation into components representing trend, seasonality, and other cyclic changes. Generalised Additive Models (GAMs), which are increasingly used in ecology to identify nonlinear relationships (Guisan et al. 2002, Hughes et al. 2018, Simpson 2018, Pedersen et al. 2019), offer a convenient way to accomplish this decomposition. Outlined in detail previously (Hastie and Tibshirani 1990, Wood 2004), GAMs can briefly be described as modified Generalised Linear Models (GLMs) in which the linear predictor includes a sum of smooth functions representing functional relationships between covariates and the response:

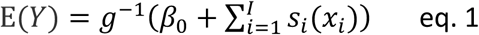

where E(*Y*) is the conditional expectation of a response assumed to be drawn from distribution within the exponential family, *β*_0_ is the unknown intercept, the *s*_*i*_’s are a set of smooth functions over one or several predictor variables (the *x*’s) and *g* is an appropriate monotonic link function. Each smooth *s*_*i*_ is composed of spline like basis expansions whose coefficients (*β*_*i*_), which must be estimated, act as weights for the basis expansions to control the smooth’s shape. The size of the basis expansion limits the smooth’s potential complexity, with a larger set of basis functions allowing greater flexibility. In addition to their ability to represent complex and non-linear ecological relationships, several other advantages of GAMs are that they can model a diversity of response families that accommodate ecological features such as zero-inflation and that they can be formulated to include hierarchical smoothing for multivariate responses (Wood 2017, Pedersen et al. 2019).

Given the set of basis coefficients that comprise each smooth, a GAM can in principle be directly estimated as a GLM. However due to their incredible flexibility, GAMs will invariably overfit if left unconstrained (Hastie and Tibshirani 1990, Wood 2004, Marra and Wood 2011). Penalised likelihood estimation avoids this overfitting by placing quadratic penalties on the basis coefficients (referred to as smoothing penalties), which penalize the function’s ‘wiggliness’ and controlling the trade-off between fit and smoothness (Wood 2004, Wood 2016). From a Bayesian perspective, one way to represent a smooth is to draw the set of *β*_*i*_ from a multivariate Gaussian distribution with the penalty acting on the prior precision for efficient regularization. Larger penalties shrink the coefficient covariances, effectively forcing the smooth toward a straight line when the data do not justify a nonlinear function (Marra and Wood 2011, Wood 2016). GAMs are particularly sought after for modelling time series to both identify nonlinear or time-varying covariate effects and to uncover periods of rapid change, though strong temporal autocorrelation can make it challenging to estimate key parameters (Yang et al. 2012, Knape 2016, Simpson 2018, Spooner et al. 2018, Camara et al. 2021).

For many ecological studies that employ GAMs, a primary objective is predicting future states (Ward et al. 2014, Kaplan et al. 2016, Clark et al. 2020, Malick et al. 2020, Koolhof et al. 2021). However, a lingering issue in using GAMs for forecasting is the way in which smooth functions predict outside the range of training data. Many of the smooths used in ecological GAMs have zero second derivatives at the boundaries, meaning they will linearly extrapolate beyond the last observation (Elith et al. 2010, Zurell et al. 2012). This projection of a straight line indefinitely into the future can produce unrealistic forecasts, particularly if the estimated function ‘wiggles’ (i.e. exhibits a pronounced change in the response-predictor relationship) near the boundary (**Figure 1 top**). There are technical solutions to help with this problem, for example by extending the evaluation of the ‘wiggliness’ penalty into the range of values that we wish to forecast (i.e. weeks or years ahead of the training data) or by forcing the smooth to use the last observed value with fixed uncertainty intervals when forecasting by imposing a first derivative penalty (**Figure 1 bottom**). However, these modifications are insufficient to generate robust ecological forecasts with appropriate probabilistic uncertainties as they do not adequately capture the temporal dependence in the data (see examples in Appendix S1 in Supporting Information).

**Figure 1:**
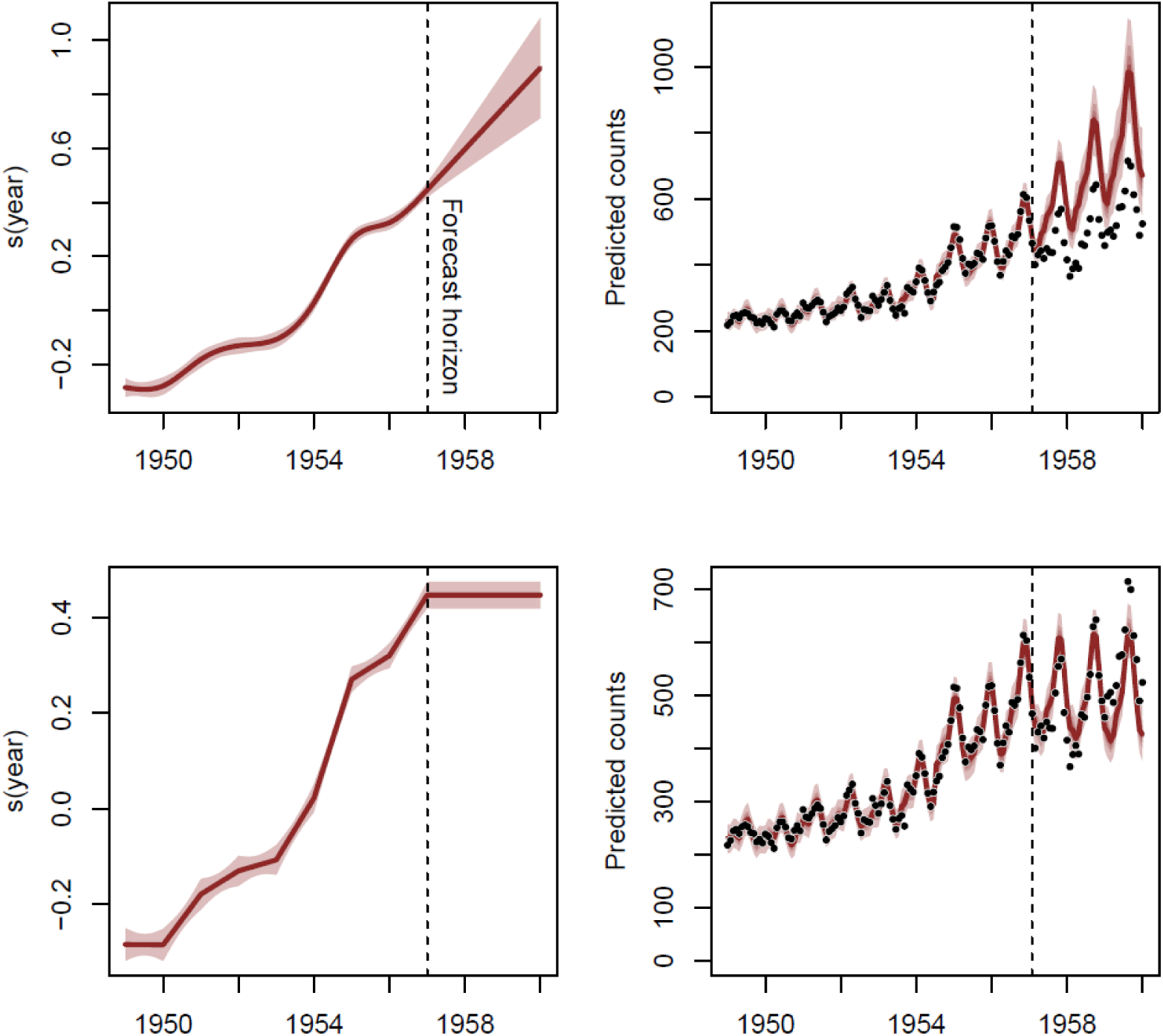
Estimated trends and forecasts from two GAMs applied to a discrete time series. In the top panel, a thin plate regression spline with a penalised second derivative is used for the trend, leading to a smooth function (top left) and linear extrapolation when forecasting (top right). In the bottom panel, the trend penalty is placed on the first derivative, resulting in flat extrapolation when forecasting. Trend shading shows 95% confidence intervals, while forecast shading shows empirical quantiles. Both models were fitted to a simulated seasonal discrete time series in R using the *mgcv* package with the general formula: *y ∼ s(year, bs = ‘tp’) + s(season, bs = ‘cc’) + ti(season, year), family = nb()*).

In this paper we outline a Bayesian Dynamic GAM (DGAM) that provides a general solution to the problem of estimating smooth functions while generating reliable forecasts for discrete time series. The approach is simple: for univariate series we augment the GAM linear predictor with a latent dynamic component (capturing possible temporal dependencies as a state-space process) to capture the series’ temporal evolution process (either as a random walk or an autoregressive process up to order 3). For modelling multivariate time series, we accommodate possible dependencies among series’ temporal components in a parsimonious way using a dynamic latent factor process. We introduce our associated R package *mvgam* (https://github.com/nicholasjclark/mvgam), which provides the following key functions:

- Estimate parameters of DGAMs in a Bayesian Markov Chain Monte Carlo framework via the Gibbs sampling software JAGS (Plummer 2003, Wood 2016)
- Perform residual diagnostic checks using randomised quantile (Dunn-Smyth) residuals (Dunn and Smyth 1996)
- Compute multivariate time series correlations
- Perform model selection using rolling window forecasts
- Update forecasts online via a Sequential Monte Carlo particle filter
- Plot posterior retrodictive and predictive checks to examine discrepancies between observed data and model-generated simulations (Gabry et al. 2019)
- Plot estimated smooths and posterior predictions, along with their probabilistic uncertainties

We begin by presenting our model, including background material for the dynamic factor process. We then illustrate our package’s utility for ecologists and other users interested in forecasting discrete time series using both simulations and a case study. An introduction to *mvgam*’s primary functions via reproducible examples is provided in the Appendices S1 – S3 (Supporting Information).

## DYNAMIC GENERALISED ADDITIVE MODELS

### Univariate models for a single ecological time series

A Bayesian framework to model fitting and parameter estimation involves defining a joint probability distribution over all observable and unobservable quantities in a statistical model that aligns with expert beliefs about the data generating process (Gelman et al. 2017). A dynamic GAM is naturally viewed from a Bayesian perspective, where prior beliefs about the nonlinearity of a function can be elicited to inform the complexity and penalisation of the smooth (Wood 2013, Miller 2019, Pedersen et al. 2019) while accounting for possible unobserved temporal dependence in line with the expectation that time series evolve as serially autocorrelated dynamic processes (Hyndman and Athanasopoulos 2018). In its basic form, the dynamic GAM is written as:

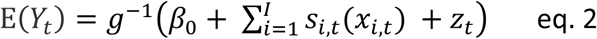

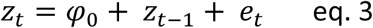

where E(*Y*_*t*_) is the conditional expectation of the response at time *t* and *z*_*t*_ is the dynamic process estimate at time *t*. In its simplest form, temporal dependence is modelled as a random walk with possible drift, where *φ*_0_ in eq. 3 is the drift parameter and the residual error *e*_*t*_ is drawn from a zero-centred Gaussian distribution with a fixed (time-invariant) standard deviation. This can easily be expanded to include autoregressive (AR) processes. For example, the following specifies a latent AR2 model:

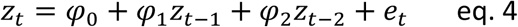

Note that while it is possible to model residual autocorrelation similarly in the R package *mgcv* using the *gamm* function (Wood 2017), there is no straightforward way to include this autocorrelation process in forecasts. Our models are coded in the JAGS probabilistic programming language using the function *mvjagam*, which relies on the *jagam* function from *mgcv* to generate a skeleton JAGS model file, smooth penalty matrices and starting values for GAM parameters (Wood 2016). The model is modified to include dynamic components and to update any prior distributions specified by the user, while all data reformatting necessary for JAGS modelling is done automatically. Employing the JAGS software through the R interface *rjags* (Plummer 2003), the model is conditioned on observed data using Markov Chain Monte Carlo (MCMC) simulation via Gibbs samplers to calculate the posterior probability distribution of the unobserved parameters of interest.

It is notable that our design permits any formula allowed in *mgcv* to be used for the GAM component of the linear predictor, providing a user-friendly way to explore dynamic ecological models that encompass nonlinear smooths. Other advantages of our framework are (1) missing values are allowed for the responses; (2) upper bounds can be used via truncated likelihoods; (3) smooth distributed lag covariate functions can be estimated alongside latent temporal components to form complex dynamic nonlinear models (Gasparrini 2011); and (4) dynamic components can easily be forecasted via their autoregressive equations, providing robust probabilistic uncertainties.

### Dynamic factor models for a set of multivariate ecological time series

Here we describe how we modify our dynamic GAM into a joint multivariate statistical model for collections of time series. Dynamic factor models that account for multivariate relationships in time series data are closely aligned with static latent factor models, which are used in quantitative ecology to jointly model multiple species by estimating shared responses to unmeasured ecological drivers (Warton et al. 2015, Thorson et al. 2016, Ovaskainen et al. 2017, Ward et al. 2021). A latent factor model is a function of unmeasured random predictors (factors) that induce correlations between responses via factor loadings while exercising dimension reduction. Often, species do demonstrate correlated responses to gradients, meaning that a smaller set of factors (i.e. a low-dimensional representation) than the total number of possible inter-species relationships can adequately capture the main axes of covariation (Letten et al. 2015, Warton et al. 2015). A dynamic factor model assumes the factors evolve as time series. The strength of this approach is that a small number of latent factors can often model the temporal behaviors of a much larger set of series. This dimension reduction simplifies the estimation and forecasting task, as only the smaller set of dynamic factors and the series’ specific factor loadings needs to be estimated to generate forecasts (De Stefani et al. 2019). In our dynamic GAM, each series’ latent trend is composed of a linear combination of these common factors:

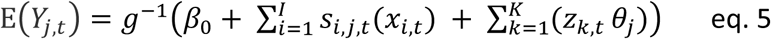

where E(*Y*_*j, t*_) is the expected response for series *j* at time *t*, the *z*_*k,t*_’s are estimates for the *K* factors at time *t* and the *θ*_*j*_’s are factor loadings. As in the univariate case, the factors can evolve either as random walks with drift or as autoregressive processes up to order 3.

A challenge with any factor model is the need to determine the number of factors *K* (Fox et al. 2009, Bhattacharya and Dunson 2011, Thorson et al. 2016, Tobler et al. 2019). Setting *K* too small prevents temporal dependencies from being adequately modelled, leading to poor convergence and difficulty estimating smooth parameters. By contrast, setting *K* too large leads to unnecessary computation. We approach this problem by formulating a prior distribution that enforces exponentially increasing penalties on the factor variances to allow any un-needed factors to evolve as flat lines. Following Welty et al (2009) and Wells et al (2016), our prior assumes that factors up to a certain threshold number π have precisions of similar magnitudes, after which they increase exponentially (leading to variances that shrink toward zero). Along with π, we estimate two other hyperparameters that control the baseline penalty and the rate at which penalties exponentially increase, respectively, allowing the data to inform the selection of dynamic factors. We caution however that setting *K* too large could result in trends that are overly flexible, making it challenging to simultaneously estimate important smooth functions such as seasonality. It is certainly worth checking whether inferences or forecasts are sensitive to *K*, perhaps using the guidelines outlined by Tobler *et al* (2019). Additional constraints preserve identifiability by setting the upper triangle of the factor loading matrix to zero and ensuring non-negative diagonals (Hui 2016, Tobler et al. 2019).

## SIMULATIONS

We used simulations to examine the performance of our model. Briefly, we simulated multivariate datasets with 72 time points (six years of data for monthly series) consisting of negative binomial observations (size = 1) for sets of series whose log linear predictors included a hierarchical seasonal pattern (where each series’ seasonal pattern was created by drawing from a global seasonal pattern with common Gaussian noise; see function *sim_mvgam()* in the mvgam package) and correlated latent trends. Temporal dependences followed a dynamic factor process in which factors evolved as random walks with drift and series’ loadings were drawn from a zero-centred multivariate Gaussian with a sparse covariance matrix. We investigated model sensitivity to missingness (proportion missing = 0, 10, or 50), dimensionality (number of series = 2, 4 or 12) and the magnitude of the temporal component relative to seasonality (0.3, for moderate dynamics, or 0.7 for a strong dynamics). Each simulated dataset was fit with the same set of three models. First, we fit a hierarchical *gam* using *mgcv* that included a random intercept per series (*s(series, bs = ‘re’)*), a cyclic smooth for global seasonality (*s(season, m = 2, k = 12, bs = ‘cc’)*), a local smooth for series-specific deviations from global seasonality (*s(season, series, m = 1, k = 4, bs = ‘fs’)*) and independent non-wiggly annual smooths (*s(year, by = series, m = 1, k = 3, bs = ‘gp’)*). We next fit a DGAM (with *K* = half the number of series) with identical random effect and seasonal smooths but no yearly smooth. Finally, we fit a ‘null’ DGAM that only estimated random intercepts. Negative binomial distributions were specified for each model and random walks were used for DGAM dynamic factor processes. Each combination of missingness, dimensionality and dynamic strength was used to generate five replicate datasets, yielding a total of 60 simulations. For *mgcv* models, estimation of smoothing penalties was performed using restricted maximum likelihood (*method = ‘REML’*). Weakly informative Gaussian priors were specified for AR parameters (*φ*) (mean = 0; variance = 0.1). Following Wood (2016), zero-centred multivariate Gaussian priors were used for each smooth’s *ß* parameters and exponential priors were used for the smoothing penalties. For *mvgam* models we ran two MCMC chains for 10,000 iterations as burnin and collected 1000 samples from the joint posterior.

The relative performances of each model were explored using out of sample forecasts. We trained models on the first five years of data (60 observations) and generated forecasts for the remaining year (12 observations). Probabilistic forecast performance was evaluated using a discrete version of the Rank Probability Score (DRPS; Gneiting and Raftery 2007) and coverage of 90% prediction intervals. Forecasts with lower DRPS and coverages closer to 0.9 were considered more accurate.

## CASE STUDY: FORECASTING TICK ABUNDANCES

*Amblyomma americanum* and *Ixodes scapularis* are two widespread species of hard ticks capable of transmitting a diversity of parasites to animals and humans, many of which are zoonotic (Rochlin and Toledo 2020). Due to the medical and ecological importance of these species, a crucial goal in disease ecology is to understand factors that influence their abundances over time. The National Ecological Observatory Network (NEON) carries out standardised long term monitoring of tick abundances as well as other important indicators of ecological change (Thorpe et al. 2016). Nymphal abundances of both tick species are routinely recorded across a series of NEON plots by drag cloth sampling, with plots nested within sites (Springer et al. 2016). These plot-level series show strong seasonality and incorporate many of the challenging features associated with ecological data including overdispersion, high proportions of missingness and irregular sampling, making them useful for exploring the utility of *mvgam*.

Temperatures between −5°C to 5°C can affects various components of tick physiological diapauses and host-seeking behaviours (Clark 1995). We therefore included a cumulative growing degree day (cum_gdd) variable using temperature records for each site’s nearest weather station from NOOA’s Daily Global Historical Climatology Network daily database as a covariate. The predictor was calculated as the total number of days up to the start of the tick season (1st June) in which the mean of the day’s maximum and minimum temperatures was above 0°C. We fit species-specific DGAMs to four years of data (2015 – 2018) for 17 *A. americanum* plots (nested in 7 NEON sites) and for eight *I. scapularis* plots (nested in three sites). Timepoints during winter (epidemiological weeks 1 – 14 and 41 – 53) were removed prior to modelling as no observations occurred during this period, leaving a seasonal frequency of 26 (calendar weeks 15 – 40). For each species we fit four models representing different hypothetical dynamics:

- Null: there is no seasonality, rather the latent factors / random site-level effects of cum_gdd fully influence the dynamics for the plot-level series. Formula: *y ∼ s(site, bs = ‘re’) + s(cum_gdd, site, k = 3, bs = ‘fs’) + Z*
- Hyp1: all plots share a seasonal pattern, with any remaining variation captured by the latent factors and site-level cum_gdd effects. Formula: *y ∼ s(site, bs = ‘re’) + s(cum_gdd, site, k = 3, bs = ‘fs’) + s(season, k = 26, m = 2, bs = ‘cc’) + Z*
- Hyp2: as above but with hierarchical seasonality, including a global seasonality smooth and a seasonal smooth that can deviate across each site. Formula: *y ∼ s(site, bs = ‘re’) + s(cum_gdd, site, k = 3, bs = ‘fs’) + s(season, k = 26, m = 2, bs = ‘cc’) + s(season, site, m = 1, k = 6, bs = ‘fs’) + Z*
- Hyp3: as above but the deviations occur at the bottom level of aggregation (plot rather than site level). Formula: *y ∼ s(site, bs = ‘re’) + s(cum_gdd, site, k = 3, bs = ‘fs’) + s(season, k = 26, m = 2, bs = ‘cc’) + s(season, plot, m = 1, k = 4, bs = ‘fs’) + Z*

We used random walk dynamic factor models (*K* = 8 for *Ixodes* and 10 for *Amblyomma*) for the temporal evolution and assumed a Negative Binomial distribution for the observations. Each model was estimated using two MCMC chains for 10,000 iterations as burnin. We collected 2,000 posterior samples to evaluate parameter estimates and inspect forecasts. The 2019 observations for each plot were held out as testing data to evaluate model forecasts using the same evaluation criteria as in the simulations above.

## RESULTS

### DGAM forecast performance - simulation results

Our simulations explored the relative forecast performance of DGAMs (using *mvgam*) models versus static GAMs (using *mgcv*). The seasonal DGAM outperformed its GAM counterpart in terms of DRPS, providing better probabilistic forecasts in all comparisons (Figures 2; Figure S1). As expected, the correctly specified seasonal DGAM was the best performer when the dynamics were moderate compared to the seasonal magnitude, while the nonseasonal DGAM performed equally well under strong dynamics (Figure 2). The seasonal GAM was the worst performer in nearly all comparisons, though it did perform markedly better than the mis-specified nonseasonal DGAM under moderate dynamics and with no missing observations (Figure 2). Results were similar when inspecting DRPS as a function of the number of series, with no comparisons favouring the GAM over the DGAMs (Figure S1).

**Figure 2:**
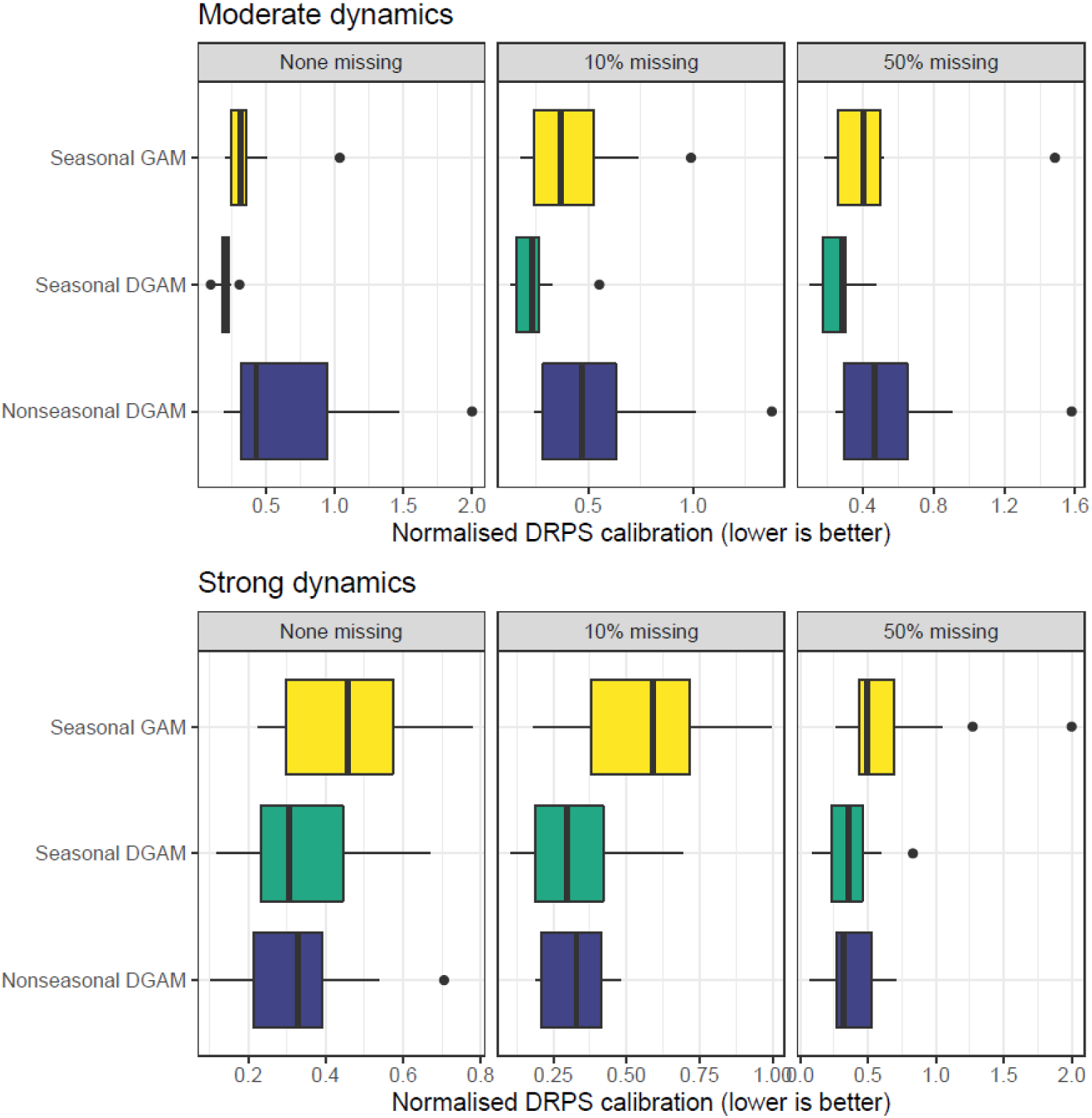
Normalised Discrete Rank Probability Score (DRPS) performance for out of sample forecasts from competing models fitted to sets of simulated discrete time series. Panels depict models fitted with different levels of data missingness (proportion of observations set to NA) and temporal dynamics strength. The Seasonal GAM was fitted using R package *mgcv*, while the Seasonal and Nonseasonal DGAMs were fitted using the *mvgam* package. Lower scores indicate better model performance.

Comparisons of 90% interval coverages strongly favoured the two DGAMs (Figure 3). Intervals for the DGAMs frequently included 25 – 35% more of the out of sample observations than did the GAM intervals. There was little distinction between the two DGAMs, even as the number of series and the strength of the underlying dynamics increased (Figure 3). Results were similar when inspecting 90% interval coverage as a function of missingness, with the DGAMs strongly outperforming the GAM (Figure S2).

**Figure 3:**
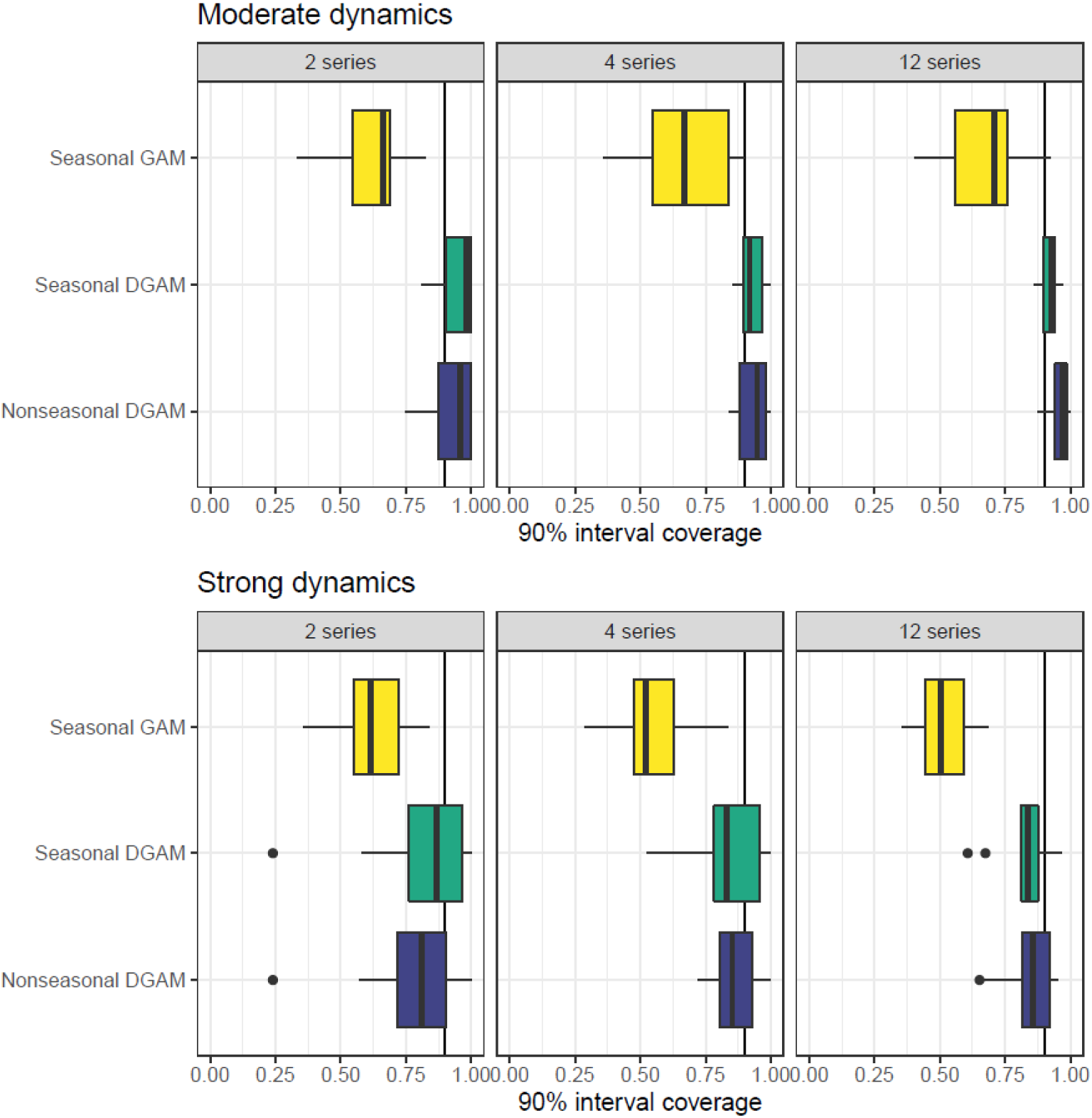
90% interval coverage for out of sample forecasts from competing models fitted to sets of simulated discrete time series, plotted as a function of dimensionality (total number of series) and dynamics strength. The vertical line in each plot marks a coverage of 0.9. The GAM was fitted using R package *mgcv*, while the DGAMs were fitted using the *mvgam* package. Scores closer to 0.9 are better.

### DGAM and NEON tick abundance forecasts

Our results suggested that Hyp3, which captured hierarchical seasonality by allowing individual plot-level seasonal patterns to deviate from global seasonality, was the best-performing model when forecasting *I. scapularis* nymphal abundance across NEON sites, while the null model that did not include seasonality was the worst-performing (Figure 4). Nominal coverages of 90% intervals were too high for most models (ranging from 92 – 98%), suggesting forecast intervals were generally wider than they needed to be (Figure 4). However, there was variation across plots in terms of forecast performance, suggesting that an ensemble forecast could improve importance (Figure S3). Inspection of PIT histograms revealed that all models apart from the null tended to overpredict to some degree (Figure S4). When conditioning on seasonality and the trend, *I. scapularis* abundances demonstrated a weak association with variation in cumulative growing degree days, with abundances tending to increase with increasing number of growing days prior to the tick season (Figure 5). Inspection of the latent dynamic components for the three seasonal models revealed strong positive within-site correlations for sites SCBI and SERC (Figure S5). Figure 5 shows example *mvgam* visualisations for a single plot, including estimated smooth functions, forecasts and dynamic trend estimates (along with their estimated uncertainties). Example *mvgam* visualisations of posterior checks for training (retrodictive) and forecast periods (predictive) are shown in Figure S6, which can indicate if a model is able to simulate time series that resemble key aspects of the observed data without notable discrepancies.

**Figure 4:**
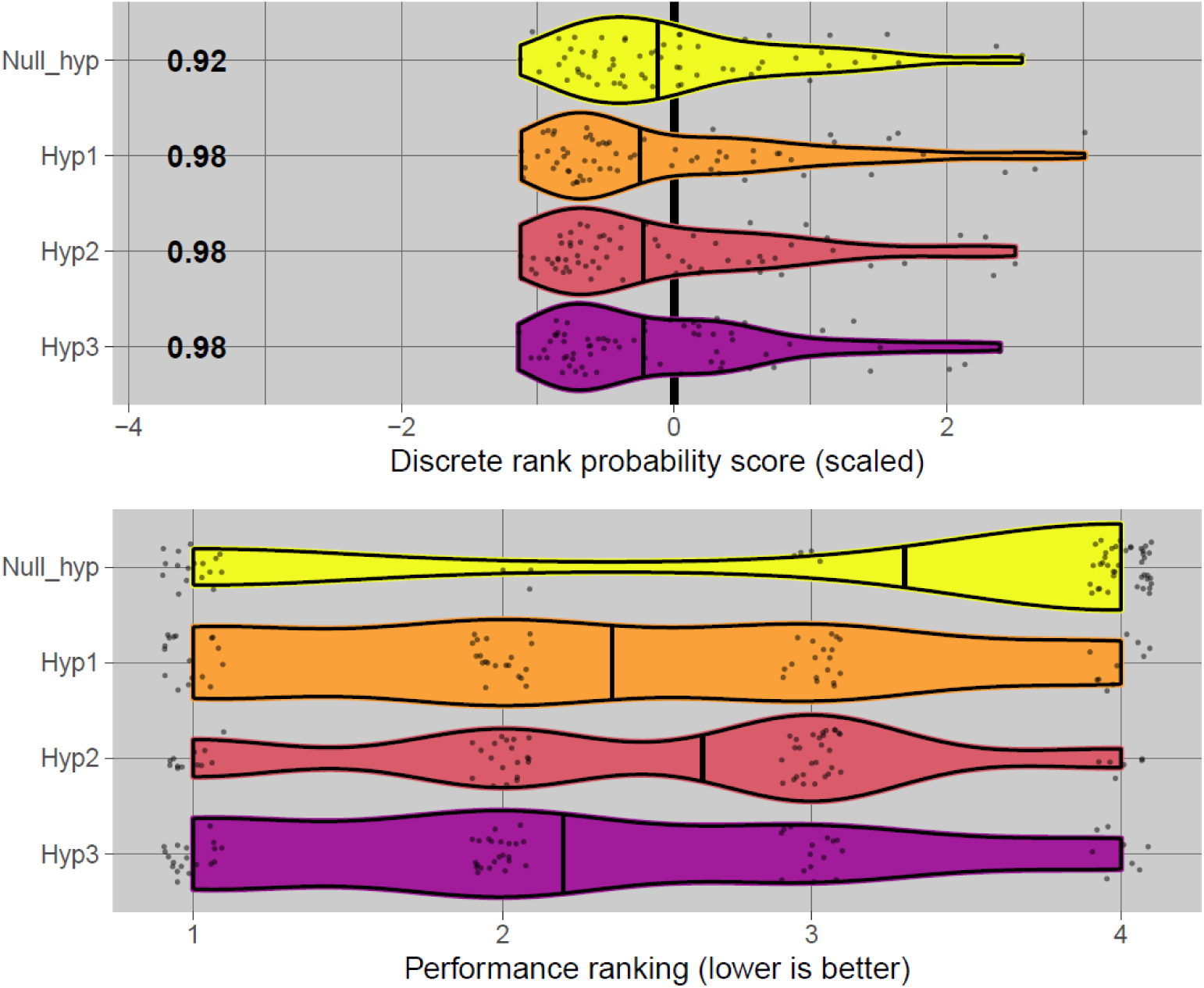
Forecast performance rank distributions based on out of sample Discrete Rank Probability Score for four competing models fitted to NEON’s *Ixodes scapularis* abundance series. Numbers on the left-hand side of the top plot indicate coverages of 90% posterior predictive intervals. Thick black lines show medians. Hypothesis definitions are outlined in section **CASE STUDY: FORECASTING TICK ABUNDANCES**.

**Figure 5:**
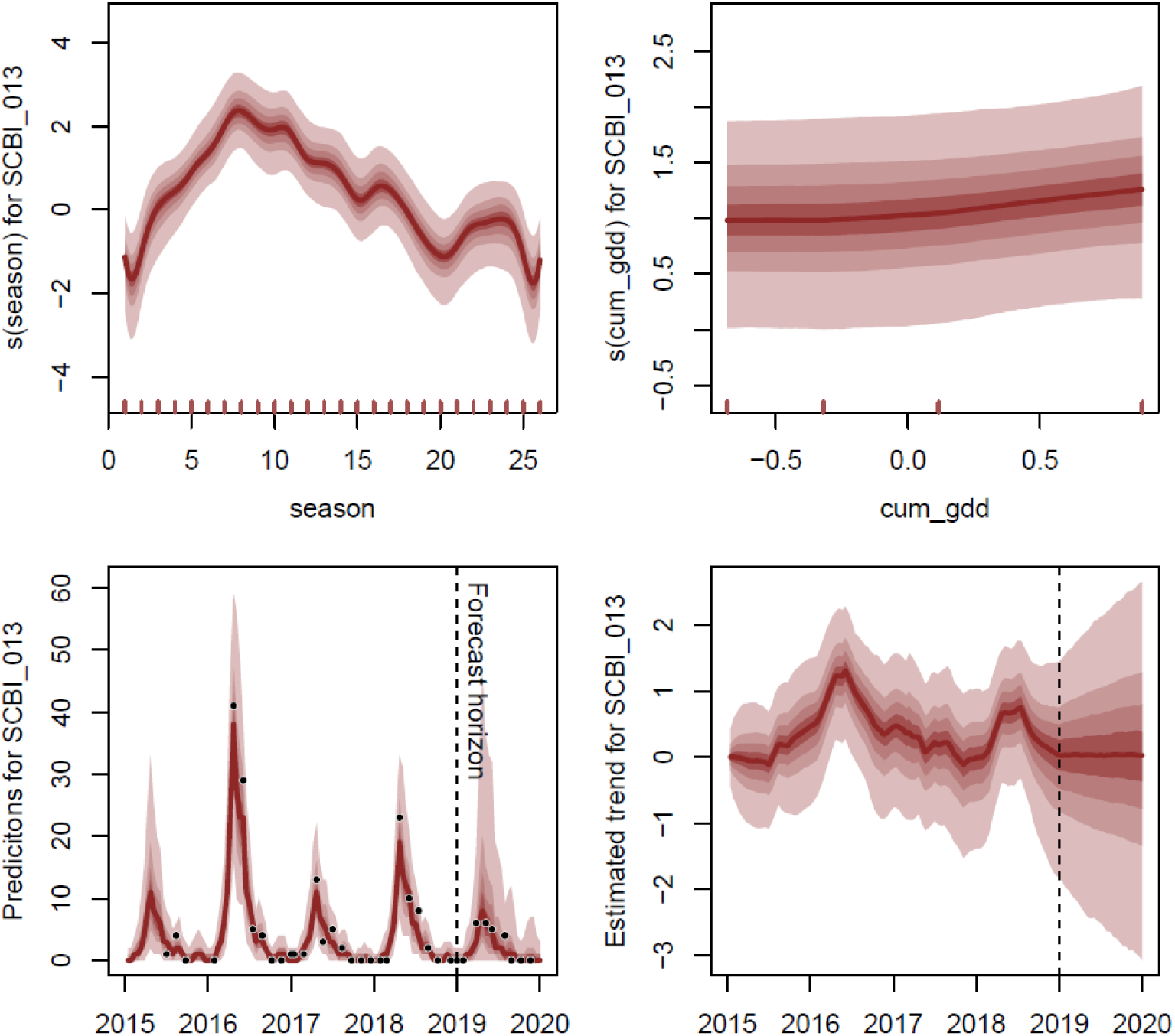
Visualisations from the best-performing *mvgam* model (Hyp3) for a single *Ixodes scapularis* plot (SCBI_013). Top left, the estimated seasonal smooth function (the seasonal time window refers to calendar weeks 15 – 40); top right, estimated cumulative growing degree days function; bottom left, predicted tick abundances over time (observed values shown as black points); bottom right, estimated latent dynamic component. For all plots shading shows posterior empirical quantiles. Hypothesis definitions are outlined in section **CASE STUDY: FORECASTING TICK ABUNDANCES**.

In agreement with the *I. scapularis* models, *A. americanum* abundance was also best predicted by the Hyp3 model. Example visualisations of estimated plot-level seasonal functions are shown in Figure 6. Our model estimated that tick abundances in some plots (i.e. SERC_001) tend to show earlier peaks followed by rapid declines, while abundance in other plots (i.e. UKFS_003) follow a broader curve with a less obvious peak (Figure 6).

**Figure 6:**
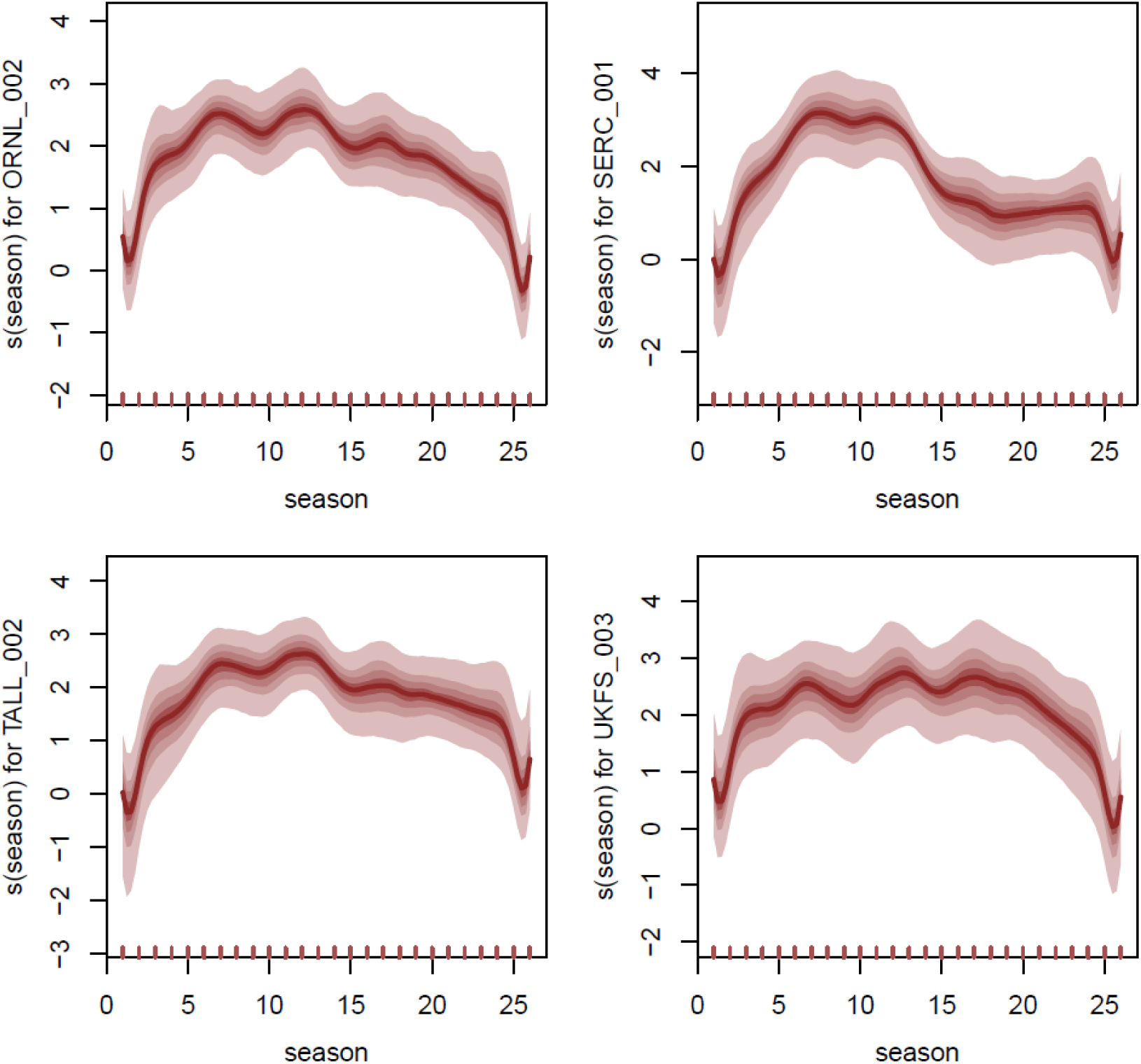
Output from the *plot_mvgam_smooth* function in *mvgam* showing seasonal smooth functions for four *Amblyomma americanum* plots estimated from a dynamic GAM with hierarchical seasonality. Shading shows posterior empirical quantiles. Note that seasonal indices 1 – 26 correspond to epidemiological weeks 15 – 41.

### Quantifying uncertainty contributions among *mvgam* model components

In addition to plotting smooth functions and forecasts, *mvgam* also offers utilities to compute relative contributions of the latent dynamic and GAM components to forecast uncertainty. This process of partitioning uncertainty into different components is an important step in analysing a model’s forecasts to diagnose the main drivers of prediction uncertainty and prioritise aspects of models or data that require further investigation (Dietze 2017, Heilman et al. 2022). Comparisons of uncertainty contributions for four of the *A. americanum* forecasts indicate that both components contribute to forecast uncertainty, but to varying degrees over time and across plots (Figure 7). However, across all plots, dynamic trend uncertainty tended to increase over time, becoming relatively more important during the peak tick season (3 – 22 weeks ahead).

**Figure 7:**
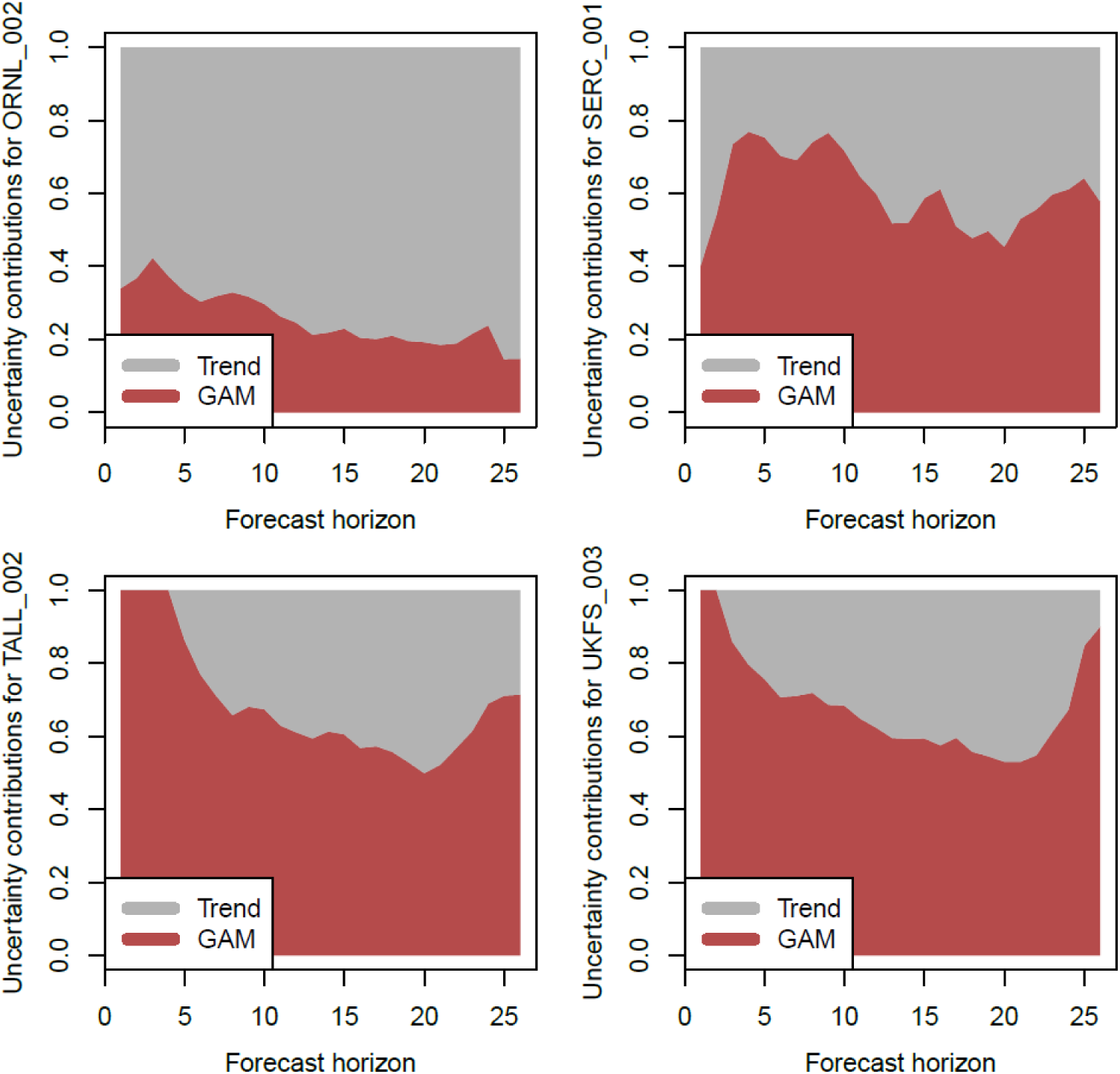
Output from the *plot_mvgam_uncertainty* function in *mvgam* showing relative contributions of the dynamic temporal (grey) and GAM (red) components to forecast uncertainty for four *Amblyomma americanum* plots estimated from a dynamic GAM with hierarchical seasonality. Forecast horizons were varied over a ‘one-year’ horizon (26 weeks matching data availability, excluding winter).

## DISCUSSION

We have introduced an R package for fitting Bayesian Dynamic GAMs (DGAM) that incorporate the flexibility of the widely popular penalised smoothing functions in *mgcv* with latent dynamic components for time series analysis and forecasting. Keys to *mvgam*’s performance are its ability to cope with the substantial missing data, scale to large sets of discrete time series and provide robust uncertainty quantification. In recent years there has been increased interest in using models for uncertainty interval estimation as opposed to point predictions, a trend that lends well to Bayesian learning (Gelman et al. 2017, Makridakis et al. 2020). This is particularly relevant for ecological forecasts, where point estimates are less important for making informed decisions than are conditional probability statements (Dietze 2017, Dietze et al. 2018, White et al. 2019).

Notably, JAGS model files and all data necessary to condition the model are made available to the user in *mvgam*, allowing an enormous diversity of bespoke models to be implemented through addition of other stochastic or hierarchical elements. The case studies (available in Appendices S1 – S3 and online at https://rpubs.com/NickClark47/mvgam, https://rpubs.com/NickClark47/mvgam2, https://rpubs.com/NickClark47/mvgam3) discuss a range of models that can be fitted and interrogated with *mvgam*, however we see several avenues for improving model flexibility and estimation. These include but are not limited to:

- Implementations in the STAN probabilistic programming language, which uses Hamiltonian Monte Carlo to more efficiently sample from complex posteriors as well as more in-depth diagnostics to identify problems with posterior estimation (Carpenter et al. 2017); this may also reduce biases in inference and speed up computation for data sets with complicated posteriors
- Recommending and accommodating ways for users to include principled prior specifications for the shapes of nonlinear smooth functions (Simpson et al. 2017)
- The addition of Markov-switching processes to allow dynamic factor loadings to be drawn from different sets of correlation ‘regimes’, allowing correlation structures to change over time in a principled way (Fox et al. 2010)
- The incorporation of covariates into the latent factor model (i.e. as dynamic linear models) to explicitly address broader hypotheses about the factors that influence temporal dynamics (Heilman et al. 2022)

## CONCLUSION

The R package *mvgam* provides a user-friendly tool for researchers and practitioners interested in fitting GAMs to analyse and forecast ecological time series. The problems associated with smooth extrapolation are not limited to ecology however, as the need to forecast sets of discrete nonlinear time series is a common challenge in areas as diverse as speech recognition, tourism demand, natural language processing and finance (Hyndman and Athanasopoulos 2018, Makridakis et al. 2018). Beyond the examples showcased here, the package can be especially useful to identify avenues for model improvement via its ability to assimilate new observations online to update forecast distributions (showcased in detail in Appendix S1 and in the *mvgam* package’s vignettes). With growing interest in both the application of hierarchical GAMs to ecological problems and the need to use iterative forecasts to make ecology a more predictive discipline, *mvgam* can become a vital addition to the applied ecologist’s analytical toolbox.

## ACKNOWLEDGEMENTS

NOAA temperature data was supplied by Daniel Ruiz-Carrascal as part of the 2021 NEON Ecological Forecasting Challenge.

## SUPPLEMENTARY MATERIALS

Appendix S1: Univariate dynamic GAMs, model comparison and data assimilation using the *mvgam* package for R

Appendix S2: Multivariate dynamic GAMs using the *mvgam* package for R

Appendix S3: Distributed lag dynamic GAMs using the *mvgam* package for R

Appendix S4: Supplementary figures

