## Appendix S1 for "Dynamic Generalised Additive Models (DGAM) for forecasting discrete ecological time series"

**APPENDIX S1.** Univariate dynamic GAMs, model comparison and data assimilation using the mvgam package for R

Nicholas Clark and Konstans Wells

Generalised Additive Models (GAMs) are incredibly flexible tools that have found particular application in the analysis of time series. In ecology, a host of recent papers and workshops (i.e. the [2018 Ecological Society of America workshop on GAMs](https://noamross.github.io/mgcv-esa-2018/) that was hosted by Eric Pedersen, David L. Miller, Gavin Simpson, and Noam Ross) have drawn special attention to the wide range of applications that GAMs have for addressing complex ecological problems. Given the many ways that GAMs can model temporal data, it is tempting to use the smooth functions estimated by a GAM to produce out of sample forecasts. Here we will inspect the behaviours of smoothing splines when extrapolating to data outside the range of the training data to examine whether this can be useful in practice.

We will work in the mvgam package, which fits dynamic GAMs (DGAMs) using MCMC sampling via the JAGS software (Note that JAGS 4.3.0 is required; installation links are found [here](https://sourceforge.net/projects/mcmc-jags/files/)). Briefly, assume ${\overset{̃}{\mathbf{y}}}_{t}$ is the conditional expectation of a discrete response variable $\mathbf{y}$ at time $\mathbf{t}$. Asssuming $\mathbf{y}$ is drawn from a distribution within the exponential family of distributions (such as Poisson or Negative Binomial). Using a log link function, the linear predictor for a dynamic GAM is written as:

$$log({\overset{̃}{\mathbf{y}}}_{t})=\beta_{0}+\sum_{i=1}^{I} \mathbf{s}_{i,t}(\mathbf{x}_{i,t})+\mathbf{z}_{t} ,$$

Here $\beta_{0}$ is the unknown intercept, the $\mathbf{s}$'s are unknown smooth functions of the covariates ($\mathbf{x}$'s) and $\mathbf{z}$ is a dynamic latent trend component. Each smooth function $\mathbf{s}_{i}$ is composed of spline like basis expansions whose coefficients, which must be estimated, control the shape of the functional relationship between $\mathbf{x}_{i}$ and $log(\overset{̃}{\mathbf{y}})$. The size of the basis expansion limits the smooth’s potential complexity, with a larger set of basis functions allowing greater flexibility. Several advantages of GAMs are that they can model a diversity of response families, including discrete distributions (i.e. Poisson or Negative Binomial) that accommodate ecological features such as zero-inflation, and that they can be formulated to include hierarchical smoothing for multivariate responses. For the dynamic component, in its most basic form we assume a random walk with drift:

$$\mathbf{z}_{t}=\phi+\mathbf{z}_{t-1}+\mathbf{e}_{t} ,$$

where $\phi$ is an optional drift parameter (if the latent trend is assumed to not be stationary) and $\mathbf{e}$ is drawn from a zero-centred Gaussian distribution. This model is easily modified to include autoregressive terms, which mvgam accommodates up to order = 3.

### Why DGAMs?

Dynamic GAMs are useful when we wish to predict future values from time series that show temporal dependence and we want to avoid extrapolating from a smooth (which, as you’ll see below, can sometimes lead to unpredictable and unrealistic behaviours). In addition, smooths can often try to wiggle excessively to capture any autocorrelation that is present in a time series, which exacerbates the problem of forecasting ahead. Here we use an exaggerated example to show how a smooth tries to wiggle excessively, leading to overfitting by lessening the smoothing penalty $\lambda$. Notice how wiggly the function becomes in the last plot when $\lambda$ is very small, which is what mgcv’s gam.check function will often encourage you to do if you are trying to model an autocorrelated time series with a smooth function

```
# Fit a model to the mcycle dataset in
# the MASS package, fixing the smoothing
# parameter at an abnormally large value
par(mfrow = c(2, 2), mgp = c(2.5, 1, 0),
    mai = c(0.6, 0.6, 0.2, 0.2))
library(mgcv)
## Loading required package: nlme
## This is mgcv 1.8-33. For overview type 'help("mgcv-package")'.
data(mcycle, package = "MASS")
m1 <- gam(accel ~ s(times, k = 50), data = mcycle,
    method = "REML", sp = 5)

plot(m1, scheme = 1, residuals = TRUE, pch = 16,
    ylab = expression(lambda == 5), xlab = "",
    shade.col = scales::alpha("#B97C7C",
        0.6))

# Fit models with increasingly relaxed
# smooths (lambdas approaching zero)
m2 <- gam(accel ~ s(times, k = 50), data = mcycle,
    method = "REML", sp = 1)
plot(m2, scheme = 1, residuals = TRUE, pch = 16,
    ylab = expression(lambda == 1), xlab = "",
    shade.col = scales::alpha("#B97C7C",
        0.6))

m3 <- gam(accel ~ s(times, k = 50), data = mcycle,
    method = "REML", sp = 0.01)
plot(m3, scheme = 1, residuals = TRUE, pch = 16,
    ylab = expression(lambda == 0.01), xlab = "",
    shade.col = scales::alpha("#B97C7C",
        0.6))

m4 <- gam(accel ~ s(times, k = 50), data = mcycle,
    method = "REML", sp = 1e-07)
plot(m4, scheme = 1, residuals = TRUE, pch = 16,
    ylab = expression(lambda == 1e-07), xlab = "",
    shade.col = scales::alpha("#B97C7C",
        0.6))
```

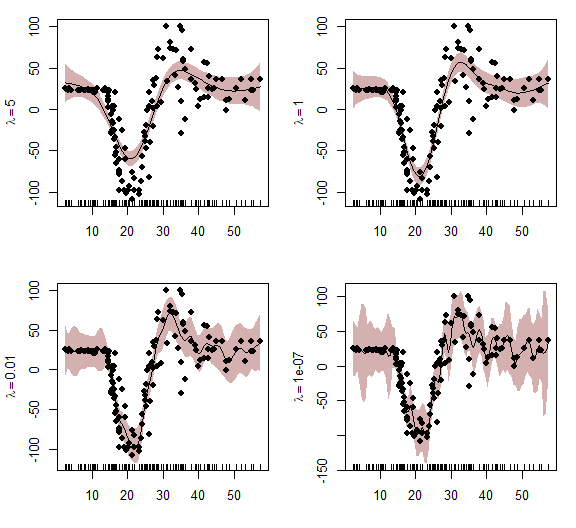

### AirPassengers example

Now onto an empirical example. First we load the AirPassengers data from the forecast package and convert to an xts object. This series is a good starting point as it should be highly forecastable given its stable seasonal pattern and nearly linear trend

```
# devtools::install_github('nicholasjclark/mvgam')
library(mvgam)
library(dplyr)
library(xts)
library(forecast)
data("AirPassengers")
series <- xts::as.xts(floor(AirPassengers))
colnames(series) <- c("Air")
```

View the raw series, its STL decomposition and the distribution of observations. There is a clear seasonal pattern as well as an increasing trend over time for this series, and the distribution shows evidence of a skew suggestive of overdispersion

plot(series)

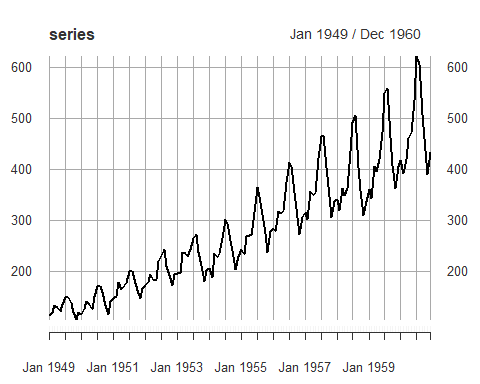

plot(stl(series, s.window = "periodic"))

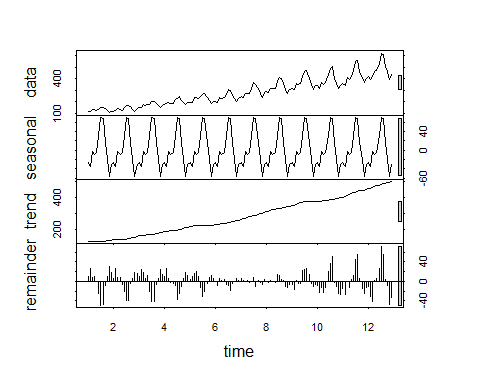

hist(series)

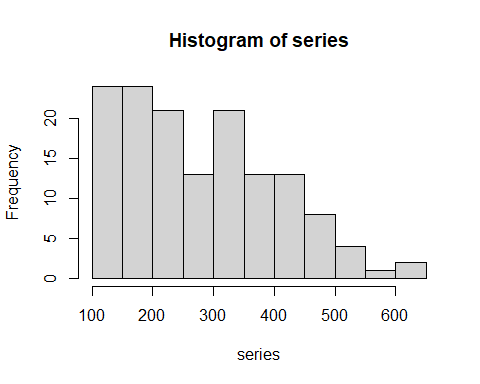

Next use the series_to_mvgam function, which converts ts or xts objects into the correct format for mvgam. Here we set train_prop = 0.75, meaning we will include ~ 75% of the observations in the training set and use the remaining ~25% for forecast validation. We also randomly set ~10% of observations to NA so that we can evaluate models based on their predictive abilities

```
series[sample(1:length(series), floor(length(series)/10),
    F)] <- NA
fake_data <- series_to_mvgam(series, freq = 12,
    train_prop = 0.75)
```

Examine the returned object

```
head(fake_data$data_train)
##     y season year       date series
## 1 112      1 1949 1949-01-01    Air
## 2 118      2 1949 1949-02-01    Air
## 3 132      3 1949 1949-03-01    Air
## 4 129      4 1949 1949-04-01    Air
## 5 121      5 1949 1949-05-01    Air
## 6 135      6 1949 1949-06-01    Air
head(fake_data$data_test)
##     y season year       date series
## 1 340      1 1958 1958-01-01    Air
## 2 318      2 1958 1958-02-01    Air
## 3 362      3 1958 1958-03-01    Air
## 4 348      4 1958 1958-04-01    Air
## 5 363      5 1958 1958-05-01    Air
## 6 435      6 1958 1958-06-01    Air
```

As a first pass at modelling this series, we will fit a GAM that includes a seasonal smooth and a yearly smooth, as well as their tensor product interaction. As the seasonality is cyclic, we will use a cyclic cubic regression spline for season. The knots are set so that the boundaries of the cyclic smooth match up between December 31 and January 1. We will stick with the default thin plate regression spline for year. This is similar to what we might do when fitting a model in mgcv to try and forecast ahead, except here we also have an explicit model for the residual component. mvgam uses the jagam function from the mgcv package to generate a skeleton JAGS file and updates that file to incorporate any dynamic trend components (so far limited to no trend, random walks or AR models up to order 3). This is advantageous as any GAM that is allowed in mgcv can in principle be used in mvgam to fit dynamic linear models with smooth functions for nonlinear covariate effects. For multivariate series, mvgam also includes an option for dimension reduction by inducing trend correlations via a dynamic factor process. Here we use the Negative Binomial family and a burnin length of 10000 iterations (in practice we would probably run these models for a bit longer, but they tend to converge rather quickly for univariate series thanks to the useful starting values supplied by jagam). Note that feeding the data_test object does not mean that these values are used in training of the model. Rather, they are included as NA so that we can automatically create a forecast from the posterior predictions for these observations. This is useful for plotting forecasts without needing to run new data through the model's equations later on

```
mod1 <- mvjagam(data_train = fake_data$data_train,
    data_test = fake_data$data_test, formula = y ~
        s(season, bs = c("cc"), k = 12) +
            s(year, k = 5) + ti(season, year,
            bs = c("cc", "tp"), k = c(12,
                3)), knots = list(season = c(0.5,
        12.5)), family = "nb", trend_model = "None",
    burnin = 10000, chains = 4)
## Compiling rjags model...
## Starting 4 rjags simulations using a PSOCK cluster with 4 nodes on host
## 'localhost'
## Simulation complete
## Note: Summary statistics were not produced as there are >50 monitored
## variables
## [To override this behaviour see ?add.summary and ?runjags.options]
## FALSEFinished running the simulation
## NOTE: Stopping adaptation
```

We can view the JAGS model file that has been generated to see the additions that have been made to the base gam model. If the user selects return_jags_data = TRUE when calling mvjagam, this file can be modified and the resulting jags_data object can also be modified to fit more complex Bayesian models. Note that here the AR and phi terms have been set to zero as this model does not include a dynamic trend component

```
mod1$model_file
##  [1] "model {"
##  [2] ""
##  [3] "## GAM linear predictor"
##  [4] "eta <- X %*% b"
##  [5] ""
##  [6] "## Mean expectations"
##  [7] "for (i in 1:n) {"
##  [8] " for (s in 1:n_series) {"
##  [9] "  mu[i, s] <- exp(eta[ytimes[i, s]] + trend[i, s])"
## [10] " }"
## [11] "}"
## [12] ""
## [13] ""
## [14] "## State space trend estimates"
## [15] "for(s in 1:n_series) {"
## [16] " trend[1, s] <- 0"
## [17] "}"
## [18] ""
## [19] "for(s in 1:n_series) {"
## [20] " trend[2, s] <- 0"
## [21] "}"
## [22] ""
## [23] "for(s in 1:n_series) {"
## [24] " trend[3, s] <- 0"
## [25] "}"
## [26] ""
## [27] "for (i in 4:n){"
## [28] " for (s in 1:n_series){"
## [29] " trend[i, s] <- 0"
## [30] " }"
## [31] "}"
## [32] ""
## [33] "## AR components"
## [34] "for (s in 1:n_series){"
## [35] " phi[s] <- 0"
## [36] " ar1[s] <- 0"
## [37] " ar2[s] <- 0"
## [38] " ar3[s] <- 0"
## [39] " tau[s] <- pow(sigma[s], -2)"
## [40] " sigma[s] ~ dexp(1)"
## [41] "}"
## [42] ""
## [43] "## Negative binomial likelihood functions"
## [44] "for (i in 1:n) {"
## [45] " for (s in 1:n_series) {"
## [46] "  y[i, s] ~ dnegbin(rate[i, s], r[s])"
## [47] "  rate[i, s] <- ifelse((r[s] / (r[s] + mu[i, s])) < min_eps, min_eps,"
## [48] "                      (r[s] / (r[s] + mu[i, s])))"
## [49] " }"
## [50] "}"
## [51] ""
## [52] "## Complexity penalising prior for the overdispersion parameter;"
## [53] "## where the likelihood reduces to a 'base' model (Poisson) unless"
## [54] "## the data support overdispersion"
## [55] "for(s in 1:n_series){"
## [56] " r[s] <- pow(r_raw[s], 2)"
## [57] " r_raw[s] ~ dexp(0.05)"
## [58] "}"
## [59] ""
## [60] ""
## [61] "## Posterior predictions"
## [62] "for (i in 1:n) {"
## [63] " for (s in 1:n_series) {"
## [64] "  ypred[i, s] ~ dnegbin(rate[i, s], r[s])"
## [65] " }"
## [66] "}"
## [67] ""
## [68] "  ## parametric effect priors (regularised for identifiability)"
## [69] "  for (i in 1:1) { b[i] ~ dnorm(p_coefs[i], p_taus[i]) }"
## [70] "  ## prior for s(season)... "
## [71] "  K1 <- S1[1:10,1:10] * lambda[1] "
## [72] "  b[2:11] ~ dmnorm(zero[2:11],K1) "
## [73] "  ## prior for s(year)... "
## [74] "  K2 <- S2[1:4,1:4] * lambda[2]  + S2[1:4,5:8] * lambda[3]"
## [75] "  b[12:15] ~ dmnorm(zero[12:15],K2) "
## [76] "  ## prior for ti(season,year)... "
## [77] "  K3 <- S3[1:20,1:20] * lambda[4]  + S3[1:20,21:40] * lambda[5]"
## [78] "  b[16:35] ~ dmnorm(zero[16:35],K3) "
## [79] "   ## smoothing parameter priors..."
## [80] "  for (i in 1:5) {"
## [81] "   lambda[i] ~ dexp(0.05)"
## [82] "    rho[i] <- log(lambda[i])"
## [83] "  }"
## [84] "}"
```

Inspect the summary from the model, which is somewhat similar to an mgcv model summary with extra information about convergences for unobserved parameters. The estimated degrees of freedom, smooth coefficients and smooth penalties are all extracted from the mvgam model using sim2jam so that approximate p-values can be calculated using Nychka's method (following Wood (2013) Biometrika 100(1), 221-228). Note however that this function is still in development and approximate p-values may not be entirely accurate

```
summary_mvgam(mod1)
## GAM formula:
## y ~ s(season, bs = c("cc"), k = 12) + s(year, k = 5) + ti(season,
##     year, bs = c("cc", "tp"), k = c(12, 3))
##
## Family:
## Negative Binomial
##
## N series:
## 1
##
## N observations per series:
## 108
##
## Dispersion parameter estimates:
##       2.5%      50%    97.5% Rhat n.eff
## r 1420.145 4808.771 19184.22    1  6931
##
## GAM smooth term approximate significances:
##                    edf Ref.df     F p-value
## s(season)        9.305 10.000 71.42  <2e-16 ***
## s(year)          2.970  4.000  1.82  0.0287 *
## ti(season,year) 13.535 20.000  0.45  0.7952
## ---
## Signif. codes:  0 '***' 0.001 '**' 0.01 '*' 0.05 '.' 0.1 ' ' 1
##
## GAM coefficient (beta) estimates:
##                             2.5%          50%       97.5% Rhat n.eff
## (Intercept)         5.3617039992  5.376516980  5.39100477 1.00  5005
## s(season).1        -0.3006532423 -0.203569750 -0.12224345 1.01   284
## s(season).2        -0.1553232053 -0.099797055 -0.04339400 1.01   410
## s(season).3        -0.0552180878  0.005721609  0.05768055 1.00   539
## s(season).4        -0.0716619601 -0.024460344  0.01923282 1.00   747
## s(season).5         0.0188869604  0.068254691  0.11684041 1.01   622
## s(season).6         0.1768893549  0.224929231  0.27131354 1.01   710
## s(season).7         0.1596064357  0.201204849  0.24338185 1.01   839
## s(season).8         0.0007097294  0.061207925  0.11915350 1.00   484
## s(season).9        -0.1931972470 -0.146302259 -0.09464402 1.01   632
## s(season).10       -0.2115236870 -0.123212416 -0.05026308 1.01   340
## s(year).1          -0.0855502842 -0.020821478  0.03566510 1.01   481
## s(year).2          -0.1153729369  0.001419933  0.10324581 1.01   301
## s(year).3          -0.1090373704  0.027093060  0.15022744 1.01   308
## s(year).4           0.3105853801  0.363200343  0.42137657 1.01   506
## ti(season,year).1  -0.0749586922  0.026768006  0.12458633 1.02   392
## ti(season,year).2  -0.1211240353 -0.029759616  0.05939996 1.01   409
## ti(season,year).3  -0.0576971299  0.039840579  0.13759629 1.01   365
## ti(season,year).4  -0.1251970869 -0.045613898  0.04189770 1.01   363
## ti(season,year).5  -0.0702068611  0.021127890  0.11390787 1.01   397
## ti(season,year).6  -0.0981392413 -0.018255370  0.05907532 1.01   473
## ti(season,year).7  -0.1105918490 -0.016483763  0.07449801 1.01   381
## ti(season,year).8  -0.0680242420  0.003584732  0.07226105 1.01   531
## ti(season,year).9  -0.1248294598 -0.033953797  0.05863891 1.00   453
## ti(season,year).10 -0.0363352179  0.038911103  0.10690080 1.00   457
## ti(season,year).11 -0.1308357927 -0.041640839  0.04068933 1.02   414
## ti(season,year).12 -0.0230462381  0.052272935  0.12604668 1.01   481
## ti(season,year).13 -0.1137691498 -0.028905971  0.06281482 1.01   428
## ti(season,year).14 -0.0278869326  0.039097675  0.11078923 1.01   520
## ti(season,year).15 -0.0864095286  0.012060225  0.10972359 1.01   381
## ti(season,year).16 -0.0823113971 -0.002223676  0.07425475 1.01   469
## ti(season,year).17 -0.0907145258  0.001427760  0.09118494 1.00   424
## ti(season,year).18 -0.0852133237 -0.011091242  0.06480228 1.01   496
## ti(season,year).19 -0.0809634915  0.018492850  0.11392207 1.02   398
## ti(season,year).20 -0.1138041323 -0.025137706  0.06146877 1.02   431
##
## GAM smoothing parameter (rho) estimates:
##                         2.5%      50%    97.5% Rhat n.eff
## s(season)         3.30366159 4.360038 5.181157    1  1564
## s(year)           1.71998758 3.415267 4.615338    1  2683
## s(year)2         -0.03415296 2.333975 3.731829    1  5991
## ti(season,year)   3.56971134 4.575152 5.322749    1  3010
## ti(season,year)2  2.54233652 3.920714 4.863448    1  2344
##
## Latent trend drift (phi) and AR parameter estimates:
##     2.5% 50% 97.5% Rhat n.eff
## phi    0   0     0  NaN     0
## ar1    0   0     0  NaN     0
## ar2    0   0     0  NaN     0
## ar3    0   0     0  NaN     0
##
```

mvgam takes a fitted gam model and adapts the model file to fit in JAGS, with possible extensions to deal with stochastic trend components and other features. But as this model has not been modified much from the original gam model with the same formula (i.e. we have not added any stochastic trend components to the linear predictor yet, which is the main feature of mvgam), the summary of the unmodified gam model is also useful and fairly accurate

```
summary(mod1$mgcv_model)
##
## Family: Negative Binomial(1235039572.565)
## Link function: log
##
## Formula:
## y ~ s(season, bs = c("cc"), k = 12) + s(year, k = 5) + ti(season,
##     year, bs = c("cc", "tp"), k = c(12, 3))
##
## Parametric coefficients:
##             Estimate Std. Error t value Pr(>|t|)
## (Intercept) 5.384894   0.007066   762.1   <2e-16 ***
## ---
## Signif. codes:  0 '***' 0.001 '**' 0.01 '*' 0.05 '.' 0.1 ' ' 1
##
## Approximate significance of smooth terms:
##                   edf Ref.df        F p-value
## s(season)       7.754 10.000   33.250  <2e-16 ***
## s(year)         1.404  1.694 1498.721  <2e-16 ***
## ti(season,year) 1.518 20.000    0.202  0.0569 .
## ---
## Signif. codes:  0 '***' 0.001 '**' 0.01 '*' 0.05 '.' 0.1 ' ' 1
##
## R-sq.(adj) =  0.989   Deviance explained = 98.9%
## fREML = 135.15  Scale est. = 1         n = 98
```

Ordinarily we would be quite pleased with this result, as we have explained most of the variation in the series with a fairly simple model. We can plot the estimated smooth functions and with their associated credible intervals, which are interpreted similarly to mgcv plots except they are not centred at zero (mvgam predicts the smooth with all other covariates at their means, rather than at zero). So the plot below, for the season smooth, shows the estimated values for the linear predictor (on the log scale in this case) if year was set to its mean

plot_mvgam_smooth(mod1, series = 1, smooth = "season")

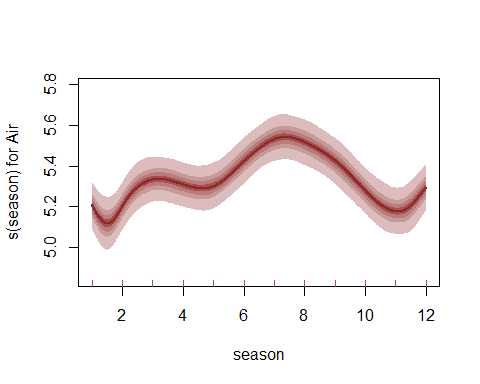

And here is the plot of the smooth function for year, which has essentially estimated a straight line

plot_mvgam_smooth(mod1, series = 1, smooth = "year")

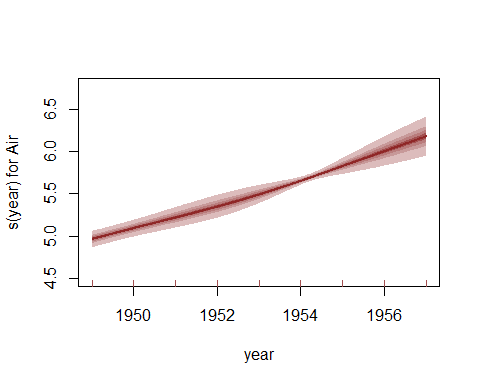

Perform a series of posterior predictive checks to see if the model is able to simulate data for the training period that looks realistic and unbiased. First, examine simulated kernel densities for posterior predictions (yhat) and compare to the density of the observations (y)

```
plot_mvgam_ppc(mod1, series = 1, type = "density",
    legend_position = "bottomright")
```

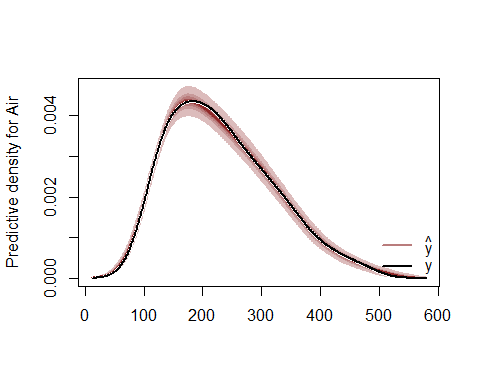

Now plot the distribution of predicted means compared to the observed mean

plot_mvgam_ppc(mod1, series = 1, type = "mean")

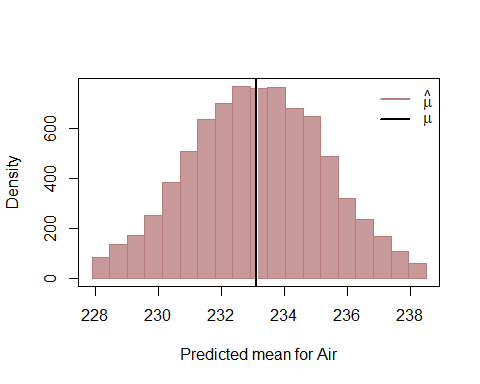

Next examine simulated empirical Cumulative Distribution Functions (CDF) for posterior predictions (yhat) and compare to the CDF of the observations (y)

plot_mvgam_ppc(mod1, series = 1, type = "cdf")

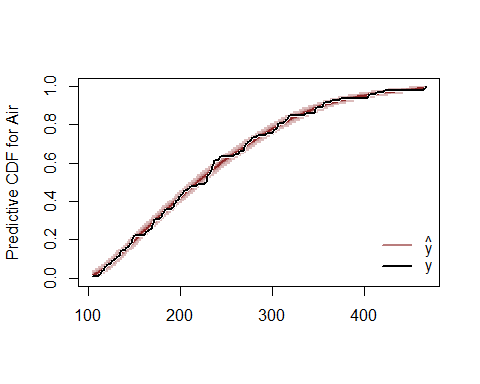

Finally look for any biases in predictions by examining a Probability Integral Transform (PIT) histogram. If our predictions are not biased one way or another (i.e. not consistently under- or over-predicting), this histogram should look roughly uniform

plot_mvgam_ppc(mod1, series = 1, type = "pit")

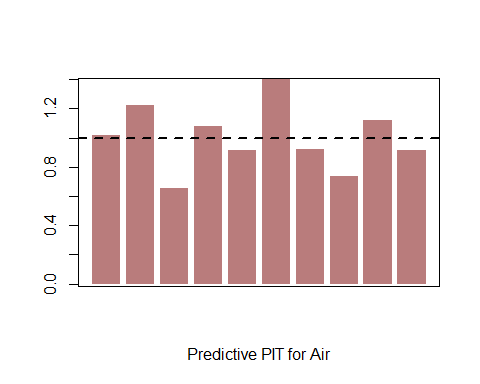

All of these plots indicate the model is well calibrated against the training data, with no apparent pathological behaviors exhibited. Now for some investigation of the estimated relationships and forecasts. We can also perform residual diagnostics using randomised quantile (Dunn-Smyth) residuals. These look reasonable overall, though there is some autocorrelation left in the residuals for this series

plot_mvgam_resids(mod1, series = 1)

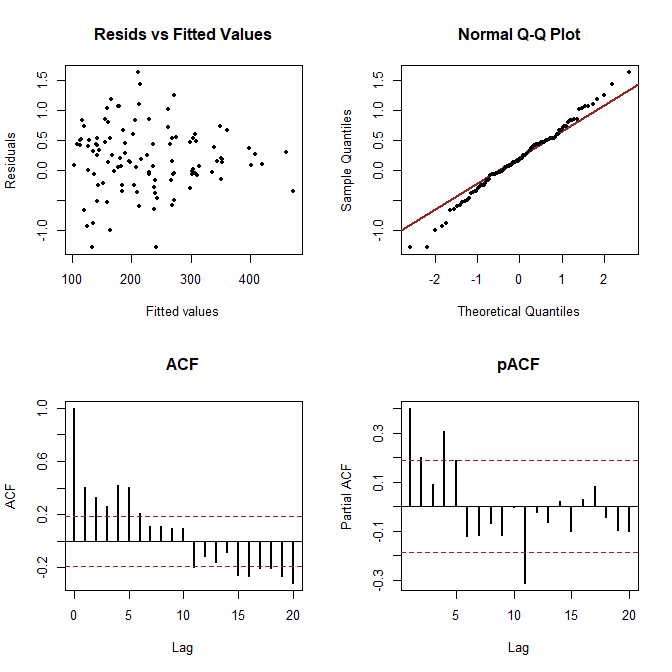

Ok so the model is doing well when fitting against the training data, but how are its forecasts? The yearly trend is being extrapolated into the future, which controls most of the shape and uncertainty in the forecast. We see there is a reasonable estimate of uncertainty and the out of sample observations (to the right of the dashed line) are all within the model's 95% HPD intervals

plot_mvgam_fc(mod1, series = 1, data_test = fake_data$data_test)

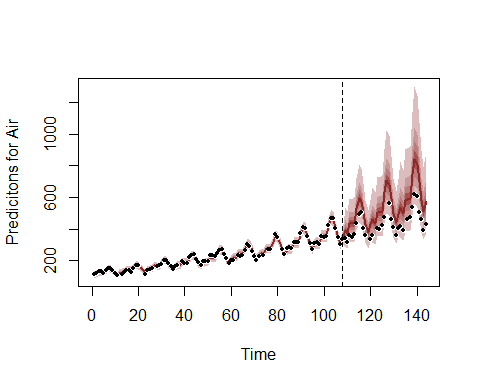

The extrapolation of the smooth for year can be better viewed by feeding new data to the plot_mvgam_smooth function. Here we feed values of year to cover the training and testing set to see how the extrapolation would continue into the future. The dashed line marks the end of the training period

```
plot_mvgam_smooth(mod1, series = 1, "year",
    newdata = expand.grid(year = seq(min(fake_data$data_train$year),
        max(fake_data$data_test$year), length.out = 500),
        season = mean(fake_data$data_train$season),
        series = unique(fake_data$data_train$series)))
abline(v = max(fake_data$data_train$year),
    lty = "dashed")
```

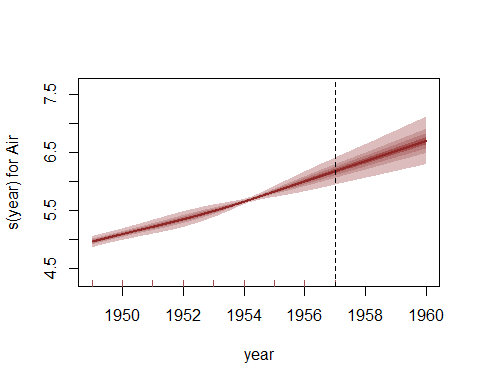

We can also re-do the posterior predictive checks, but this time focusing only on the out of sample period. This will give us better insight into how the model is performing and whether it is able to simulate realistic and unbiased future values

```
plot_mvgam_ppc(mod1, series = 1, type = "density",
    data_test = fake_data$data_test, legend_position = "bottomright")
```

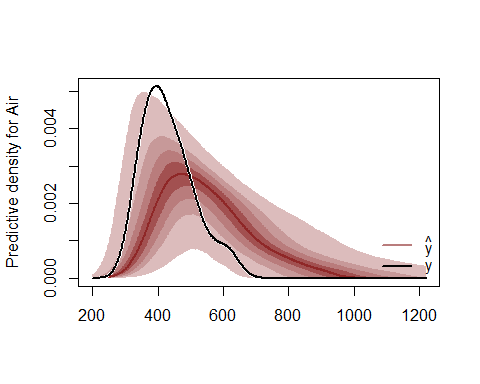

```
plot_mvgam_ppc(mod1, series = 1, type = "mean",
    data_test = fake_data$data_test)
```

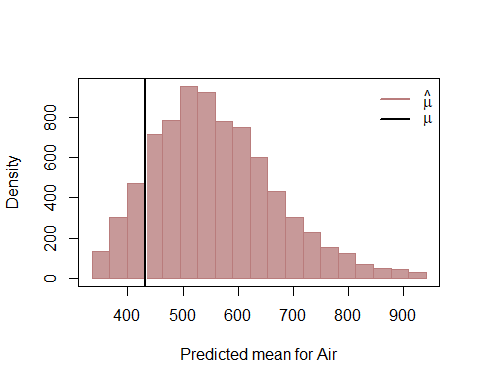

```
plot_mvgam_ppc(mod1, series = 1, type = "cdf",
    data_test = fake_data$data_test)
```

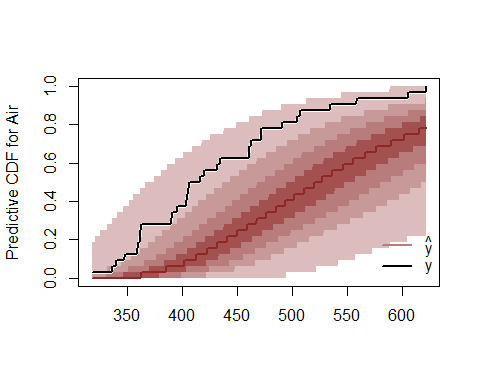

```
plot_mvgam_ppc(mod1, series = 1, type = "pit",
    data_test = fake_data$data_test)
```

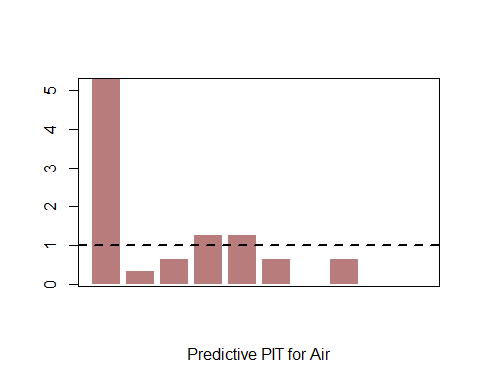

There are some problems with the way this model is generating future predictions. Perhaps a different smooth function for year can help? Here we fit our original model again but use a different smooth term for year to try and capture the long-term trend (using B splines with multiple penalties, following the excellent example by Gavin Simpson about [extrapolating with smooth terms](https://fromthebottomoftheheap.net/2020/06/03/extrapolating-with-gams/)). This is similar to what we might do when trying to forecast ahead from a more wiggly function, as B splines have useful properties by allowing the penalty to be extended into the range of values we wish to predict (in this case, the years in data_test). Note that we drop the year*season tensor product interaction to help emphasize the extrapolation behaviour of the B spline

```
mod2 <- mvjagam(data_train = fake_data$data_train,
    data_test = fake_data$data_test, formula = y ~
        s(season, bs = c("cc"), k = 12) +
            s(year, bs = "bs", m = c(2, 1)),
    knots = list(season = c(0.5, 12.5), year = c(min(fake_data$data_train$year) -
        1, min(fake_data$data_train$year),
        max(fake_data$data_train$year), max(fake_data$data_test$year))),
    family = "nb", trend_model = "None",
    burnin = 10000, chains = 4)
## Compiling rjags model...
## Starting 4 rjags simulations using a PSOCK cluster with 4 nodes on host
## 'localhost'
## Simulation complete
## Note: Summary statistics were not produced as there are >50 monitored
## variables
## [To override this behaviour see ?add.summary and ?runjags.options]
## FALSEFinished running the simulation
## NOTE: Stopping adaptation
```

Again as we haven't modified the base gam much, the summary from the mgcv model is fairly accurate

```
summary_mvgam(mod2)
## GAM formula:
## y ~ s(season, bs = c("cc"), k = 12) + s(year, bs = "bs", m = c(2,
##     1))
##
## Family:
## Negative Binomial
##
## N series:
## 1
##
## N observations per series:
## 108
##
## Dispersion parameter estimates:
##       2.5%      50%    97.5% Rhat n.eff
## r 1485.573 5028.517 19800.27    1  6869
##
## GAM smooth term approximate significances:
##              edf Ref.df      F p-value
## s(season)  7.922 10.000  42.38  <2e-16 ***
## s(year)    6.708  9.000 472.31  <2e-16 ***
## ---
## Signif. codes:  0 '***' 0.001 '**' 0.01 '*' 0.05 '.' 0.1 ' ' 1
##
## GAM coefficient (beta) estimates:
##                     2.5%         50%        97.5% Rhat n.eff
## (Intercept)   5.36008604  5.37475618  5.389383163 1.00  5427
## s(season).1  -0.22266004 -0.15399652 -0.100807220 1.00   511
## s(season).2  -0.14103290 -0.09592817 -0.050298269 1.00   707
## s(season).3  -0.03161467  0.01395403  0.057867194 1.00   725
## s(season).4  -0.05212099 -0.01005199  0.030555165 1.01   795
## s(season).5   0.04126523  0.08463668  0.126382792 1.01   819
## s(season).6   0.20631996  0.24470179  0.283691356 1.00   937
## s(season).7   0.18107956  0.21760027  0.254943176 1.00   971
## s(season).8   0.02057670  0.06537401  0.109687492 1.01   786
## s(season).9  -0.16580342 -0.12127261 -0.077331617 1.00   672
## s(season).10 -0.15112490 -0.10259435 -0.053397573 1.01   723
## s(year).1    -1.44045225 -0.53117655  0.177054758 1.05   120
## s(year).2    -0.45955139 -0.34572213 -0.227631173 1.00   464
## s(year).3    -0.14673112 -0.07351254 -0.005736511 1.00  1007
## s(year).4     0.08306412  0.15862606  0.232464873 1.00   738
## s(year).5     0.17731921  0.24740738  0.318807391 1.01   836
## s(year).6     0.37008889  0.44379555  0.518091036 1.00   787
## s(year).7     0.60411254  0.67751265  0.751618949 1.01   548
## s(year).8     0.53544611  0.70157355  0.859392737 1.01   351
## s(year).9    -0.16496156  0.54361975  1.336530243 1.06   151
##
## GAM smoothing parameter (rho) estimates:
##               2.5%      50%    97.5% Rhat n.eff
## s(season) 5.220748 6.466272 7.507741    1  1038
## s(year)   2.991927 4.101116 4.917211    1  1218
##
## Latent trend drift (phi) and AR parameter estimates:
##     2.5% 50% 97.5% Rhat n.eff
## phi    0   0     0  NaN     0
## ar1    0   0     0  NaN     0
## ar2    0   0     0  NaN     0
## ar3    0   0     0  NaN     0
##
summary(mod2$mgcv_model)
##
## Family: Negative Binomial(323170648.711)
## Link function: log
##
## Formula:
## y ~ s(season, bs = c("cc"), k = 12) + s(year, bs = "bs", m = c(2,
##     1))
##
## Parametric coefficients:
##             Estimate Std. Error t value Pr(>|t|)
## (Intercept) 5.384149   0.007072   761.3   <2e-16 ***
## ---
## Signif. codes:  0 '***' 0.001 '**' 0.01 '*' 0.05 '.' 0.1 ' ' 1
##
## Approximate significance of smooth terms:
##             edf Ref.df      F p-value
## s(season) 7.720     10  36.87  <2e-16 ***
## s(year)   6.573      7 365.24  <2e-16 ***
## ---
## Signif. codes:  0 '***' 0.001 '**' 0.01 '*' 0.05 '.' 0.1 ' ' 1
##
## R-sq.(adj) =  0.989   Deviance explained =   99%
## fREML = 142.55  Scale est. = 1         n = 98
```

This model explains even more of the variation than the thin plate yearly model above, so we'd be tempted to use it for prediction (though of course we'd want to perform a series of checks following advice from Simon Wood; see [lectures by Gavin Simpson for more information on how to perform these checks](https://www.youtube.com/user/ucfagls)). So how do the forecasts look?

plot_mvgam_fc(mod2, series = 1, data_test = fake_data$data_test)

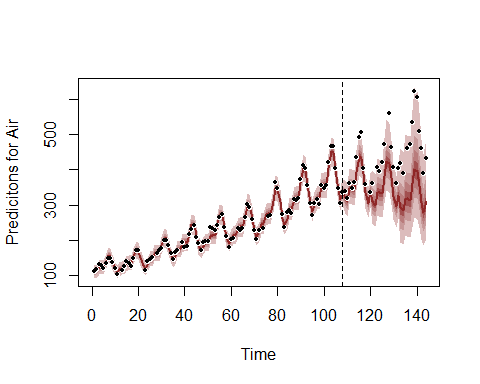

Again the forecast is being driven primarily by the extrapolation behaviour of the B spline. Look at this behaviour for the year smooth as we did above by feeding new data for prediction

```
plot_mvgam_smooth(mod2, series = 1, "year",
    newdata = expand.grid(year = seq(min(fake_data$data_train$year),
        max(fake_data$data_test$year), length.out = 500),
        season = mean(fake_data$data_train$season),
        series = unique(fake_data$data_train$series)))
abline(v = max(fake_data$data_train$year),
    lty = "dashed")
```

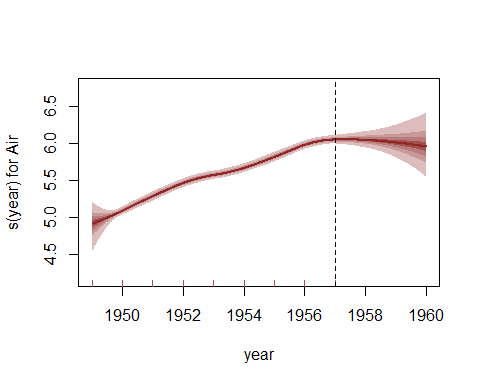

Residual and posterior in-training predictive checks for this model will look similar to the above, with some autocorrelation left in residuals but nothing terribly alarming. But the forecast checks again show some problems:

```
plot_mvgam_ppc(mod2, series = 1, type = "density",
    data_test = fake_data$data_test)
```

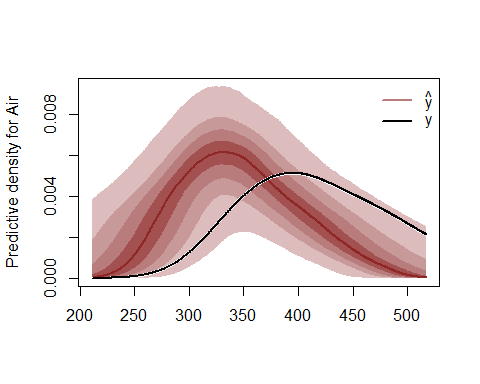

```
plot_mvgam_ppc(mod2, series = 1, type = "mean",
    data_test = fake_data$data_test)
```

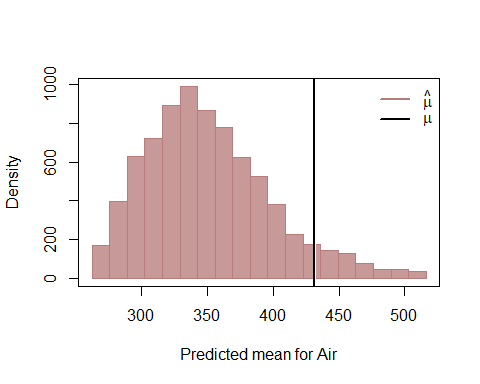

```
plot_mvgam_ppc(mod2, series = 1, type = "cdf",
    data_test = fake_data$data_test)
```

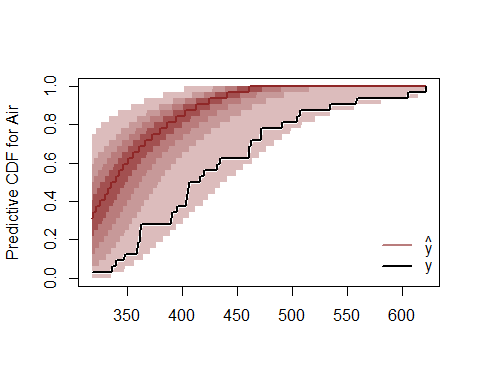

```
plot_mvgam_ppc(mod2, series = 1, type = "pit",
    data_test = fake_data$data_test)
```

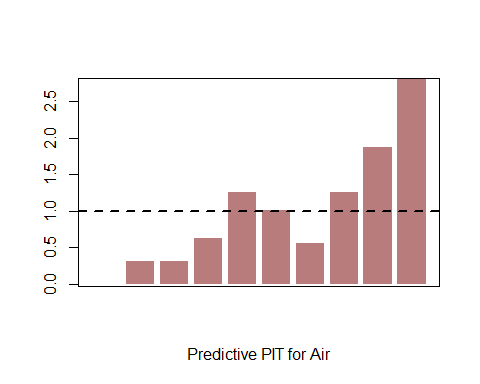

Can we improve the forecasts by removing our reliance on extrapolation? Now we will fit a model in which the GAM component of the linear predictor captures the repeated seasonality (again with a cyclic smooth) and a dynamic latent trend captures the residual process using AR parameters (up to order 3). This model is a mildly modified version of the base mgcv model where the linear predictor is augmented with the latent trend component. Slightly longer burnin is used here due to the added complexity of the time series component, but the model still fits in ~ 30 seconds on most machines

```
mod3 <- mvjagam(data_train = fake_data$data_train,
    data_test = fake_data$data_test, formula = y ~
        s(season, bs = c("cc"), k = 12),
    knots = list(season = c(0.5, 12.5)),
    family = "nb", trend_model = "AR3", drift = TRUE,
    burnin = 10000, chains = 4)
## Compiling rjags model...
## Starting 4 rjags simulations using a PSOCK cluster with 4 nodes on host
## 'localhost'
## Simulation complete
## Note: Summary statistics were not produced as there are >50 monitored
## variables
## [To override this behaviour see ?add.summary and ?runjags.options]
## FALSEFinished running the simulation
## NOTE: Stopping adaptation
```

In this case the fitted model is more different to the base mgcv model that was used in jagam to produce the skeleton JAGS file, so the summary of that base model is less accurate. But we can still check the model summary for the mvjagam mdoel to examine convergence for key parameters

```
summary_mvgam(mod3)
## GAM formula:
## y ~ s(season, bs = c("cc"), k = 12)
##
## Family:
## Negative Binomial
##
## N series:
## 1
##
## N observations per series:
## 108
##
## Dispersion parameter estimates:
##       2.5%      50%    97.5% Rhat n.eff
## r 1457.319 4909.302 20093.45    1  6820
##
## GAM smooth term approximate significances:
##              edf Ref.df     F p-value
## s(season)  6.627 10.000 28.71  <2e-16 ***
## ---
## Signif. codes:  0 '***' 0.001 '**' 0.01 '*' 0.05 '.' 0.1 ' ' 1
##
## GAM coefficient (beta) estimates:
##                      2.5%         50%        97.5% Rhat n.eff
## (Intercept)   4.713415371  4.76645030  4.842108788 1.02    17
## s(season).1  -0.127167526 -0.07727746 -0.035023213 1.01   559
## s(season).2  -0.077745831 -0.03385860  0.008845667 1.00   601
## s(season).3   0.003655087  0.04279386  0.083387717 1.01   780
## s(season).4  -0.027788756  0.01592638  0.057512453 1.00   656
## s(season).5   0.056972712  0.09839778  0.137721410 1.01   666
## s(season).6   0.195668168  0.23477662  0.272101494 1.00   818
## s(season).7   0.167061412  0.20228189  0.237402135 1.00   849
## s(season).8  -0.004628294  0.03834552  0.080722616 1.00   721
## s(season).9  -0.209217621 -0.16329473 -0.118736011 1.00   635
## s(season).10 -0.209095121 -0.16272187 -0.121171869 1.00   684
##
## GAM smoothing parameter (rho) estimates:
##               2.5%      50%    97.5% Rhat n.eff
## s(season) 5.912033 7.189075 8.226405    1  1140
##
## Latent trend drift (phi) and AR parameter estimates:
##            2.5%        50%      97.5% Rhat n.eff
## phi  0.01252322 0.02363859 0.03492748 1.01   352
## ar1 -0.12695748 0.32221560 0.76128027 1.00  1099
## ar2 -0.12303751 0.31941747 0.74306221 1.00  1469
## ar3 -0.08587543 0.36182508 0.78239429 1.01   823
##
```

The seasonal term is obviously still very important. Plot it here

plot_mvgam_smooth(mod3, series = 1, "season")

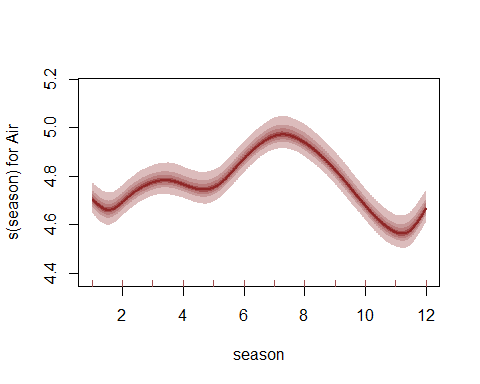

Plot diagnostics of posterior median Dunn-Smyth residuals, where the autocorrelation is now captured by the latent trend process

plot_mvgam_resids(mod3, series = 1)

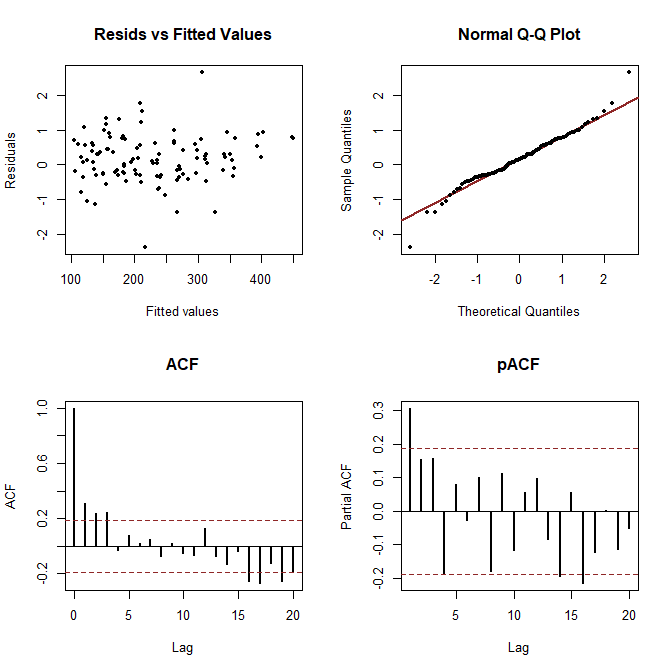

How does the model's posterior forecast distribution compare to the previous models?

plot_mvgam_fc(mod3, series = 1, data_test = fake_data$data_test)

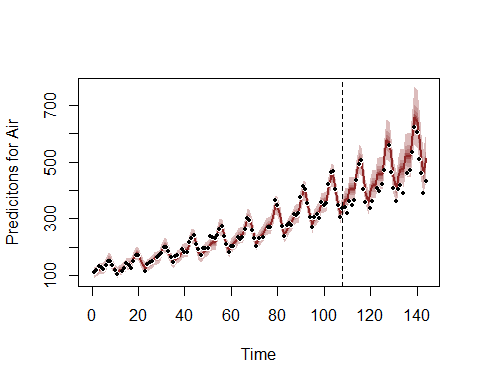

```
plot_mvgam_ppc(mod3, series = 1, type = "density",
    data_test = fake_data$data_test)
```

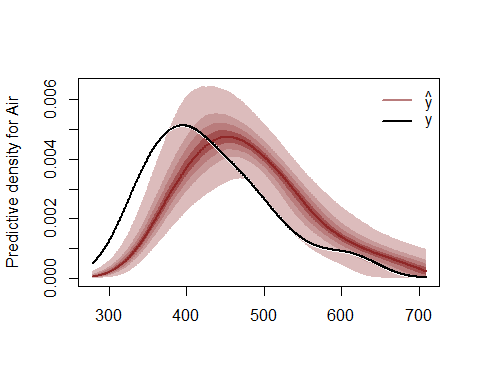

```
plot_mvgam_ppc(mod3, series = 1, type = "mean",
    data_test = fake_data$data_test)
```

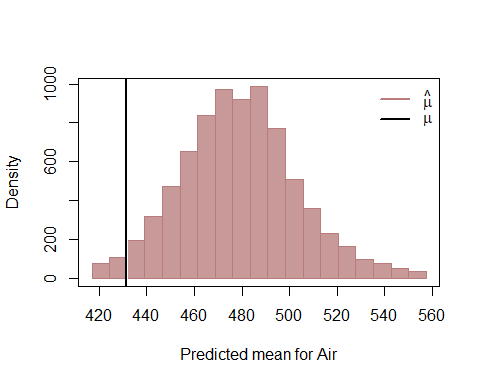

```
plot_mvgam_ppc(mod3, series = 1, type = "cdf",
    data_test = fake_data$data_test)
```

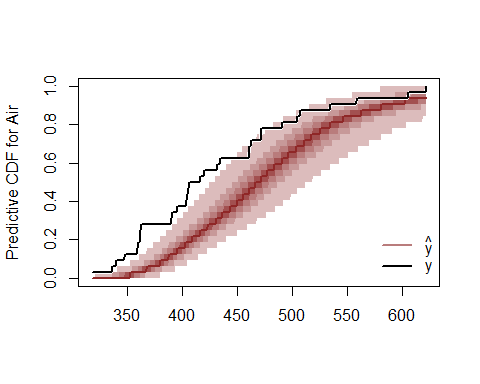

```
plot_mvgam_ppc(mod3, series = 1, type = "pit",
    data_test = fake_data$data_test)
```

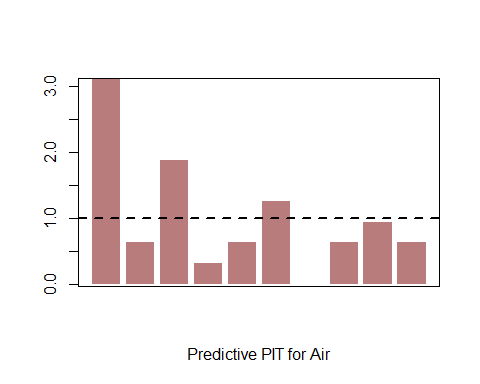

None of the model's is a perfect representation of the data generating process, but the forecast for our dynamic GAM is more stable and more accurate than the previous models, with the added advantage that we can place more trust our estimated smooth for season because we have captured the residual autocorrelation. The posterior checks for our dynamic GAM also look much better than the two previous models. Plot the estimated latent dynamic trend (which is on the log scale)

plot_mvgam_trend(mod3, series = 1, data_test = fake_data$data_test)

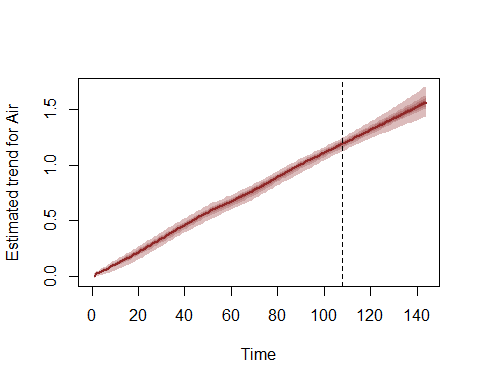

Benchmarking against "null" models is a very important part of evaluating a proposed forecast model. After all, if our complex dynamic model can't generate better predictions then a random walk or mean forecast, is it really telling us anything new about the data-generating process? Here we examine the model comparison utilities in mvgam. Here we illustrate how this can be done in mvgam by fitting a simpler model by smoothing on a white noise covariate rather than on the seasonal variable. Because the white noise covariate is not informative and we are using a random walk for the trend process, this model essentially becomes a Poisson observation model over a dynamic random walk process.

```
fake_data$data_train$fake_cov <- rnorm(NROW(fake_data$data_train))
fake_data$data_test$fake_cov <- rnorm(NROW(fake_data$data_test))
mod4 <- mvjagam(data_train = fake_data$data_train,
    data_test = fake_data$data_test, formula = y ~
        s(fake_cov, k = 3), family = "poisson",
    trend_model = "RW", drift = TRUE, burnin = 10000,
    chains = 4)
## Compiling rjags model...
## Starting 4 rjags simulations using a PSOCK cluster with 4 nodes on host
## 'localhost'
## Simulation complete
## Note: Summary statistics were not produced as there are >50 monitored
## variables
## [To override this behaviour see ?add.summary and ?runjags.options]
## FALSEFinished running the simulation
## NOTE: Stopping adaptation
```

Look at this model's proposed forecast

plot_mvgam_fc(mod4, series = 1, data_test = fake_data$data_test)

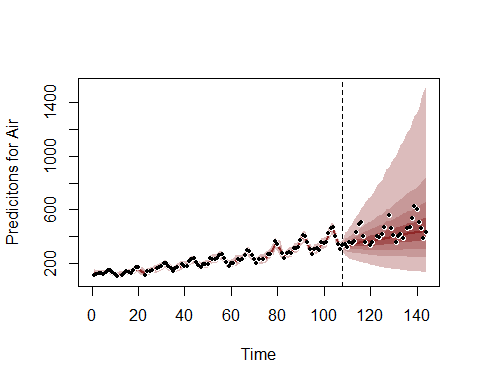

Here we showcase how different dynamic models can be compared using rolling probabilistic forecast evaluation, which is especially useful if we don't already have out of sample observations for comparing forecasts. This function sets up a sequence of evaluation timepoints along a rolling window within the training data to evaluate 'out-of-sample' forecasts. The trends are rolled forward a total of fc_horizon timesteps according to their estimated state space dynamics to generate an 'out-of-sample' forecast that is evaluated against the true observations in the horizon window. We are therefore simulating a situation where the model's parameters had already been estimated but we have only observed data up to the evaluation timepoint and would like to generate forecasts that consider the possible future paths for the latent trends and the true observed values for any other covariates in the horizon window. Evaluation involves calculating the Discrete Rank Probability Score and a binary indicator for whether or not the true value lies within the forecast's 90% prediction interval. For this test we compare the two models on the exact same sequence of 30 evaluation points using horizon = 6

```
compare_mvgams(model1 = mod3, model2 = mod4,
    fc_horizon = 6, n_evaluations = 30, n_cores = 3)
## DRPS summaries per model (lower is better)
##             Min.   1st Qu.    Median      Mean  3rd Qu.     Max. NA's
## Model 1 2.645059  3.939968  4.942927  5.693311  5.94467 16.31657   18
## Model 2 6.430293 13.804852 21.154800 25.844825 30.67976 95.72758   18
##
## 90% interval coverages per model (closer to 0.9 is better)
## Model 1 1
## Model 2 1
```

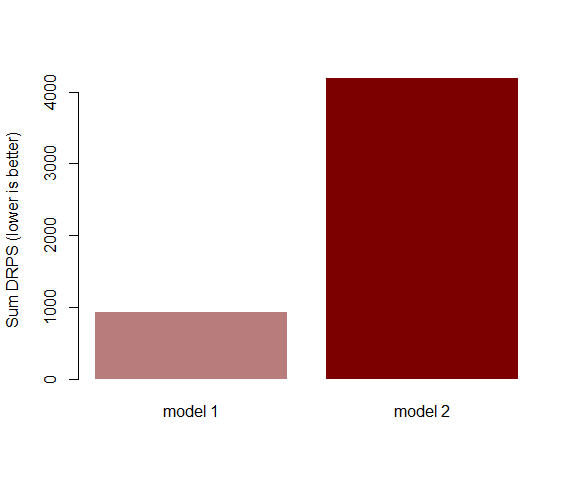

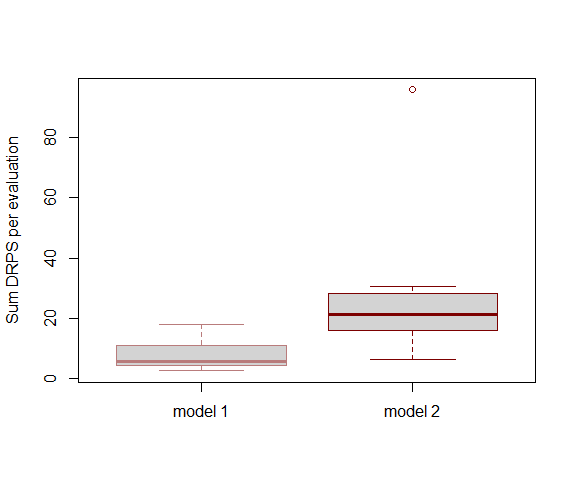

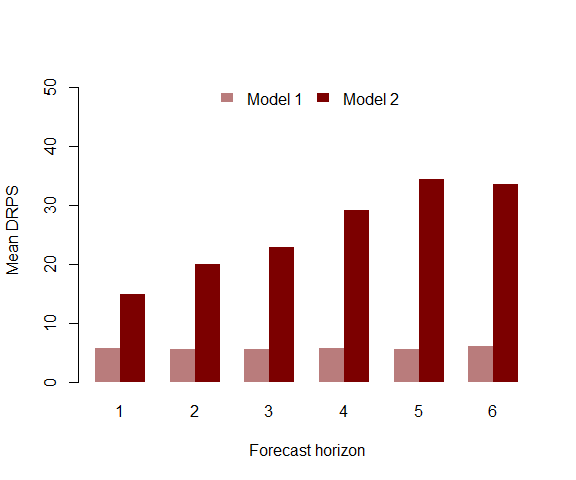

The series of plots generated by compare_mvgams clearly show that the first dynamic model generates better predictions. In each plot, DRPS for the forecast horizon is lower for the first model than for the second model. This kind of evaluation is often more appropriate for forecast models than complexity-penalising fit metrics such as [AIC or BIC](https://www.sciencedirect.com/science/article/pii/S0169207020301096) However, comparing forecasts of the dynamic models against the two models with yearly smooth terms (mod1 and mod2) using the rolling window approach is actually not recommended, as the yearly smooth models have already seen all the possible in-sample values of year and so should be able to predict incredibly well by interpolating through the range of the fitted smooth. By contrast, the dynamic component in our dynamic GAMs (models 3 and 4) produce true forecasts when running the rolling window approach. Nevertheless, when we compare some of these models (here mod1 with the thin plate yearly smooth vs mod3 with the AR3 trend process) as we did above for the random walk model, we still find that our dynamic GAM produces superior probabilistic forecasts

```
compare_mvgams(model1 = mod1, model2 = mod3,
    fc_horizon = 6, n_evaluations = 30, n_cores = 3)
## DRPS summaries per model (lower is better)
##              Min.   1st Qu.    Median     Mean   3rd Qu.     Max. NA's
## Model 1 19.570158 29.497692 37.949647 40.02268 46.794927 74.09630   18
## Model 2  2.760663  3.947724  4.918484  5.69030  5.725024 16.13515   18
##
## 90% interval coverages per model (closer to 0.9 is better)
## Model 1 1
## Model 2 1
```

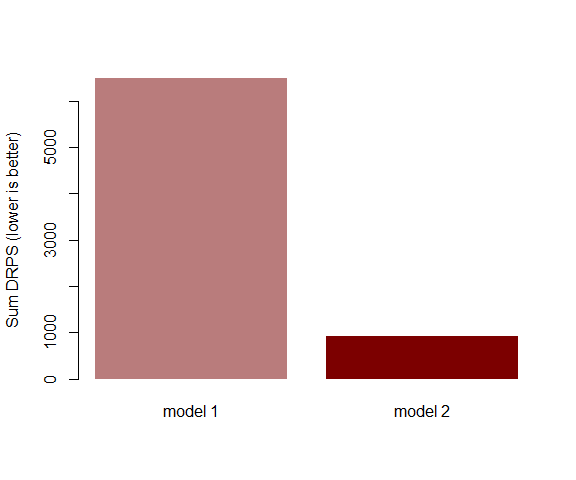

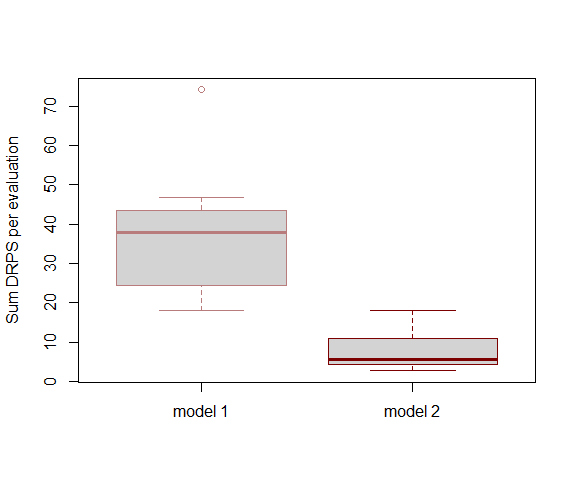

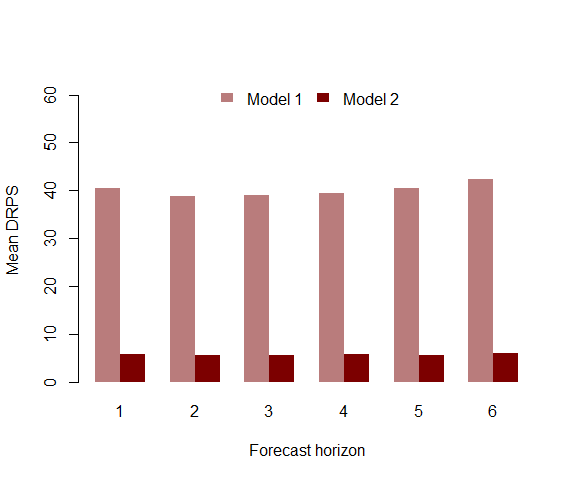

The same holds true when comparing against the B spline model (mod2)

```
compare_mvgams(model1 = mod2, model2 = mod3,
    fc_horizon = 6, n_evaluations = 30, n_cores = 3)
## DRPS summaries per model (lower is better)
##              Min.   1st Qu.    Median      Mean   3rd Qu.     Max. NA's
## Model 1 20.595417 30.046477 38.299026 39.017945 48.338757 68.45039   18
## Model 2  2.511966  3.905224  4.913061  5.704745  5.942385 16.34070   18
##
## 90% interval coverages per model (closer to 0.9 is better)
## Model 1 1
## Model 2 1
```

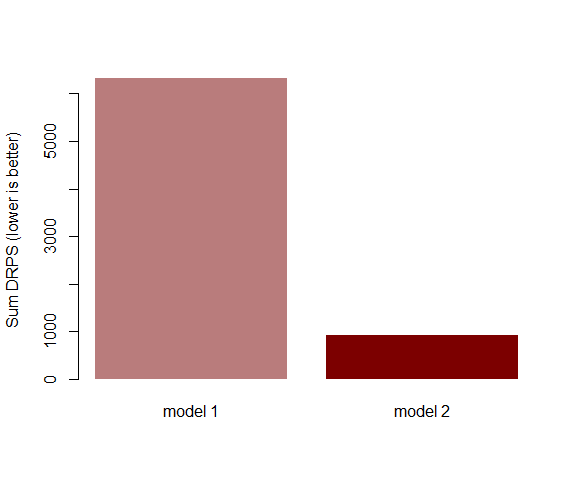

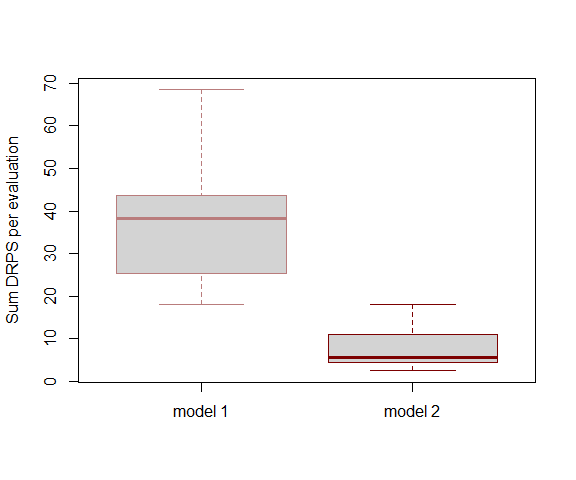

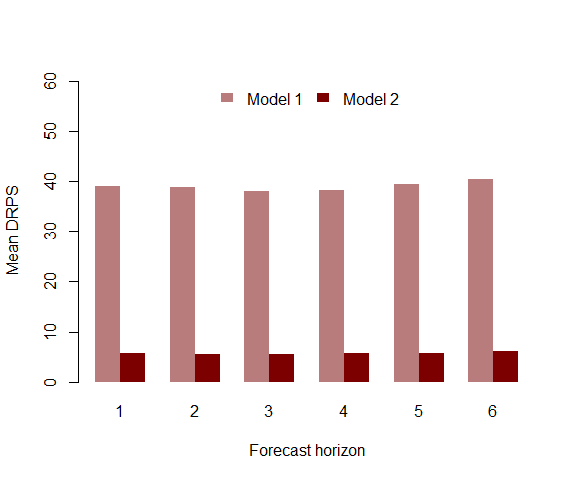

Now we proceed by exploring how forecast distributions from an mvgam object can be automatically updated in light of new incoming observations. This works by generating a set of "particles" that each captures a unique proposal about the current state of the system (in this case, the current estimate of the latent trend component). The next observation in data_assim is assimilated and particles are weighted by how well their proposal (i.e. their proposed forecast, prior to seeing the new data) matched the new observations. For univariate models such as the ones we've fitted so far, this weight is represented by the proposal's Negative Binomial log-likelihood. For multivariate models, a multivariate composite likelihood is used for weights. Once weights are calculated, we use importance sampling to update the model's forecast distribution for the remaining forecast horizon. Begin by initiating a set of 15000 particles by assimilating the next observation in data_test and storing the particles in the default location (in a directory called particles within the working directory)

```
pfilter_mvgam_init(object = mod3, n_particles = 15000,
    n_cores = 3, data_assim = fake_data$data_test)
## Saving particles to pfilter/particles.rda
##  ESS = 14999.64
```

Now we are ready to run the particle filter. This function will assimilate the next six out of sample observations in data_test and update the forecast after each assimilation step. This works in an iterative fashion by calculating each particle's weight, then using a kernel smoothing algorithm to "pull" low weight particles toward the high-likelihood space before assimilating the next observation. The strength of the kernel smoother is controlled by kernel_lambda, which in our experience works well when left to the default of 1. If the Effective Sample Size of particles drops too low, suggesting we are putting most of our belief in a very small set of particles, an automatic resampling step is triggered to increase particle diversity and reduce the chance that our forecast intervals become too narrow and incapable of adapting to changing conditions

```
pfilter_mvgam_online(data_assim = fake_data$data_test[1:7,
    ], n_cores = 3, kernel_lambda = 1)
## Particles have already assimilated one or more observations. Skipping these
##
## Assimilating the next 6 observations
##
## Effective sample size is 14999.91 ...
##
## Smoothing particles ...
##
## Effective sample size is 14999.68 ...
##
## Smoothing particles ...
##
## Effective sample size is 14999.89 ...
##
## Smoothing particles ...
##
## Effective sample size is 14999.55 ...
##
## Smoothing particles ...
##
## Effective sample size is 14999.54 ...
##
## Smoothing particles ...
##
## Effective sample size is 14999.63 ...
##
## Smoothing particles ...
##
## Last assimilation time was 7 1958
##
## Saving particles to pfilter/particles.rda
##  ESS = 15000
```

Once assimilation is complete, generate the updated forecast from the particles using the covariate information in remaining data_test observations. This function is designed to hopefully make it simpler to assimilate observations, as all that needs to be provided once the particles are initiated as a dataframe of test data in exactly the same format as the data that were used to train the initial model. If no new observations are found (observations are arranged by year and then by season so the consistent indexing of these two variables is very important!) then the function returns a NULL and the particles remain where they are in state space.

```
fc <- pfilter_mvgam_fc(file_path = "pfilter",
    n_cores = 3, data_test = fake_data$data_test,
    ylim = c(min(fake_data$data_train$y,
        na.rm = T), max(fake_data$data_test$y,
        na.rm = T) * 1.25))
```

Compare the updated forecast to the original forecast to see how it has changed in light of the most recent observations

```
par(mfrow = c(1, 2))
plot_mvgam_fc(mod3, series = 1, data_test = fake_data$data_test,
    ylim = c(min(fake_data$data_train$y,
        na.rm = T), max(fake_data$data_test$y,
        na.rm = T) * 1.25))
fc$Air()
```

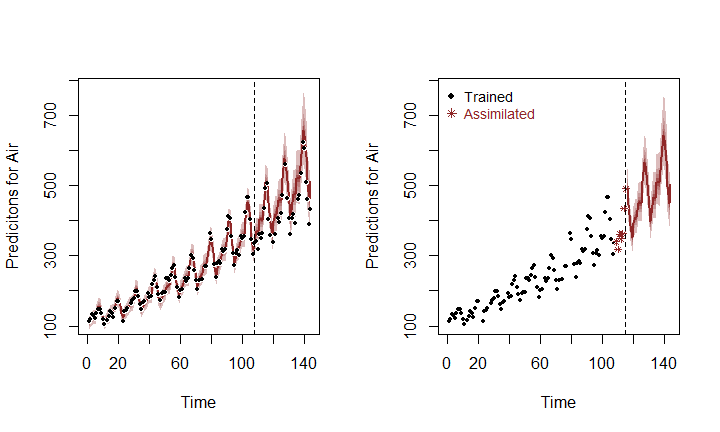

Here it is apparent that the distribution has shifted slightly in light of the 6 observations that have been assimilated, and that our confidence in the remaining forecast horizon has improved (tighter uncertainty intervals). This is an advantageous way of allowing a model to slowly adapt to new conditions while breaking free of restrictive assumptions about residual distributions. [See some of the many particle filtering lectures by Nathaniel Osgood for more details](https://www.youtube.com/user/NathanielOsgood). Remove the particles from their stored directory when finished

unlink("pfilter", recursive = T)

### Lynx example

For our next univariate example, we will again pursue how challenging it can be to forecast ahead with conventional GAMs and how mvgam overcomes these challenges. We begin by replicating the lynx analysis from [2018 Ecological Society of America workshop on GAMs](https://noamross.github.io/mgcv-esa-2018/) that was hosted by Eric Pedersen, David L. Miller, Gavin Simpson, and Noam Ross, with some minor adjustments. First, load the data and plot the series as well as its estimated autocorrelation function

```
library(mvgam)
data(lynx)
lynx_full = data.frame(year = 1821:1934,
    population = as.numeric(lynx))
plot(lynx_full$population, type = "l", ylab = "Lynx trappings",
    xlab = "Time")
acf(lynx_full$population, main = "")
```

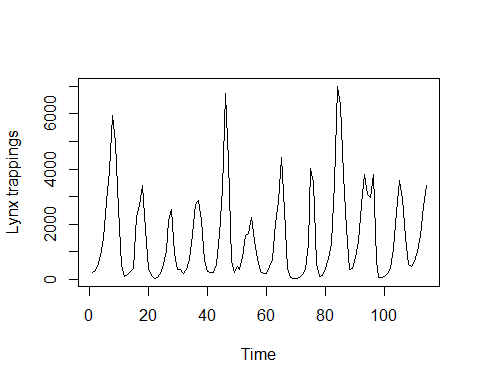

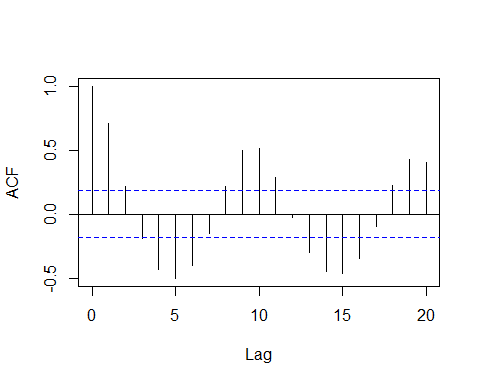

There is a clear ~19-year cyclic pattern to the data, so I create a season term that can be used to model this effect and give a better representation of the data generating process. I also create a new year term that represents which long-term cycle each observation is in

```
plot(stl(ts(lynx_full$population, frequency = 19),
    s.window = "periodic"))
```

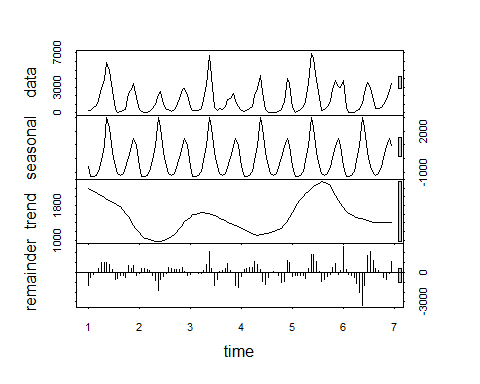

```
lynx_full$season <- (lynx_full$year%%19) +
    1
cycle_ends <- c(which(lynx_full$season ==
    19), NROW(lynx_full))
cycle_starts <- c(1, cycle_ends[1:length(which(lynx_full$season ==
    19))] + 1)
cycle <- vector(length = NROW(lynx_full))
for (i in 1:length(cycle_starts)) {
    cycle[cycle_starts[i]:cycle_ends[i]] <- i
}
lynx_full$year <- cycle
```

Add lag indicators needed to fit the nonlinear lag models that gave the best one step ahead point forecasts in the ESA workshop example. As in the example, we specify the default argument in the lag function as the mean log population.

```
mean_pop_l = mean(log(lynx_full$population))
lynx_full = dplyr::mutate(lynx_full, popl = log(population),
    lag1 = dplyr::lag(popl, 1, default = mean_pop_l),
    lag2 = dplyr::lag(popl, 2, default = mean_pop_l),
    lag3 = dplyr::lag(popl, 3, default = mean_pop_l),
    lag4 = dplyr::lag(popl, 4, default = mean_pop_l),
    lag5 = dplyr::lag(popl, 5, default = mean_pop_l),
    lag6 = dplyr::lag(popl, 6, default = mean_pop_l))
```

For mvgam models, the response needs to be labelled y and we also need an indicator of the series name as a factor variable

lynx_full$y <- lynx_full$population
lynx_full$series <- factor("series1")

Split the data into training (first 40 years) and testing (next 10 years of data) to evaluate multi-step ahead forecasts

lynx_train = lynx_full[1:40, ]
lynx_test = lynx_full[41:50, ]

The best-forecasting model in the course was with nonlinear smooths of lags 1 and 2; we use those here is that we also include a cyclic smooth for the 19-year cycles as this seems like an important feature, as well as a yearly smooth for the long-term trend. Following the information about spline extrapolation above, we again fit a cubic B spline for the trend with a mix of penalties to try and reign in wacky extrapolation behaviours, and we extend the penalty to cover the years that we wish to predict. This will hopefully give us better uncertainty estimates for the forecast. In this example we assume the observations are Poisson distributed

```
lynx_mgcv = gam(population ~ s(season, bs = "cc",
    k = 19) + s(year, bs = "bs", m = c(3,
    2, 1, 0)) + s(lag1, k = 5) + s(lag2,
    k = 5), knots = list(season = c(0.5,
    19.5), year = c(min(lynx_train$year -
    1), min(lynx_train$year), max(lynx_test$year),
    max(lynx_test$year) + 1)), data = lynx_train,
    family = "poisson", method = "REML")
```

Inspect the model's summary and estimated smooth functions for the season, year and lag terms

```
summary(lynx_mgcv)
##
## Family: poisson
## Link function: log
##
## Formula:
## population ~ s(season, bs = "cc", k = 19) + s(year, bs = "bs",
##     m = c(3, 2, 1, 0)) + s(lag1, k = 5) + s(lag2, k = 5)
##
## Parametric coefficients:
##             Estimate Std. Error z value Pr(>|z|)
## (Intercept) 6.666631   0.007511   887.6   <2e-16 ***
## ---
## Signif. codes:  0 '***' 0.001 '**' 0.01 '*' 0.05 '.' 0.1 ' ' 1
##
## Approximate significance of smooth terms:
##              edf Ref.df Chi.sq p-value
## s(season) 16.229 17.000 6770.2  <2e-16 ***
## s(year)    1.990  2.000  244.2  <2e-16 ***
## s(lag1)    3.984  3.999  712.0  <2e-16 ***
## s(lag2)    3.892  3.993  488.1  <2e-16 ***
## ---
## Signif. codes:  0 '***' 0.001 '**' 0.01 '*' 0.05 '.' 0.1 ' ' 1
##
## R-sq.(adj) =  0.967   Deviance explained = 97.7%
## -REML = 880.28  Scale est. = 1         n = 40
plot(lynx_mgcv, select = 1)
plot(lynx_mgcv, select = 2)
plot(lynx_mgcv, select = 3)
plot(lynx_mgcv, select = 4)
```

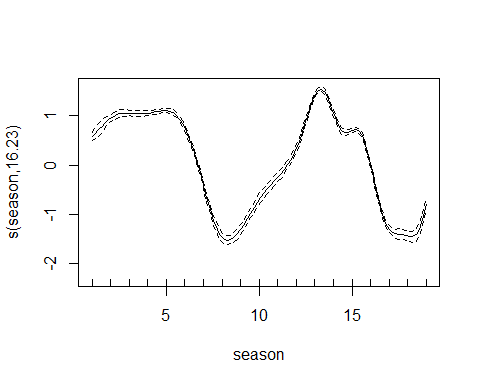

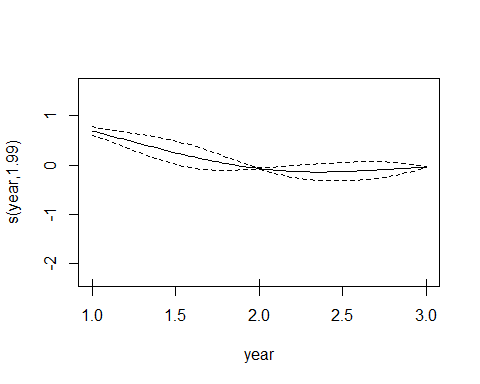

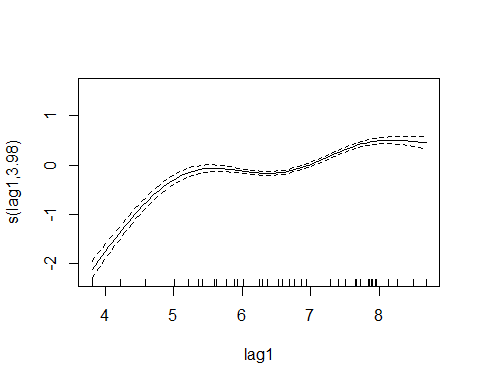

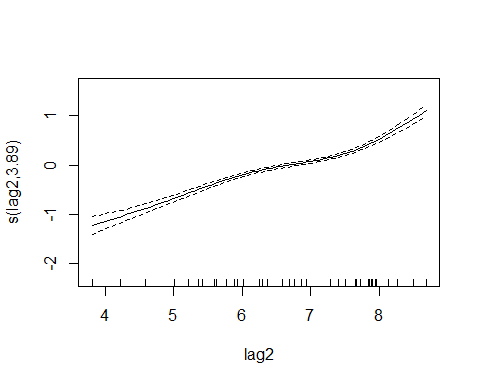

This model captures most of the deviance in the series and the functions are all confidently estimated to be non-zero and non-flat. So far, so good. Now for some forecasts for the out of sample period. First we must take posterior draws of smooth beta coefficients to incorporate the uncertainties around smooth functions when simulating forecast paths

coef_sim <- gam.mh(lynx_mgcv)$bs

Now we define a function to perform forecast simulations from the nonlinear lag model in a recursive fashion. Using starting values for the last two lags, the function will iteratively project the path ahead with a random sample from the model's coefficient posterior

```
recurse_nonlin = function(model, lagged_vals,
    h) {
    # Initiate state vector
    states <- rep(NA, length = h + 2)
    # Last two values of the conditional
    # expectations begin the state vector
    states[1] <- as.numeric(exp(lagged_vals[2]))
    states[2] <- as.numeric(exp(lagged_vals[1]))
    # Get a random sample of the smooth
    # coefficient uncertainty matrix to use
    # for the entire forecast horizon of this
    # particular path
    gam_coef_index <- sample(seq(1, NROW(coef_sim)),
        1, T)
    # For each following timestep,
    # recursively predict based on the
    # predictions at each previous lag
    for (t in 3:(h + 2)) {
        # Build the GAM linear predictor matrix
        # using the two previous lags of the
        # (log) density
        newdata <- data.frame(lag1 = log(states[t -
            1] + 0.01), lag2 = log(states[t -
            2] + 0.01), season = lynx_test$season[t -
            2], year = lynx_test$year[t -
            2])
        colnames(newdata) <- c("lag1", "lag2",
            "season", "year")
        Xp <- predict(model, newdata = newdata,
            type = "lpmatrix")
        # Calculate the posterior prediction for
        # this timepoint
        mu <- rpois(1, lambda = exp(Xp %*%
            coef_sim[gam_coef_index, ]))
        # Fill in the state vector and iterate to
        # the next timepoint
        states[t] <- mu
    }
    # Return the forecast path
    states[-c(1:2)]
}
```

Create the GAM's forecast distribution by generating 1000 simulated forecast paths. Each path is fed the true observed values for the last two lags of the first out of sample timepoint, but they can deviate when simulating ahead depending on their particular draw of possible coefficients. Note, this is a bit slow and could easily be parallelised to speed up computations

```
gam_sims <- matrix(NA, nrow = 1000, ncol = 10)
for (i in 1:1000) {
    gam_sims[i, ] <- recurse_nonlin(lynx_mgcv,
        lagged_vals = c(lynx_test$lag1[1],
            lynx_test$lag2[1]), h = 10)
}
```

Plot the mgcv model's out of sample forecast for the next 10 years ahead

```
cred_ints <- apply(gam_sims, 2, function(x) hpd(x,
    0.95))
yupper <- max(cred_ints) * 1.25
plot(cred_ints[3, ] ~ seq(1:NCOL(cred_ints)),
    type = "l", col = rgb(1, 0, 0, alpha = 0),
    ylim = c(0, yupper), ylab = "Predicted lynx trappings",
    xlab = "Forecast horizon", main = "mgcv")
polygon(c(seq(1:(NCOL(cred_ints))), rev(seq(1:NCOL(cred_ints)))),
    c(cred_ints[1, ], rev(cred_ints[3, ])),
    col = rgb(150, 0, 0, max = 255, alpha = 100),
    border = NA)
cred_ints <- apply(gam_sims, 2, function(x) hpd(x,
    0.68))
polygon(c(seq(1:(NCOL(cred_ints))), rev(seq(1:NCOL(cred_ints)))),
    c(cred_ints[1, ], rev(cred_ints[3, ])),
    col = rgb(150, 0, 0, max = 255, alpha = 180),
    border = NA)
lines(cred_ints[2, ], col = rgb(150, 0, 0,
    max = 255), lwd = 2, lty = "dashed")
points(lynx_test$population[1:10], pch = 16)
lines(lynx_test$population[1:10])
```

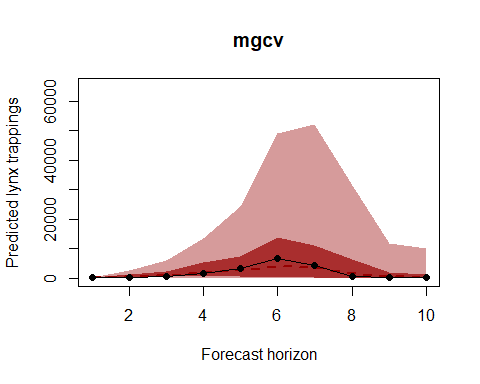

A decent forecast? The shape is certainly correct, but the 95% uncertainty intervals appear to be far too wide (i.e. our upper interval extends to up to 8 times the maximum number of trappings that have ever been recorded up to this point). This is almost entirely due to the extrapolation behaviour of the B spline, as the lag smooth functions are not encountering values very far outside the ranges they've already been trained on so they are resorting mostly to interpolation. But a better way to evaluate than simply using visuals is to calculate a probabilistic score. Here we use the Discrete Rank Probabiilty Score, which gives us an indication of how well calibrated our forecast's uncertainty intervals are by comparing the mass of the forecast density against the true observed values. Forecasts with overly wide intervals are penalised, as are forecasts with overly narrow intervals that do not contain the true observations. At the same time we calculate coverage of the forecast's 90% intervals, which is another useful way of evaluating different forecast proposals

```
# Discrete Rank Probability Score and
# coverage of 90% interval
drps_score <- function(truth, fc, interval_width = 0.9) {
    nsum <- 1000
    Fy = ecdf(fc)
    ysum <- 0:nsum
    indicator <- ifelse(ysum - truth >= 0,
        1, 0)
    score <- sum((indicator - Fy(ysum))^2)

    # Is value within 90% HPD?
    interval <- hpd(fc, interval_width)
    in_interval <- ifelse(truth <= interval[3] &
        truth >= interval[1], 1, 0)
    return(c(score, in_interval))
}

# Wrapper to operate on all observations
# in fc_horizon
drps_mcmc_object <- function(truth, fc, interval_width = 0.9) {
    indices_keep <- which(!is.na(truth))
    if (length(indices_keep) == 0) {
        scores = data.frame(drps = rep(NA,
            length(truth)), interval = rep(NA,
            length(truth)))
    } else {
        scores <- matrix(NA, nrow = length(truth),
            ncol = 2)
        for (i in indices_keep) {
            scores[i, ] <- drps_score(truth = as.vector(truth)[i],
                fc = fc[, i], interval_width)
        }
    }
    scores
}

# Calculate DRPS over the 10-year horizon
# for the mgcv model
lynx_mgcv1_drps <- drps_mcmc_object(truth = lynx_test$population[1:10],
    fc = gam_sims)
```

What if we remove the yearly trend and let the lag smooths capture more of the temporal dependencies? Will that improve the forecast distribution? Run a second model and plot the forecast (note that this plot will be on quite a different y-axis scale compared to the first plot above)

```
lynx_mgcv2 = gam(population ~ s(season, bs = "cc",
    k = 19) + s(lag1, k = 5) + s(lag2, k = 5),
    data = lynx_train, family = "poisson",
    method = "REML")
coef_sim <- gam.mh(lynx_mgcv2)$bs
gam_sims <- matrix(NA, nrow = 1000, ncol = 10)
for (i in 1:1000) {
    gam_sims[i, ] <- recurse_nonlin(lynx_mgcv2,
        lagged_vals = c(lynx_test$lag1[1],
            lynx_test$lag2[1]), h = 10)
}
cred_ints <- apply(gam_sims, 2, function(x) hpd(x,
    0.95))
yupper <- max(cred_ints) * 1.25
plot(cred_ints[3, ] ~ seq(1:NCOL(cred_ints)),
    type = "l", col = rgb(1, 0, 0, alpha = 0),
    ylim = c(0, yupper), ylab = "Predicted lynx trappings",
    xlab = "Forecast horizon", main = "mgcv")
polygon(c(seq(1:(NCOL(cred_ints))), rev(seq(1:NCOL(cred_ints)))),
    c(cred_ints[1, ], rev(cred_ints[3, ])),
    col = rgb(150, 0, 0, max = 255, alpha = 100),
    border = NA)
cred_ints <- apply(gam_sims, 2, function(x) hpd(x,
    0.68))
polygon(c(seq(1:(NCOL(cred_ints))), rev(seq(1:NCOL(cred_ints)))),
    c(cred_ints[1, ], rev(cred_ints[3, ])),
    col = rgb(150, 0, 0, max = 255, alpha = 180),
    border = NA)
lines(cred_ints[2, ], col = rgb(150, 0, 0,
    max = 255), lwd = 2, lty = "dashed")
points(lynx_test$population[1:10], pch = 16)
lines(lynx_test$population[1:10])
```

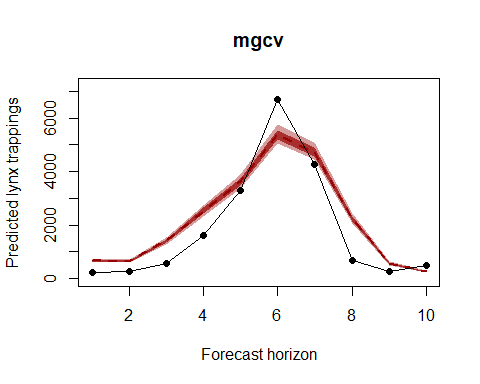

Calculate DRPS over the 10-year horizon for the second mgcv model

```
lynx_mgcv2_drps <- drps_mcmc_object(truth = lynx_test$population[1:10],
    fc = gam_sims)
```

This forecast is highly overconfident, with very unrealistic uncertainty intervals due to the interpolation behaviours of the lag smooths. You can certainly keep trying different formulations (our experience is that the B spline variant above produces the best forecasts from any tested mgcv model, but we did not test an exhaustive set), but hopefully it is clear that forecasting using splines is tricky business and it is likely that each time you do it you'll end up honing in on different combinations of penalties, knot selections etc.... Now we will fit an mvgam model for comparison. This model fits a similar model to the mgcv model directly above but with a full time series model for the errors (in this case an AR1 process), rather than smoothing splines that do not incorporate a concept of the future. We do not use a year term to reduce any possible extrapolation and because the latent dynamic component should capture this temporal variation. We estimate the model in JAGS using MCMC sampling (Note that JAGS 4.3.0 is required; installation links are found [here](https://sourceforge.net/projects/mcmc-jags/files/)).

```
lynx_mvgam <- mvjagam(data_train = lynx_train,
    data_test = lynx_test, formula = y ~
        s(season, bs = "cc", k = 19), knots = list(season = c(0.5,
        19.5)), family = "poisson", trend_model = "AR1",
    burnin = 5000, chains = 4)
## Compiling rjags model...
## Starting 4 rjags simulations using a PSOCK cluster with 4 nodes on host
## 'localhost'
## Simulation complete
## Note: Summary statistics were not produced as there are >50 monitored
## variables
## [To override this behaviour see ?add.summary and ?runjags.options]
## FALSEFinished running the simulation
## NOTE: Stopping adaptation
```

Calculate the out of sample forecast from the fitted mvgam model and plot

```
fits <- MCMCvis::MCMCchains(lynx_mvgam$jags_output,
    "ypred")
fits <- fits[, (NROW(lynx_mvgam$obs_data) +
    1):(NROW(lynx_mvgam$obs_data) + 10)]
cred_ints <- apply(fits, 2, function(x) hpd(x,
    0.95))
yupper <- max(cred_ints) * 1.25
plot(cred_ints[3, ] ~ seq(1:NCOL(cred_ints)),
    type = "l", col = rgb(1, 0, 0, alpha = 0),
    ylim = c(0, yupper), ylab = "", xlab = "Forecast horizon",
    main = "mvgam")
polygon(c(seq(1:(NCOL(cred_ints))), rev(seq(1:NCOL(cred_ints)))),
    c(cred_ints[1, ], rev(cred_ints[3, ])),
    col = rgb(150, 0, 0, max = 255, alpha = 100),
    border = NA)
cred_ints <- apply(fits, 2, function(x) hpd(x,
    0.68))
polygon(c(seq(1:(NCOL(cred_ints))), rev(seq(1:NCOL(cred_ints)))),
    c(cred_ints[1, ], rev(cred_ints[3, ])),
    col = rgb(150, 0, 0, max = 255, alpha = 180),
    border = NA)
lines(cred_ints[2, ], col = rgb(150, 0, 0,
    max = 255), lwd = 2, lty = "dashed")
points(lynx_test$population[1:10], pch = 16)
lines(lynx_test$population[1:10])
```

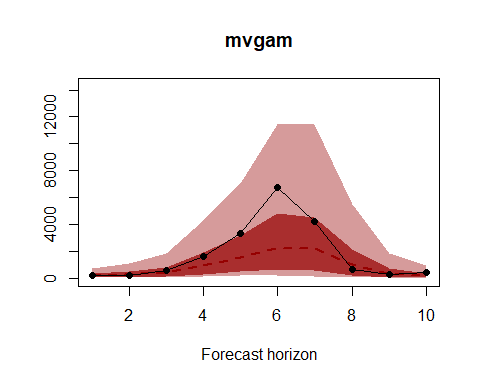

Calculate DRPS over the 10-year horizon for the mvgam model

```
lynx_mvgam_drps <- drps_mcmc_object(truth = lynx_test$population[1:10],
    fc = fits)
```

How do the out of sample DRPS scores stack up for these three models? Remember, our goal is to minimise DRPS while providing 90% intervals that are near, but not less than, 0.9. The DRPS and 90% interval coverage for the first mgcv model (with the B spline year term)

```
sum(lynx_mgcv1_drps[, 1])
## [1] 1549.239
mean(lynx_mgcv1_drps[, 2])
## [1] 0.8
```

For the second mgcv model

```
sum(lynx_mgcv2_drps[, 1])
## [1] 2045.216
mean(lynx_mgcv2_drps[, 2])
## [1] 0
```

And for the mvgam model

```
sum(lynx_mvgam_drps[, 1])
## [1] 730.1052
mean(lynx_mvgam_drps[, 2])
## [1] 1
```

The mvgam has much more realistic uncertainty than the mgcv versions above. Of course this is just one out of sample comparison, and to really determine which model is most appropriate for forecasting we would want to run many of these tests using a [rolling window approach](https://robjhyndman.com/hyndsight/tscv/). Have a look at this model's summary to see what is being estimated (note that longer MCMC runs would probably be needed to increase effective sample sizes)

```
summary_mvgam(lynx_mvgam)
## GAM formula:
## y ~ s(season, bs = "cc", k = 19)
##
## Family:
## Poisson
##
## N series:
## 1
##
## N observations per series:
## 40
##
## GAM smooth term approximate significances:
##             edf Ref.df Chi.sq p-value
## s(season) 15.96  17.00  457.2  <2e-16 ***
## ---
## Signif. codes:  0 '***' 0.001 '**' 0.01 '*' 0.05 '.' 0.1 ' ' 1
##
## GAM coefficient (beta) estimates:
##                    2.5%          50%       97.5% Rhat n.eff
## (Intercept)   6.6309538  6.765705609  6.92252599 1.08   114
## s(season).1  -1.2509540 -0.719730223 -0.07225877 1.28    89
## s(season).2  -0.5058470  0.152328956  0.70670566 1.23    89
## s(season).3   0.2562693  0.883407169  1.37813907 1.21    65
## s(season).4   0.7633532  1.393949443  2.08928493 1.50    53
## s(season).5   0.9030323  1.599838501  2.50803566 2.17    49
## s(season).6   0.2976096  1.075953746  2.06986609 1.61    46
## s(season).7  -0.7581317  0.000952735  0.81794877 1.19    65
## s(season).8  -1.6255559 -0.839499338 -0.18332564 1.14   126
## s(season).9  -1.7347709 -0.881069310 -0.19416546 1.19   103
## s(season).10 -1.1621566 -0.399103407  0.39549291 1.24    71
## s(season).11 -0.3198006  0.433311253  1.22438330 1.34    63
## s(season).12  0.6627665  1.380597334  2.32388686 1.45    32
## s(season).13  1.0027312  1.671777143  2.65014804 1.43    35
## s(season).14  0.8379473  1.309510498  2.08223644 1.27    44
## s(season).15 -0.2957625  0.254361420  0.74392656 1.11    97
## s(season).16 -1.0738876 -0.594373164 -0.06509267 1.02   212
## s(season).17 -1.5721241 -1.080726019 -0.54855987 1.10   154
##
## GAM smoothing parameter (rho) estimates:
##               2.5%      50%    97.5% Rhat n.eff
## s(season) 3.584502 4.472515 5.221236 1.02   957
##
## Latent trend drift (phi) and AR parameter estimates:
##         2.5%       50%     97.5% Rhat n.eff
## phi 0.000000 0.0000000 0.0000000  NaN     0
## ar1 0.454928 0.6850694 0.8896392    1  2908
## ar2 0.000000 0.0000000 0.0000000  NaN     0
## ar3 0.000000 0.0000000 0.0000000  NaN     0
##
```

Now inspect each model's estimated smooth for the 19-year cyclic pattern. Note that the mvgam smooth plot is on a different scale compared to the mgcv plot, but interpretation is similar. The mgcv smooth is much wigglier, likely because it is compensating for any remaining autocorrelation not captured by the lag smooths. We could probably remedy this by reducing k in the seasonal smooth for the mgcv model (in practice this works well, but leaving k larger for the mvgam's seasonal smooth is recommended as our experience is that this tends to lead to better performance and convergence)

plot(lynx_mgcv, select = 1, shade = T)
plot_mvgam_smooth(lynx_mvgam, 1, "season")

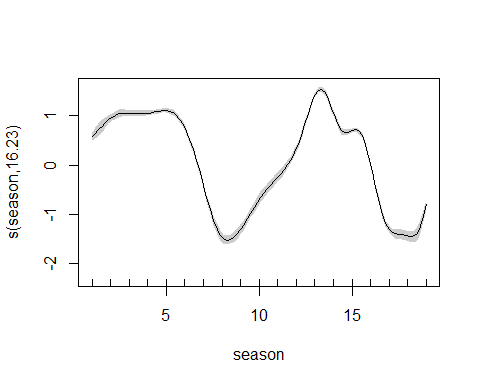

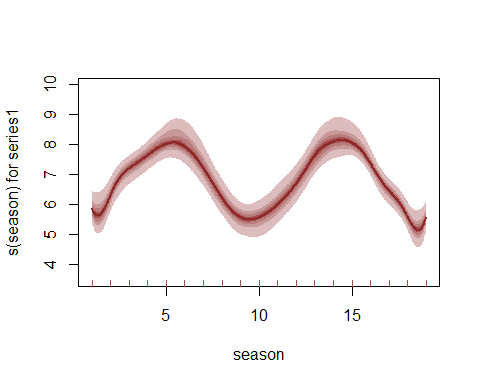

We can also view the mvgam's posterior predictions for the entire series (testing and training)

plot_mvgam_fc(lynx_mvgam, data_test = lynx_test)

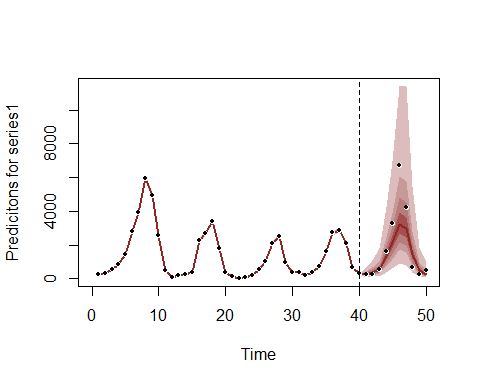

And the estimated trend

plot_mvgam_trend(lynx_mvgam, data_test = lynx_test)

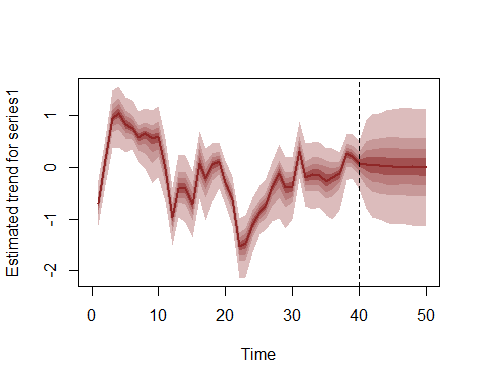

A key aspect of ecological forecasting is to understand [how different components of a model contribute to forecast uncertainty](https://esajournals.onlinelibrary.wiley.com/doi/full/10.1002/eap.1589). We can estimate contributions to forecast uncertainty for the GAM smooth functions and the latent trend using mvgam

plot_mvgam_uncertainty(lynx_mvgam, data_test = lynx_test)

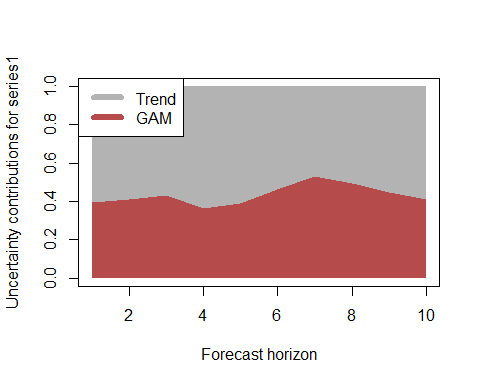

Both components contribute to forecast uncertainty, with the trend component contributing more over time (as it should since this is the stochastic forecast component). This suggests we would still need some more work to learn about factors driving the dynamics of the system. But we will leave the model as-is for this example. Diagnostics of the model can also be performed using mvgam. Have a look at the model's residuals, which are posterior medians of Dunn-Smyth randomised quantile residuals so should follow approximate normality. We are primarily looking for a lack of autocorrelation, which would suggest our AR1 model is appropriate for the latent trend

plot_mvgam_resids(lynx_mvgam)

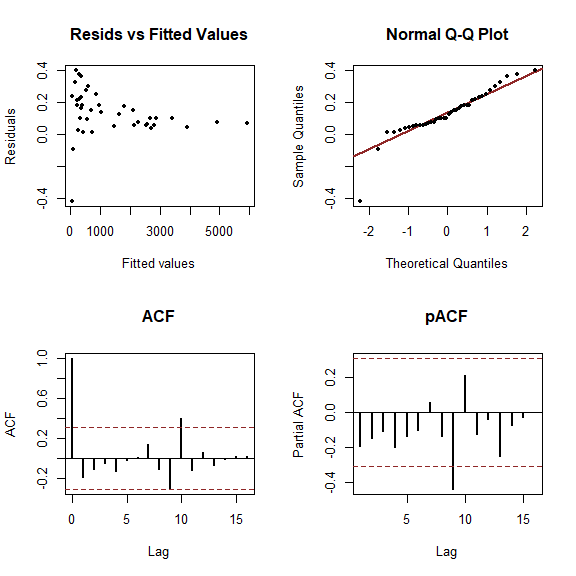
