## Appendix S2 for "Dynamic Generalised Additive Models (DGAM) for forecasting discrete ecological time series"

**APPENDIX S2.** Multivariate dynamic GAMs using the mvgam package for R

Nicholas Clark and Konstans Wells

In this example we will examine multivariate forecasting models using mvgam, which fits GAMs using MCMC sampling via the JAGS software (Note that JAGS 4.3.0 is required; installation links are found [here](https://sourceforge.net/projects/mcmc-jags/files/)). First a simulation experiment to determine whether mvgam's inclusion of complexity penalisation works by reducing the number of un-needed dynamic factors. In any factor model, choosing the appropriate number of factors K can be difficult. The approach used by mvgam is to estimate a penalty for each factor that squeezes the factor's variance toward zero, effectively forcing the factor to evolve as a flat white noise process. By allowing each factor's penalty to be estimated in an exponentially increasing manner (following Welty, Leah J., et al. Bayesian distributed lag models: estimating effects of particulate matter air pollution on daily mortality Biometrics 65.1 (2009): 282-291), we hope that we can guard against specying too large a K. But we should test this property to see how it behaves in different scenarios. Begin by simulating 8 series that evolve with a shared seasonal pattern and that depend on 2 latent random walk factors. Each series is 100 time steps long, with a seasonal frequency of 12. We give the trend moderate importance by setting trend_rel = 0.6 and we allow each series' observation process to be drawn from slightly different Negative Binomial distributions

```
set.seed(100)
library(mvgam)
## Loading required package: mgcv
## Loading required package: nlme
## This is mgcv 1.8-33. For overview type 'help("mgcv-package")'.
## Loading required package: rjags
## Loading required package: coda
## Linked to JAGS 4.3.0
## Loaded modules: basemod,bugs
## Loading required package: parallel
## Loading required package: runjags
## Warning: package 'runjags' was built under R version 4.0.5
dat <- sim_mvgam(T = 100, n_series = 8, n_trends = 2,
    mu_obs = runif(8, 2, 6), size_obs = runif(8,
        0.5, 2), trend_rel = 0.6, train_prop = 0.8)
## Registered S3 method overwritten by 'quantmod':
##   method            from
##   as.zoo.data.frame zoo
```

Have a look at the series

```
par(mfrow = c(4, 2))
for (i in 1:8) {
    plot(dat$data_train$y[which(as.numeric(dat$data_train$series) ==
        i)], type = "l", ylab = paste("Series",
        i), xlab = "Time")
}
```

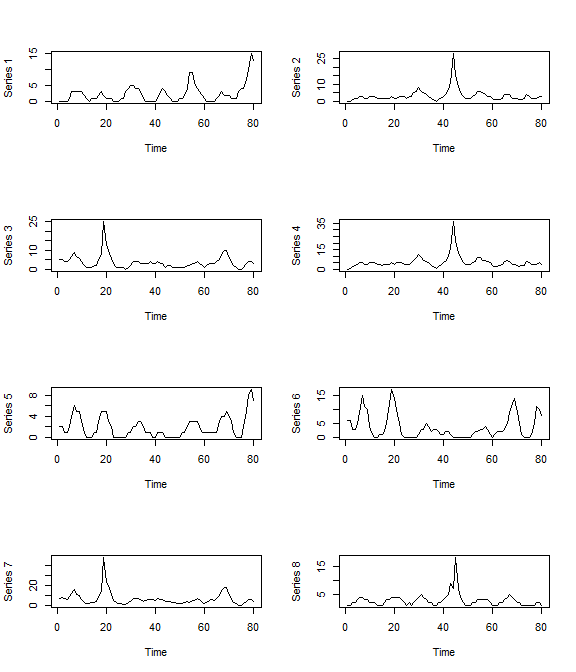

par(mfrow = c(1, 1))

Clearly there are some correlations in the trends for these series. But how does a dynamic factor process allow us to potentially capture these dependencies? The below example demonstrates how. Essentially, a dynamic factor is an *unmeasured* (latent) random process that induces correlations between time series via a set of factor loadings ($\beta$) while exercising dimension reduction. The loadings represent constant associations between the observed time series and the dynamic factor, but each series can still deviate from the factor through its error process and its associations with other factors (if we estimate >1 latent factor in our model).

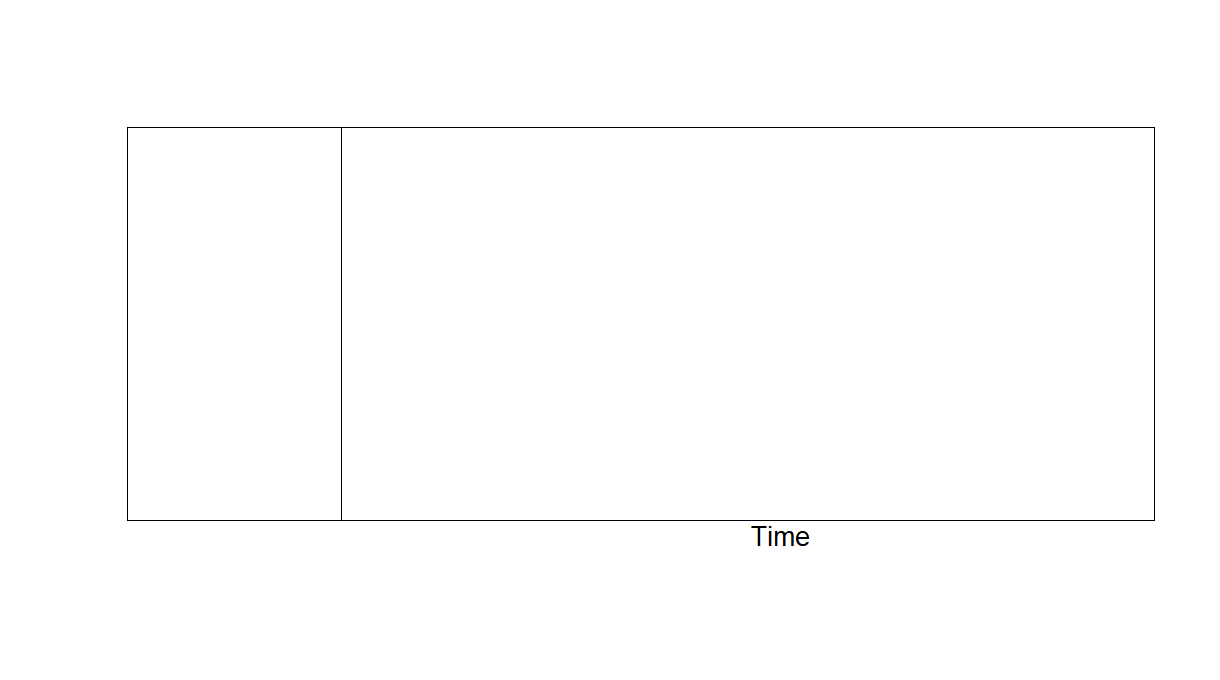

A challenge with any factor model is the need to determine the number of factors K. Setting K too small prevents temporal dependencies from being adequately modelled, leading to poor convergence and difficulty estimating smooth parameters. By contrast, setting K too large leads to unnecessary computation. mvgam approaches this problem by formulating a prior distribution that enforces exponentially increasing penalties on the factor variances to allow any un-needed factors to evolve as flat lines. Now let's fit a well-specified model for our simulated series in which we estimate random intercepts, a shared seasonal cyclic smooth and 2 latent dynamic factors

```
mod1 <- mvjagam(data_train = dat$data_train,
    data_test = dat$data_test, formula = y ~
        s(series, bs = "re") + s(season,
            bs = c("cc"), k = 12), knots = list(season = c(0.5,
        12.5)), use_lv = T, n_lv = 2, family = "nb",
    trend_model = "RW", chains = 4, burnin = 2000)
## Fitting a multivariate GAM with latent dynamic factors for the trends...
## Compiling rjags model...
## Starting 4 rjags simulations using a PSOCK cluster with 4 nodes on host
## 'localhost'
## Simulation complete
## Note: Summary statistics were not produced as there are >50 monitored
## variables
## [To override this behaviour see ?add.summary and ?runjags.options]
## FALSEFinished running the simulation
## NOTE: Stopping adaptation
```

Look at a few plots. The estimated smooth function

```
plot_mvgam_smooth(object = mod1, series = 1,
    smooth = "season")
```

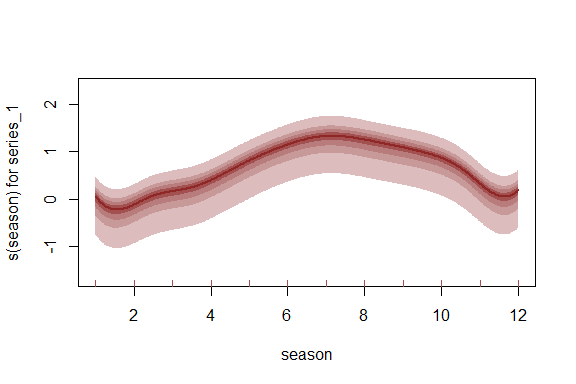

And the true seasonal function in the simulation

plot(dat$global_seasonality[1:12], type = "l")

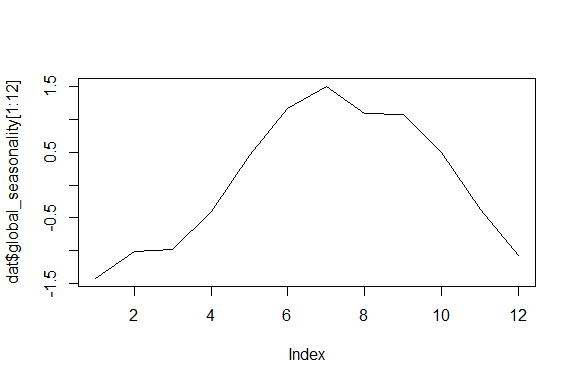

Check whether each factor was retained using the plot_mvgam_factors function. Here, each factor is tested against a null hypothesis of white noise by calculating the sum of the factor's 1st derivatives. A factor that has a larger contribution to the series' latent trends will have a larger sum, both because that factor's absolute magnitudes will be larger (due to the weaker penalty on the factor's precision) and because the factor will move around more. By normalising these estimated first derivative sums, it should be apparent whether some factors have been dropped from the model. Here we see that each factor is contributing to the series' latent trends, and the plots show that neither has been forced to evolve as white noise

plot_mvgam_factors(mod1)

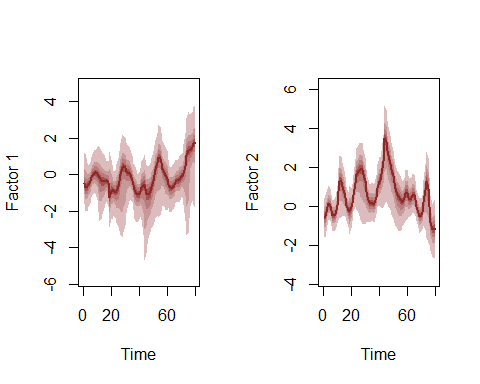

```
##         Contribution
## Factor1    0.3240424
## Factor2    0.6759576
```

Now we fit the same model but assume that we no nothing about how many factors to use, so we specify the maximum allowed (the total number of series; 8). Note that this model is computationally more expensive so it will take longer to fit

```
mod2 <- mvjagam(data_train = dat$data_train,
    data_test = dat$data_test, formula = y ~
        s(series, bs = "re") + s(season,
            bs = c("cc"), k = 12), knots = list(season = c(0.5,
        12.5)), use_lv = T, n_lv = 8, family = "nb",
    trend_model = "RW", chains = 4, burnin = 2000)
## Fitting a multivariate GAM with latent dynamic factors for the trends...
## Compiling rjags model...
## Starting 4 rjags simulations using a PSOCK cluster with 4 nodes on host
## 'localhost'
## Simulation complete
## Note: Summary statistics were not produced as there are >50 monitored
## variables
## [To override this behaviour see ?add.summary and ?runjags.options]
## FALSEFinished running the simulation
## NOTE: Stopping adaptation
```

The same plots as model 1 to see if this model has also fit the data well

```
plot_mvgam_smooth(object = mod2, series = 1,
    smooth = "season")
```

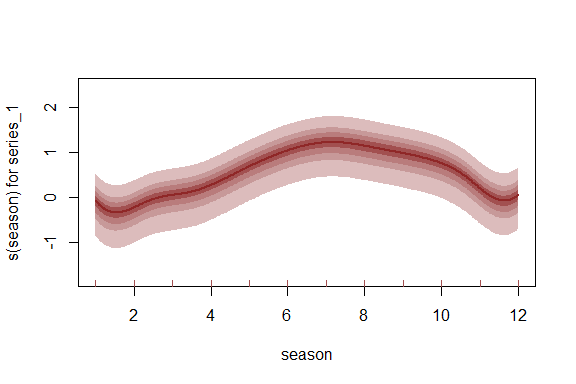

plot(dat$global_seasonality[1:12], type = "l")

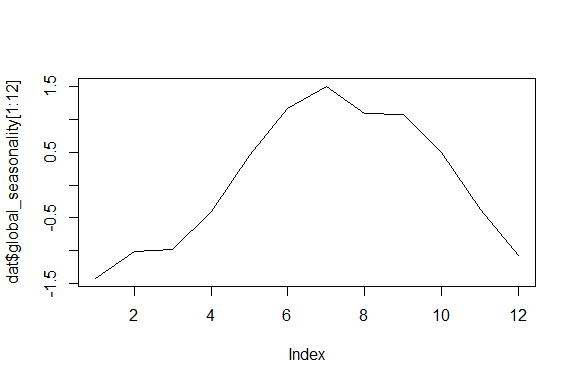

Examining the factor contributions gives us some insight into whether we set n_lv larger than we perhaps needed to. These contributions can be interpreted similarly to ordination axes when deciding how many latent variables to specify

plot_mvgam_factors(mod2)

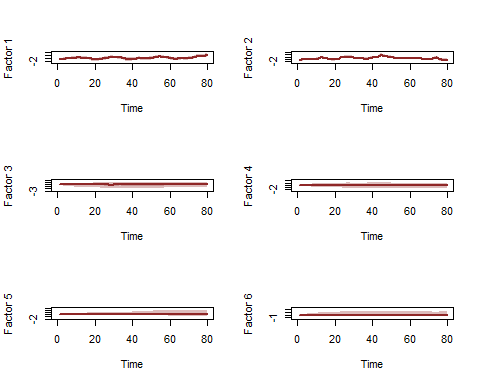

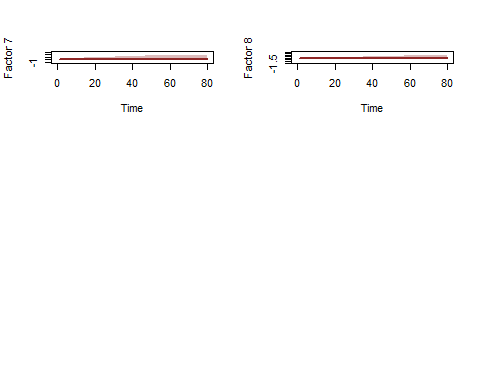

```
##         Contribution
## Factor1  0.343726370
## Factor2  0.604954597
## Factor3  0.017647869
## Factor4  0.010857205
## Factor5  0.008953251
## Factor6  0.005164580
## Factor7  0.005328183
## Factor8  0.003367945
```

The very weak contributions by some of the factors are a result of the penalisation, which will become more important as the dimensionality of the data grows. Now onto an empirical example. Here we will access monthly search volume data from Google Trends, focusing on relative importances of search terms related to tick paralysis in Queensland, Australia

```
library(mvgam)
library(tidyr)
if (!require(gtrendsR)) {
    install.packages("gtrendsR")
}

terms = c("tick bite", "tick paralysis",
    "dog tick", "paralysis tick dog")
trends <- gtrendsR::gtrends(terms, geo = "AU-QLD",
    time = "all", onlyInterest = T)
```

Google Trends modified their algorithm for extracting search volume data in 2012, so we filter the series to only include observations after this point in time

```
gtest <- trends$interest_over_time %>% tidyr::spread(keyword,
    hits) %>% dplyr::select(-geo, -time,
    -gprop, -category) %>% dplyr::mutate(date = lubridate::ymd(date)) %>%
    dplyr::mutate(year = lubridate::year(date)) %>%
    dplyr::filter(year > 2012) %>% dplyr::select(-year)
```

Convert to an xts object and then to the required mvgam format, holding out the final 10% of observations as the test data

```
series <- xts::xts(x = gtest[, -1], order.by = gtest$date)
trends_data <- series_to_mvgam(series, freq = 12,
    train_prop = 0.9)
```

Plot the series to see how similar their seasonal shapes are over time

plot(series, legend.loc = "topleft")

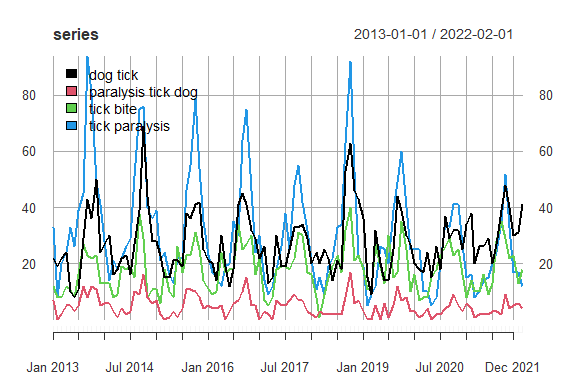

Now we will fit an mvgam model with shared seasonality and random intercepts per series. Our first attempt will ignore any temporal component in the residuals so that we can identidy which GAM predictor combination gives us the best fit, prior to investigating how to deal with any remaining autocorrelation. We assume a Negative Binomial distrubution with series-specific overdispersion parameters for the response. We use a complexity-penalising prior for the overdispersion, which allows the model to reduce toward a more simple Poisson observation process unless the data provide adequate information to support overdispersion. Also note that any smooths using the random effects basis (s(series, bs = "re") below) are automatically re-parameterised to use the [non-centred parameterisation that is necessary to help avoid common posterior degeneracies in hierarchical models](https://betanalpha.github.io/assets/case_studies/hierarchical_modeling.html). We suppress the global intercept as it is not needed and will lead to identifiability issues when estimating the series-specific random intercepts

```
trends_mod1 <- mvjagam(data_train = trends_data$data_train,
    data_test = trends_data$data_test, formula = y ~
        s(series, bs = "re") + s(season,
            k = 12, m = 2, bs = "cc") - 1,
    knots = list(season = c(0.5, 12.5)),
    trend_model = "None", family = "nb",
    chains = 4, burnin = 8000)
## Compiling rjags model...
## Starting 4 rjags simulations using a PSOCK cluster with 4 nodes on host
## 'localhost'
## Simulation complete
## Note: Summary statistics were not produced as there are >50 monitored
## variables
## [To override this behaviour see ?add.summary and ?runjags.options]
## FALSEFinished running the simulation
## NOTE: Stopping adaptation
```

Given that these series could potentially be following a hierarchical seasonality, we will also trial a slghtly more complex model with an extra smoothing term per series that allows its seasonal curve to deviate from the global seasonal smooth. Ignore the warning about repeated smooths, as this is not an issue for estimation.

```
trends_mod2 <- mvjagam(data_train = trends_data$data_train,
    data_test = trends_data$data_test, formula = y ~
        s(season, k = 12, m = 2, bs = "cc") +
            s(season, series, k = 5, bs = "fs",
                m = 1), knots = list(season = c(0.5,
        12.5)), trend_model = "None", family = "nb",
    chains = 4, burnin = 8000)
## Compiling rjags model...
## Starting 4 rjags simulations using a PSOCK cluster with 4 nodes on host
## 'localhost'
## Simulation complete
## Note: Summary statistics were not produced as there are >50 monitored
## variables
## [To override this behaviour see ?add.summary and ?runjags.options]
## FALSEFinished running the simulation
## NOTE: Stopping adaptation
```

How can we compare these models to ensure we choose one that performs well and provides useful inferences? Beyond posterior retrodictive and predictive checks, we can take advantage of the fact that mvgam fits an mgcv model to provide all the necessary penalty matrices, as well as to identify good initial values for smoothing and beta parameters. Because we did not modify this model by adding a trend component (the only modification is that we estimated series-specific overdispersion parameters), we can still inspect the usual mgcv model comparison routines

```
anova(trends_mod1$mgcv_model, trends_mod2$mgcv_model,
    test = "LRT")
## Analysis of Deviance Table
##
## Model 1: y ~ s(series, bs = "re") + s(season, k = 12, m = 2, bs = "cc") -
##     1
## Model 2: y ~ s(season, k = 12, m = 2, bs = "cc") + s(season, series, k = 5,
##     bs = "fs", m = 1)
##   Resid. Df Resid. Dev     Df Deviance  Pr(>Chi)
## 1    383.41     2463.1
## 2    371.72     2384.0 11.695   79.081 4.546e-12 ***
## ---
## Signif. codes:  0 '***' 0.001 '**' 0.01 '*' 0.05 '.' 0.1 ' ' 1
AIC(trends_mod1$mgcv_model, trends_mod2$mgcv_model)
##                              df      AIC
## trends_mod1$mgcv_model 13.28318 2489.630
## trends_mod2$mgcv_model 23.17777 2430.339
summary(trends_mod1$mgcv_model)
##
## Family: Negative Binomial(18.21)
## Link function: log
##
## Formula:
## y ~ s(series, bs = "re") + s(season, k = 12, m = 2, bs = "cc") -
##     1
##
## Approximate significance of smooth terms:
##             edf Ref.df     F p-value
## s(series) 3.999      4  7455  <2e-16 ***
## s(season) 6.927     10 12298  <2e-16 ***
## ---
## Signif. codes:  0 '***' 0.001 '**' 0.01 '*' 0.05 '.' 0.1 ' ' 1
##
## R-sq.(adj) =  0.785   Deviance explained = 81.1%
## fREML = 600.14  Scale est. = 1         n = 396
summary(trends_mod2$mgcv_model)
##
## Family: Negative Binomial(29.34)
## Link function: log
##
## Formula:
## y ~ s(season, k = 12, m = 2, bs = "cc") + s(season, series, k = 5,
##     bs = "fs", m = 1)
##
## Parametric coefficients:
##             Estimate Std. Error t value Pr(>|t|)
## (Intercept)   2.7046     0.4374   6.183 1.64e-09 ***
## ---
## Signif. codes:  0 '***' 0.001 '**' 0.01 '*' 0.05 '.' 0.1 ' ' 1
##
## Approximate significance of smooth terms:
##                     edf Ref.df     F p-value
## s(season)         6.668     10 12.77  <2e-16 ***
## s(season,series) 12.923     19 65.73  <2e-16 ***
## ---
## Signif. codes:  0 '***' 0.001 '**' 0.01 '*' 0.05 '.' 0.1 ' ' 1
##
## R-sq.(adj) =  0.825   Deviance explained = 84.6%
## fREML = 612.76  Scale est. = 1         n = 396
```

WE can also use JAGS routines for interrogating models. Which model minimises in-sample DIC?

```
runjags::extract(trends_mod1$jags_model,
    "dic")
## Compiling rjags model and adapting for 1000 iterations...
## Obtaining DIC samples from 2000 iterations...
## Mean deviance:  2479
## penalty 16.34
## Penalized deviance: 2495
runjags::extract(trends_mod2$jags_model,
    "dic")
## Compiling rjags model and adapting for 1000 iterations...
## Obtaining DIC samples from 2000 iterations...
## Mean deviance:  2407
## penalty 26.5
## Penalized deviance: 2433
```

Model 2 seems to fit better so far, suggesting that hierarchical seasonality gives better performance for these series. But a problem with both of the above models is that their forecast uncertainties will not increase into the future, which is not how time series forecasts should behave. Here we fit Model 2 again but specifying a time series model for the latent trends. We assume the dynamic trends can be represented using latent factors that each follow an AR1 process, and we will rely on the exponential penalties to help regularise any un-needed factors by setting n_lv = 4

```
trends_mod3 <- mvjagam(data_train = trends_data$data_train,
    data_test = trends_data$data_test, formula = y ~
        s(season, k = 12, m = 2, bs = "cc") +
            s(season, series, k = 5, bs = "fs",
                m = 1), knots = list(season = c(0.5,
        12.5)), trend_model = "AR1", use_lv = T,
    n_lv = 4, family = "nb", chains = 4,
    burnin = 8000)
## Fitting a multivariate GAM with latent dynamic factors for the trends...
## Compiling rjags model...
## Starting 4 rjags simulations using a PSOCK cluster with 4 nodes on host
## 'localhost'
## Simulation complete
## Note: Summary statistics were not produced as there are >50 monitored
## variables
## [To override this behaviour see ?add.summary and ?runjags.options]
## FALSEFinished running the simulation
## NOTE: Stopping adaptation
```

Have a look at the returned JAGS model file to see how the dynamic factors are incorporated. Notice that the precisions of factors, together with each factor's set of loadings, are penalised using a regularised horseshoe prior

```
trends_mod3$model_file
##   [1] "model {"
##   [2] ""
##   [3] "        ## GAM linear predictor"
##   [4] "        eta <- X %*% b"
##   [5] ""
##   [6] "        ## Mean expectations"
##   [7] "        for (i in 1:n) {"
##   [8] "        for (s in 1:n_series) {"
##   [9] "        mu[i, s] <- exp(eta[ytimes[i, s]] + trend[i, s])"
##  [10] "        }"
##  [11] "        }"
##  [12] ""
##  [13] "        ## Latent factors evolve as time series with penalised precisions;"
##  [14] "        ## the penalty terms force any un-needed factors to evolve as flat lines"
##  [15] "        for(j in 1:n_lv){"
##  [16] "         LV[1, j] ~ dnorm(0, penalty[j])"
##  [17] "        }"
##  [18] ""
##  [19] "        for(j in 1:n_lv){"
##  [20] "         LV[2, j] ~ dnorm(phi[j] + ar1[j]*LV[1, j], penalty[j])"
##  [21] "        }"
##  [22] ""
##  [23] "        for(j in 1:n_lv){"
##  [24] "         LV[3, j] ~ dnorm(phi[j] + ar1[j]*LV[2, j] + ar2[j]*LV[1, j], penalty[j])"
##  [25] "        }"
##  [26] ""
##  [27] "        for(i in 4:n){"
##  [28] "         for(j in 1:n_lv){"
##  [29] "          LV[i, j] ~ dnorm(phi[j] + ar1[j]*LV[i - 1, j] +"
##  [30] "                          ar2[j]*LV[i - 2, j] + ar3[j]*LV[i - 3, j], penalty[j])"
##  [31] "         }"
##  [32] "        }"
##  [33] ""
##  [34] "        ## AR components"
##  [35] "        for (s in 1:n_lv){"
##  [36] "        phi[s] <- 0"
##  [37] "        ar1[s] ~ dnorm(0, 10)"
##  [38] "        ar2[s] <- 0"
##  [39] "        ar3[s] <- 0"
##  [40] "        }"
##  [41] ""
##  [42] "        ## Shrinkage penalties for each factor squeeze the factor to a flat line and squeeze"
##  [43] "        ## the entire factor toward a flat white noise process if supported by"
##  [44] "        ## the data. The prior for individual factor penalties allows each factor to possibly"
##  [45] "        ## have a relatively large penalty, which shrinks the prior for that factor's variance"
##  [46] "        ## substantially. Penalties increase exponentially with the number of factors following"
##  [47] "        ## Welty, Leah J., et al. Bayesian distributed lag models: estimating effects of particulate"
##  [48] "        ## matter air pollution on daily mortality Biometrics 65.1 (2009): 282-291."
##  [49] "        pi ~ dunif(0, n_lv)"
##  [50] "        X2 ~ dnorm(0, 1)T(0, )"
##  [51] ""
##  [52] "        # eta1 controls the baseline penalty"
##  [53] "        eta1 ~ dunif(-1, 1)"
##  [54] ""
##  [55] "        # eta2 controls how quickly the penalties exponentially increase"
##  [56] "        eta2 ~ dunif(-1, 1)"
##  [57] ""
##  [58] "        for(t in 1:n_lv){"
##  [59] "         X1[t] ~ dnorm(0, 1)T(0, )"
##  [60] "         l.dist[t] <- max(t, pi[])"
##  [61] "         l.weight[t] <- exp(eta2[] * l.dist[t])"
##  [62] "         l.var[t] <- exp(eta1[] * l.dist[t] / 2) * 1"
##  [63] "         theta.prime[t] <- l.weight[t] * X1[t] + (1 - l.weight[t]) * X2[]"
##  [64] "         penalty[t] <- max(min_eps, theta.prime[t] * l.var[t])"
##  [65] "        }"
##  [66] ""
##  [67] "        ## Latent factor loadings: standard normal with identifiability constraints"
##  [68] "        ## Upper triangle of loading matrix set to zero"
##  [69] "        for(j in 1:(n_lv - 1)){"
##  [70] "          for(j2 in (j + 1):n_lv){"
##  [71] "           lv_coefs[j, j2] <- 0"
##  [72] "          }"
##  [73] "         }"
##  [74] ""
##  [75] "        ## Positive constraints on loading diagonals"
##  [76] "        for(j in 1:n_lv) {"
##  [77] "         lv_coefs[j, j] ~ dnorm(0, 1)T(0, 1);"
##  [78] "        }"
##  [79] ""
##  [80] "        ## Lower diagonal free"
##  [81] "        for(j in 2:n_lv){"
##  [82] "         for(j2 in 1:(j - 1)){"
##  [83] "          lv_coefs[j, j2] ~ dnorm(0, 1)T(-1, 1);"
##  [84] "         }"
##  [85] "       }"
##  [86] ""
##  [87] "        ## Other elements also free"
##  [88] "        for(j in (n_lv + 1):n_series) {"
##  [89] "         for(j2 in 1:n_lv){"
##  [90] "          lv_coefs[j, j2] ~ dnorm(0, 1)T(-1, 1);"
##  [91] "         }"
##  [92] "        }"
##  [93] ""
##  [94] "        ## Trend evolution for the series depends on latent factors"
##  [95] "        for (i in 1:n){"
##  [96] "        for (s in 1:n_series){"
##  [97] "         trend[i, s] <- inprod(lv_coefs[s,], LV[i,])"
##  [98] "        }"
##  [99] "        }"
## [100] ""
## [101] "        ## Negative binomial likelihood functions"
## [102] "        for (i in 1:n) {"
## [103] "        for (s in 1:n_series) {"
## [104] "       y[i, s] ~ dnegbin(rate[i, s], r[s])"
## [105] "        rate[i, s] <- ifelse((r[s] / (r[s] + mu[i, s])) < min_eps, min_eps,"
## [106] "                            (r[s] / (r[s] + mu[i, s])))"
## [107] "        }"
## [108] "        }"
## [109] ""
## [110] "        ## Complexity penalising prior for the overdispersion parameter;"
## [111] "        ## where the likelihood reduces to a 'base' model (Poisson) unless"
## [112] "        ## the data support overdispersion"
## [113] "        for(s in 1:n_series){"
## [114] "         r[s] <- pow(r_raw[s], 2)"
## [115] "         r_raw[s] ~ dexp(0.05)"
## [116] "        }"
## [117] ""
## [118] "        ## Posterior predictions"
## [119] "        for (i in 1:n) {"
## [120] "        for (s in 1:n_series) {"
## [121] "       ypred[i, s] ~ dnegbin(rate[i, s], r[s])"
## [122] "        }"
## [123] "        }"
## [124] "        "
## [125] "  ## parametric effect priors (regularised for identifiability)"
## [126] "  for (i in 1:1) { b[i] ~ dnorm(p_coefs[i], p_taus[i]) }"
## [127] "  ## prior for s(season)... "
## [128] "  K1 <- S1[1:10,1:10] * lambda[1] "
## [129] "  b[2:11] ~ dmnorm(zero[2:11],K1) "
## [130] "  ## prior for s(season,series)... "
## [131] "  for (i in c(12:15,17:20,22:25,27:30)) { b[i] ~ dnorm(0, lambda[2]) }"
## [132] "  for (i in c(16,21,26,31)) { b[i] ~ dnorm(0, lambda[3]) }"
## [133] "   ## smoothing parameter priors..."
## [134] "  for (i in 1:3) {"
## [135] "   lambda[i] ~ dexp(1/sp[i])"
## [136] "    rho[i] <- log(lambda[i])"
## [137] "  }"
## [138] "}"
```

Inspection of the dynamic factors and their relative contributions indicates that the first factor is by far the most important

plot_mvgam_factors(trends_mod3)

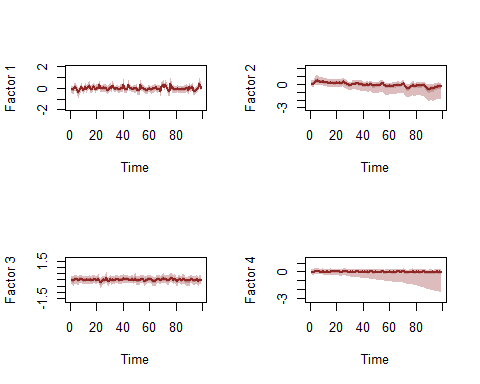

```
##         Contribution
## Factor1    0.5105248
## Factor2    0.1793097
## Factor3    0.2028833
## Factor4    0.1072822
```

How do forecasts for this model compare to the previous one that did not include any trend component?

```
compare_mvgams(trends_mod2, trends_mod3,
    fc_horizon = 6, n_cores = 3, n_evaluations = 20)
## DRPS summaries per model (lower is better)
##              Min.   1st Qu.   Median     Mean  3rd Qu.     Max.
## Model 1 11.719772 16.242471 20.09782 23.10294 27.45231 48.19016
## Model 2  4.288775  7.695406 11.07234 12.54388 15.19869 43.78522
##
## 90% interval coverages per model (closer to 0.9 is better)
## Model 1 1
## Model 2 0.91875
```

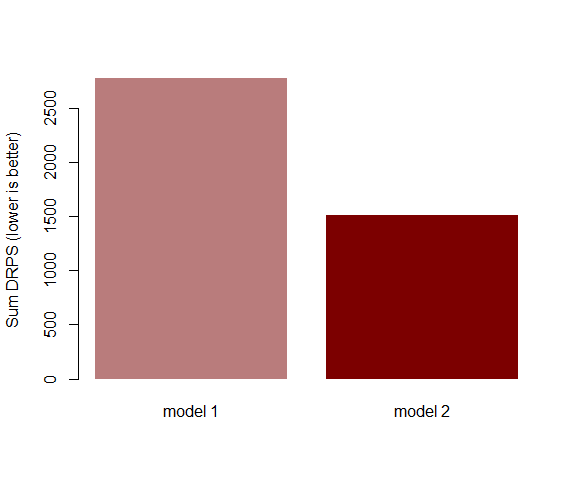

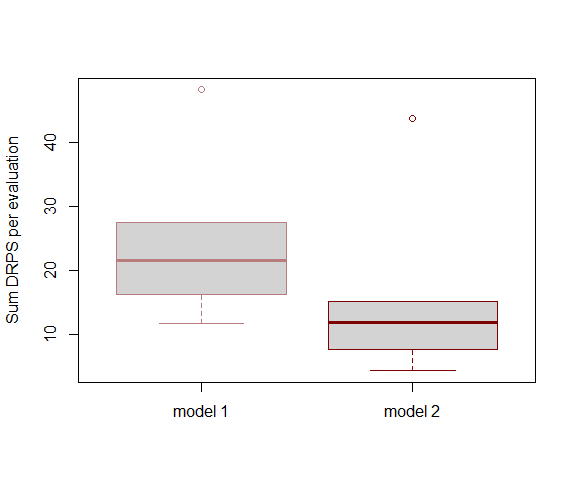

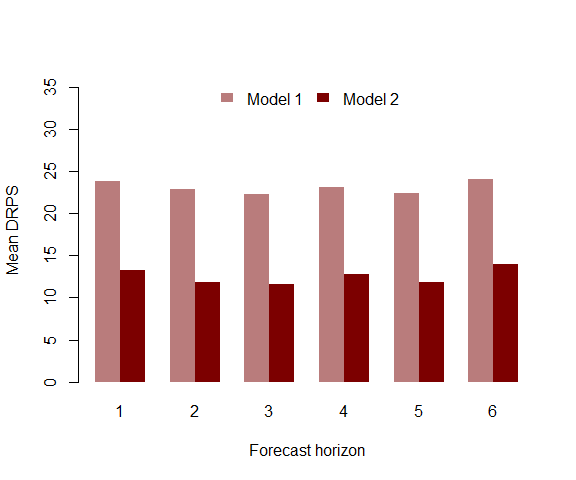

Model 3 (with the dynamic trend) provides far superior forecasts than relying only on the estimated smooths. Inspect the model summary (note again that p-value approximations are a work in progress here and so may not be totally reliable).

```
summary_mvgam(trends_mod3)
## GAM formula:
## y ~ s(season, k = 12, m = 2, bs = "cc") + s(season, series, k = 5,
##     bs = "fs", m = 1)
##
## Family:
## Negative Binomial
##
## N latent factors:
## 4
##
## N series:
## 4
##
## N observations per series:
## 99
##
## Dispersion parameter estimates:
##          2.5%      50%    97.5% Rhat n.eff
## r[1] 52.96422 540.0562 7190.867 1.01  4512
## r[2] 39.67143 520.2147 7346.430 1.01  4195
## r[3] 29.16466 277.0741 5656.721 1.03  3039
## r[4] 53.12430 510.2104 6635.951 1.01  4077
##
## GAM smooth term approximate significances:
##                     edf Ref.df     F p-value
## s(season)         7.759 10.000 18.92  <2e-16 ***
## s(season,series)  9.857 20.000 59.58  <2e-16 ***
## ---
## Signif. codes:  0 '***' 0.001 '**' 0.01 '*' 0.05 '.' 0.1 ' ' 1
##
## GAM coefficient (beta) estimates:
##                            2.5%          50%       97.5% Rhat n.eff
## (Intercept)          2.03818690  2.981472346  3.88329880 1.59    19
## s(season).1         -0.53646687 -0.368471250 -0.16127914 1.01    94
## s(season).2         -0.88731670 -0.619064966 -0.36908567 1.07    84
## s(season).3         -0.80479436 -0.506536566 -0.25691687 1.09    85
## s(season).4         -0.57124956 -0.278218084  0.05149052 1.10    62
## s(season).5         -0.49399967 -0.211418012  0.15917542 1.13    55
## s(season).6         -0.19180208  0.065634454  0.43212975 1.05    58
## s(season).7          0.34784091  0.586055228  0.86416703 1.14    82
## s(season).8          0.50798430  0.739529079  0.97939552 1.19    89
## s(season).9          0.36637827  0.565677328  0.76486253 1.09    82
## s(season).10        -0.08052955  0.165625661  0.38920461 1.06    66
## s(season,series).1  -0.16588866  0.031085852  0.24795060 1.12   144
## s(season,series).2  -0.38794206 -0.164065700  0.05715237 1.02   152
## s(season,series).3  -0.41547290 -0.142973365  0.04916587 1.06    57
## s(season,series).4   0.01597268  0.114489207  0.21302030 1.11   104
## s(season,series).5  -0.60724390  0.325387582  1.26780252 1.60    22
## s(season,series).6  -0.45486120 -0.198540974  0.05030305 1.06   334
## s(season,series).7  -0.26603055  0.015676164  0.29937699 1.00   554
## s(season,series).8  -0.29067230 -0.022166693  0.19434285 1.05   134
## s(season,series).9  -0.14203196 -0.029575602  0.08812504 1.06   200
## s(season,series).10 -2.46050761 -1.484476502 -0.54149734 1.55    35
## s(season,series).11 -0.06928346  0.135132818  0.36407215 1.12   165
## s(season,series).12 -0.13992257  0.094798654  0.33118634 1.01   210
## s(season,series).13 -0.13513915  0.146009373  0.34941917 1.06    70
## s(season,series).14 -0.02938660  0.076656021  0.18064757 1.11   112
## s(season,series).15 -1.22602096 -0.321755266  0.61903357 1.60    24
## s(season,series).16 -0.34667743 -0.137392749  0.08058991 1.12   134
## s(season,series).17 -0.14875931  0.074014803  0.29729062 1.02   167
## s(season,series).18 -0.27145928  0.005394192  0.20035978 1.06    74
## s(season,series).19 -0.12588790 -0.028998872  0.06830106 1.09   132
## s(season,series).20 -0.59326275  0.425013019  1.32652625 1.49    25
##
## GAM smoothing parameter (rho) estimates:
##                        2.5%       50%      97.5% Rhat n.eff
## s(season)          4.336378  5.430925  6.3554268 1.01   733
## s(season,series)   2.116037  2.917922  3.5796902 1.00  1585
## s(season,series)2 -3.469179 -1.998967 -0.9881267 1.00  7850
##
## Latent trend drift (phi) and AR parameter estimates:
##               2.5%        50%     97.5% Rhat n.eff
## phi[1]  0.00000000 0.00000000 0.0000000  NaN     0
## phi[2]  0.00000000 0.00000000 0.0000000  NaN     0
## phi[3]  0.00000000 0.00000000 0.0000000  NaN     0
## phi[4]  0.00000000 0.00000000 0.0000000  NaN     0
## ar1[1] -0.22694839 0.08859013 0.3922679 1.00  4613
## ar1[2] -0.05639405 0.74993073 1.0146659 1.28   245
## ar1[3] -0.41566974 0.05797748 0.5204217 1.00  1918
## ar1[4] -0.47890400 0.11156277 1.0306890 1.39   860
## ar2[1]  0.00000000 0.00000000 0.0000000  NaN     0
## ar2[2]  0.00000000 0.00000000 0.0000000  NaN     0
## ar2[3]  0.00000000 0.00000000 0.0000000  NaN     0
## ar2[4]  0.00000000 0.00000000 0.0000000  NaN     0
## ar3[1]  0.00000000 0.00000000 0.0000000  NaN     0
## ar3[2]  0.00000000 0.00000000 0.0000000  NaN     0
## ar3[3]  0.00000000 0.00000000 0.0000000  NaN     0
## ar3[4]  0.00000000 0.00000000 0.0000000  NaN     0
##
```

Look at Dunn-Smyth residuals for some series from this preferred model to ensure that our dynamic factor process has captured most of the temporal dependencies in the observations

plot_mvgam_resids(trends_mod3, 1)

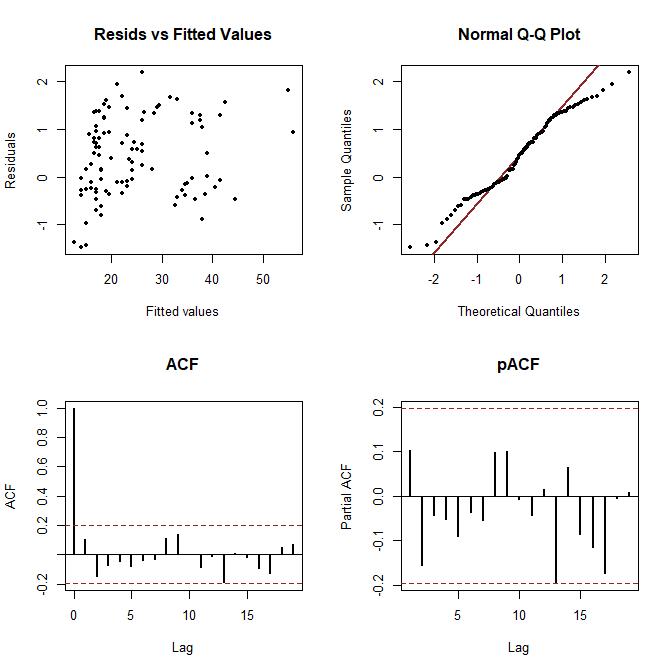

plot_mvgam_resids(trends_mod3, 2)

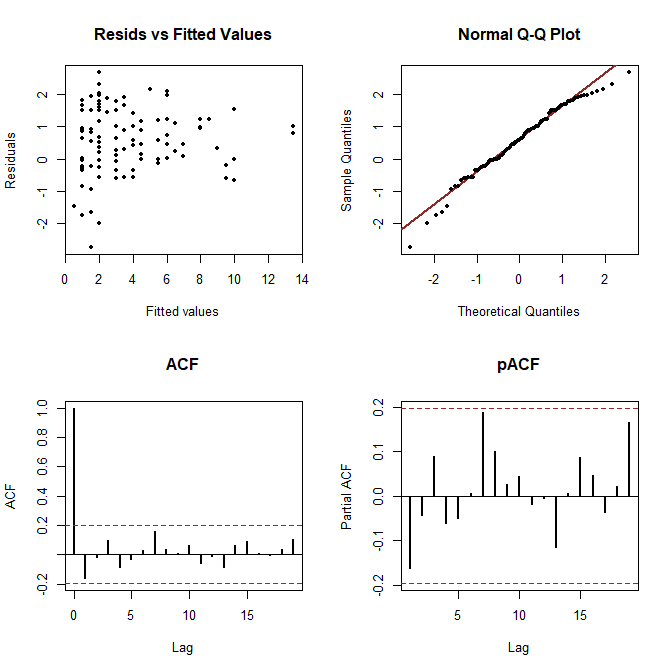

plot_mvgam_resids(trends_mod3, 3)

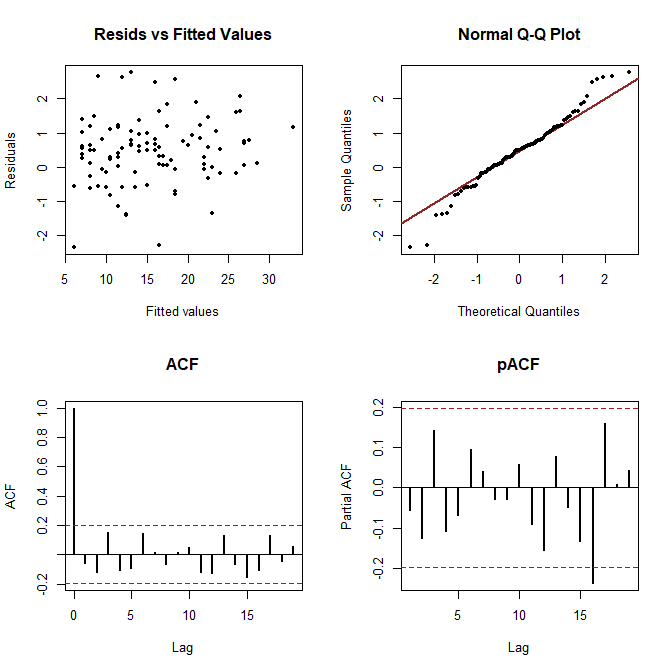

plot_mvgam_resids(trends_mod3, 4)

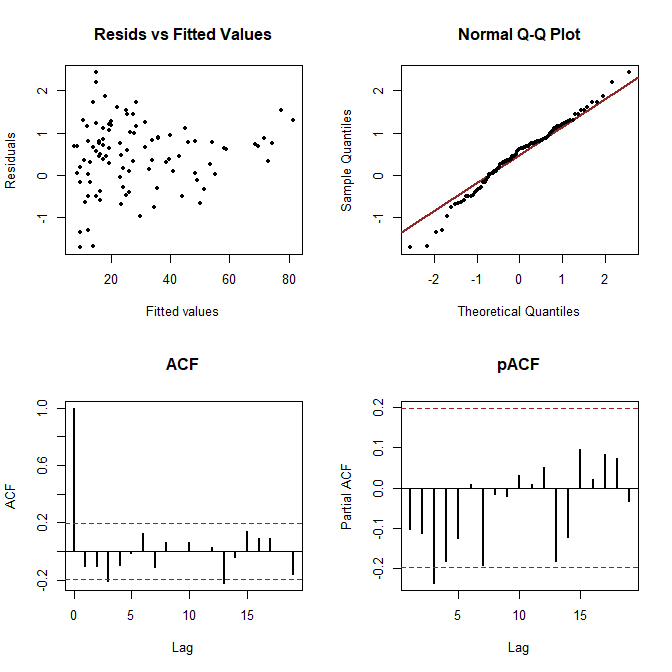

Perform posterior predictive checks to see if the model is able to simulate data that looks realistic and unbiased by examining simulated kernel densities for posterior predictions (yhat) compared to the density of the observations (y). This will be particularly useful for examining whether the Negative Binomial observation model is able to produce realistic looking simulations for each individual series.

plot_mvgam_ppc(trends_mod3, series = 1, type = "density")

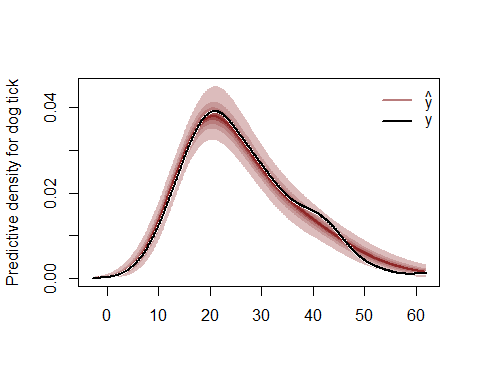

plot_mvgam_ppc(trends_mod3, series = 2, type = "density")

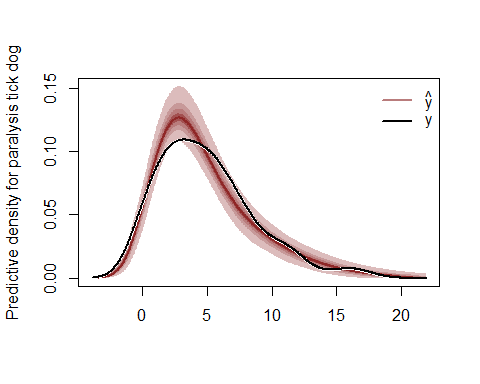

plot_mvgam_ppc(trends_mod3, series = 3, type = "density")

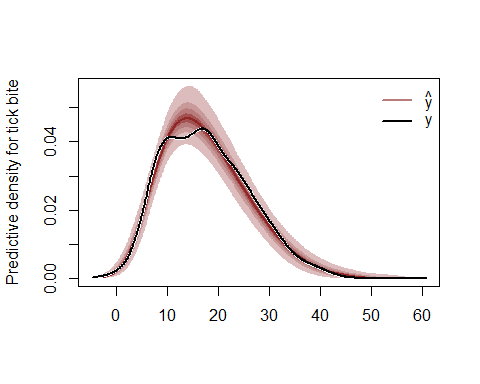

plot_mvgam_ppc(trends_mod3, series = 4, type = "density")

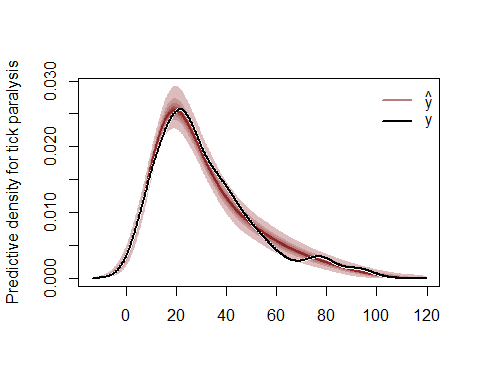

Look at traceplots for the smoothing parameters (rho)

plot_mvgam_trace(object = trends_mod3, param = "rho")

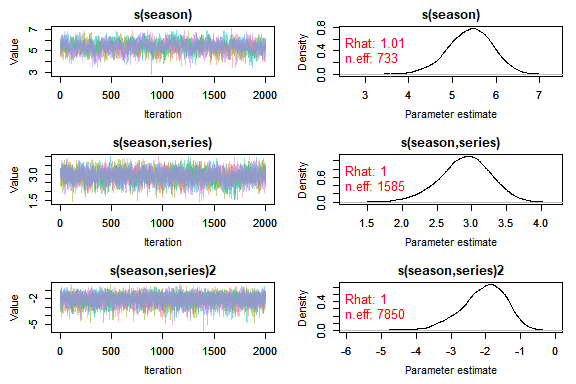

Plot posterior predictive distributions for the training and testing periods for each series

```
plot_mvgam_fc(object = trends_mod3, series = 1,
    data_test = trends_data$data_test)
```

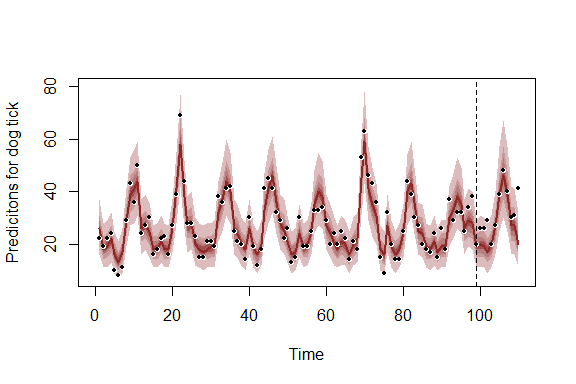

```
plot_mvgam_fc(object = trends_mod3, series = 2,
    data_test = trends_data$data_test)
```

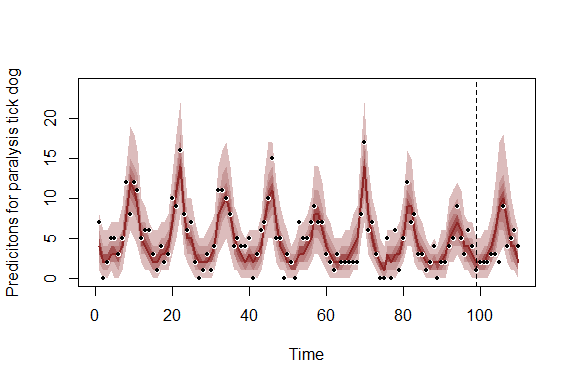

```
plot_mvgam_fc(object = trends_mod3, series = 3,
    data_test = trends_data$data_test)
```

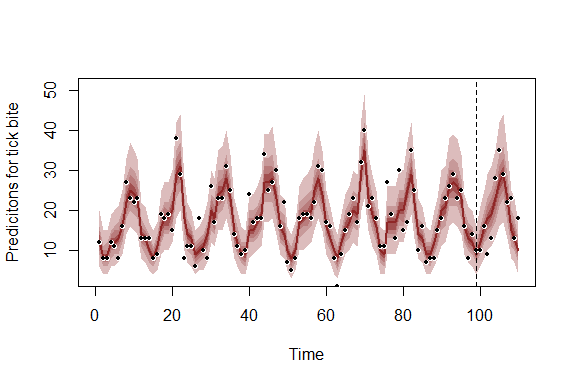

```
plot_mvgam_fc(object = trends_mod3, series = 4,
    data_test = trends_data$data_test)
```

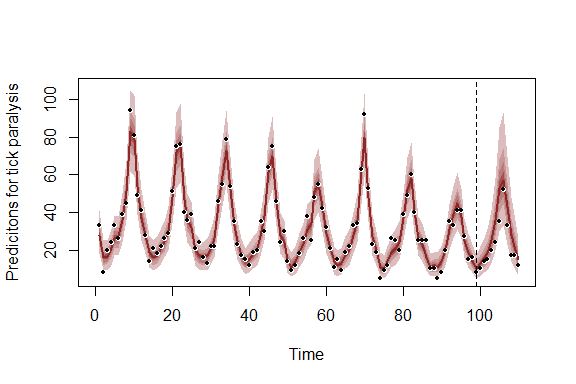

Plot posterior distributions for the latent trend estimates, again for the training and testing periods

```
plot_mvgam_trend(object = trends_mod3, series = 1,
    data_test = trends_data$data_test)
```

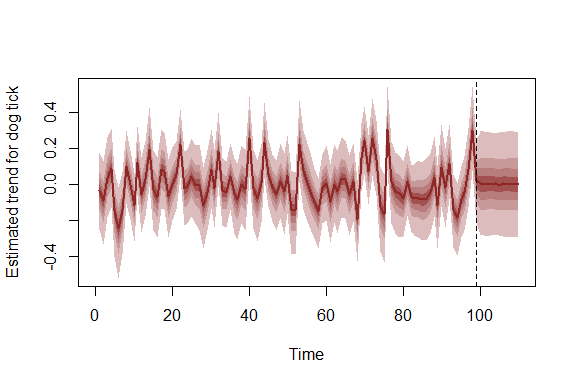

```
plot_mvgam_trend(object = trends_mod3, series = 2,
    data_test = trends_data$data_test)
```

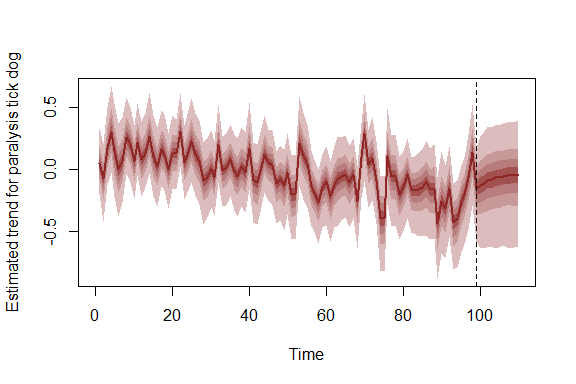

```
plot_mvgam_trend(object = trends_mod3, series = 3,
    data_test = trends_data$data_test)
```

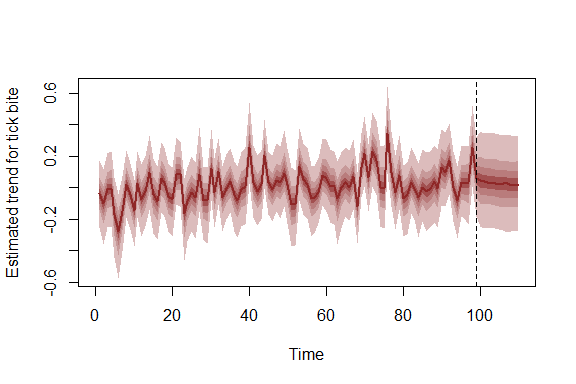

```
plot_mvgam_trend(object = trends_mod3, series = 4,
    data_test = trends_data$data_test)
```

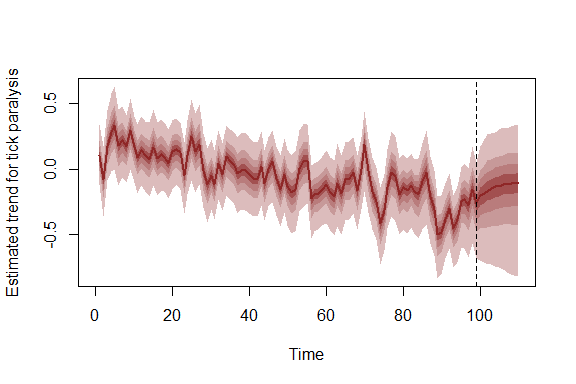

Given that we fit a model with hierarchical seasonality, the seasonal smooths are able to deviate from one another (though they share the same wiggliness and all deviate from a common 'global' seasonal function)

```
plot_mvgam_smooth(object = trends_mod3, series = 1,
    smooth = "season")
```

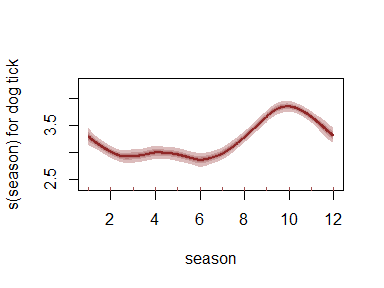

```
plot_mvgam_smooth(object = trends_mod3, series = 2,
    smooth = "season")
```

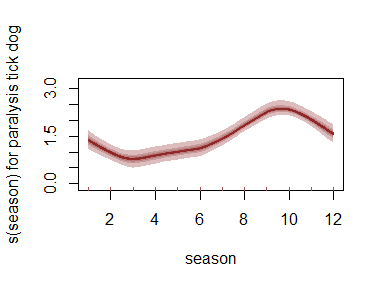

```
plot_mvgam_smooth(object = trends_mod3, series = 3,
    smooth = "season")
```

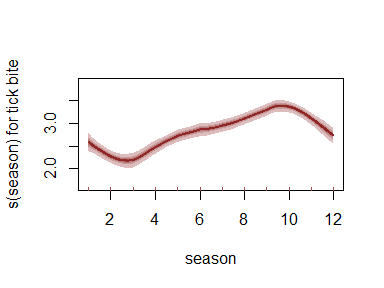

```
plot_mvgam_smooth(object = trends_mod3, series = 4,
    smooth = "season")
```

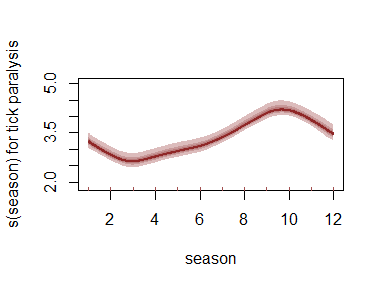

Plot posterior mean estimates of latent trend correlations. These correlations are more useful than looking at latent factor loadings, for example to inspect ordinations. This is because the orders of the loadings (although constrained for identifiability purposes) can vary from chain to chain

```
correlations <- lv_correlations(object = trends_mod3)
library(ggplot2)
mean_correlations <- correlations$mean_correlations
mean_correlations[upper.tri(mean_correlations)] <- NA
mean_correlations <- data.frame(mean_correlations)
ggplot(mean_correlations %>% tibble::rownames_to_column("series1") %>%
    tidyr::pivot_longer(-c(series1), names_to = "series2",
        values_to = "Correlation"), aes(x = series1,
    y = series2)) + geom_tile(aes(fill = Correlation)) +
    scale_fill_gradient2(low = "darkred",
        mid = "white", high = "darkblue",
        midpoint = 0, breaks = seq(-1, 1,
            length.out = 5), limits = c(-1,
            1), name = "Trend\ncorrelation") +
    labs(x = "", y = "") + theme_dark() +
    theme(axis.text.x = element_text(angle = 45,
        hjust = 1))
```

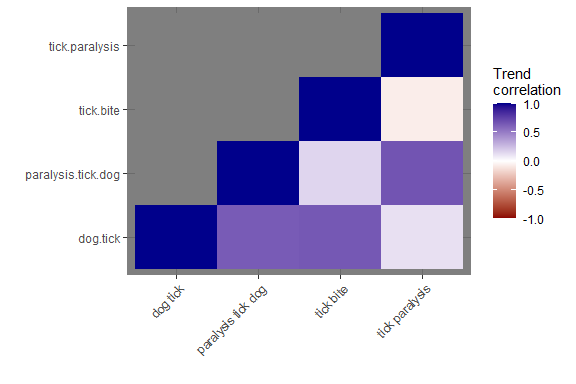

There is certainly some evidence of positive trend correlations for a few of these search terms, which is not surprising given how similar some of them are and how closely linked they should be to interest about tick paralysis in Queensland. Plot some STL decompositions of these series to see if these trends are noticeable in the data

```
plot(stl(ts(as.vector(series$`tick paralysis`),
    frequency = 12), "periodic"))
```

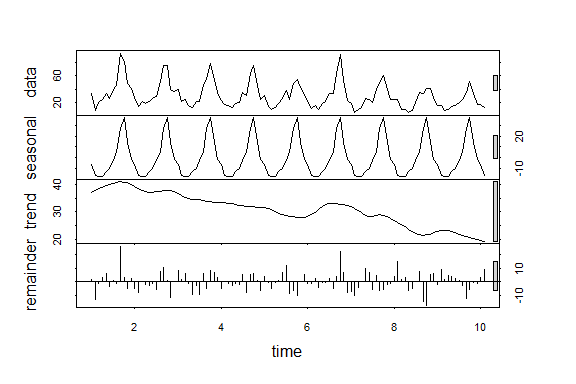

```
plot(stl(ts(as.vector(series$`paralysis tick dog`),
    frequency = 12), "periodic"))
```

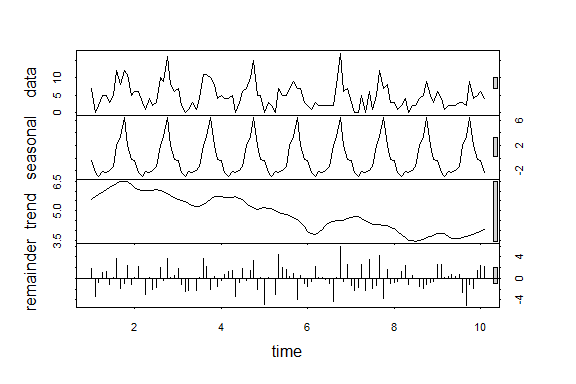

```
plot(stl(ts(as.vector(series$`dog tick`),
    frequency = 12), "periodic"))
```

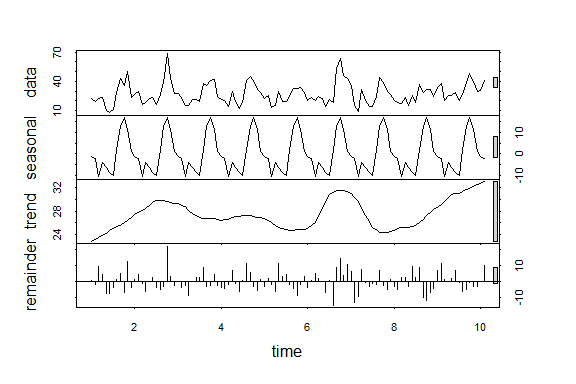

```
plot(stl(ts(as.vector(series$`tick bite`),
    frequency = 12), "periodic"))
```

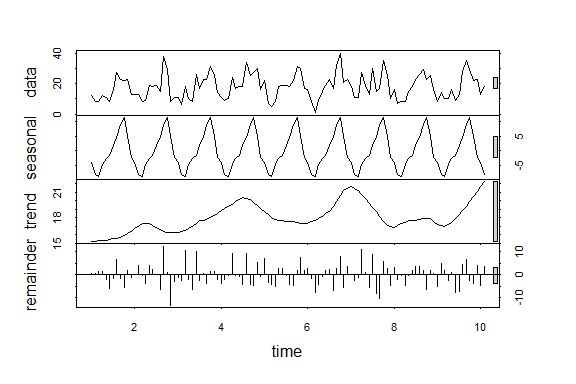

Forecast period posterior predictive checks suggest that the model still has room for improvement:

```
plot_mvgam_ppc(trends_mod3, series = 1, type = "density",
    data_test = trends_data$data_test)
```

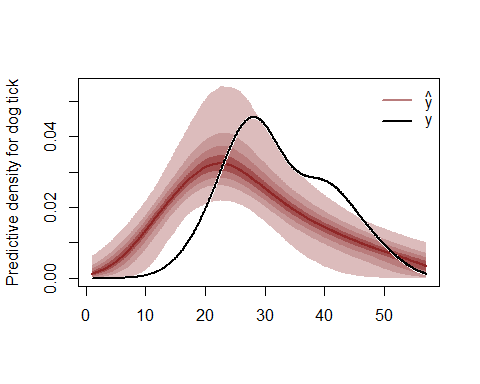

```
plot_mvgam_ppc(trends_mod3, series = 1, type = "mean",
    data_test = trends_data$data_test)
```

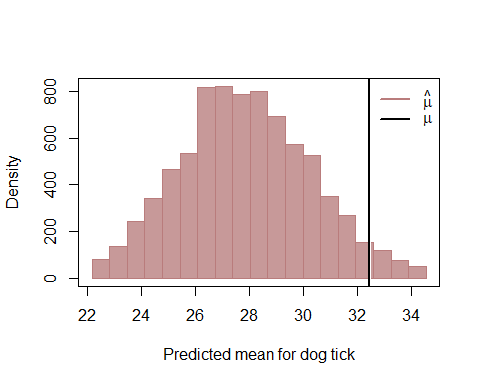

```
plot_mvgam_ppc(trends_mod3, series = 2, type = "density",
    data_test = trends_data$data_test)
```

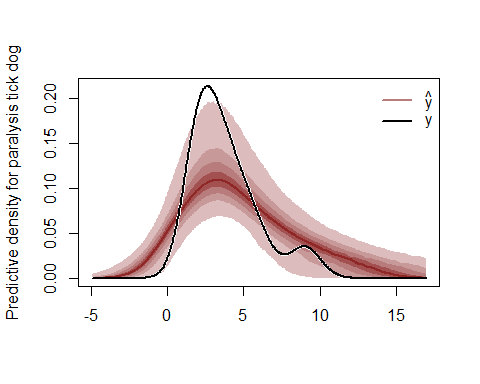

```
plot_mvgam_ppc(trends_mod3, series = 2, type = "mean",
    data_test = trends_data$data_test)
```

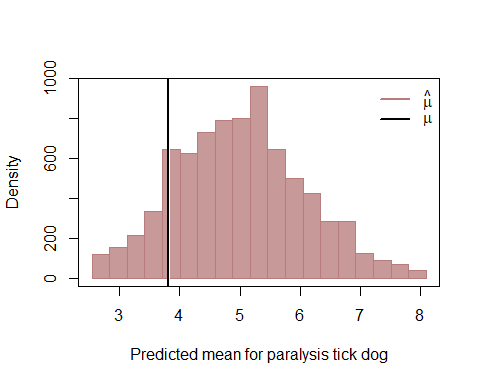

```
plot_mvgam_ppc(trends_mod3, series = 3, type = "density",
    data_test = trends_data$data_test)
```

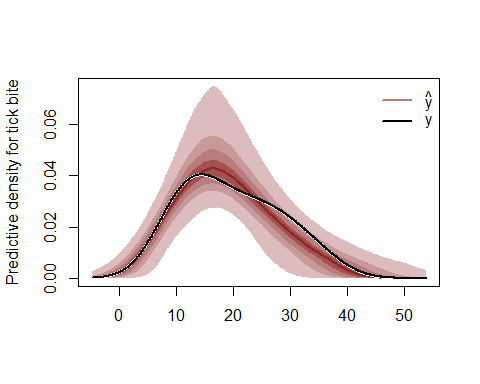

```
plot_mvgam_ppc(trends_mod3, series = 3, type = "mean",
    data_test = trends_data$data_test)
```

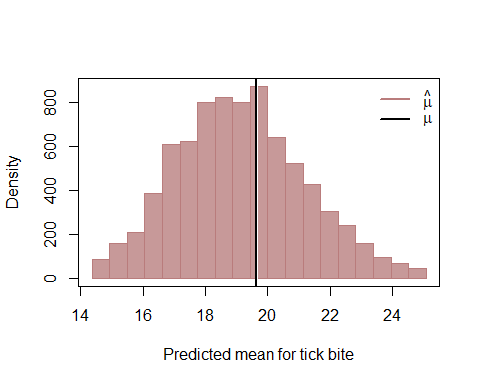

```
plot_mvgam_ppc(trends_mod3, series = 4, type = "density",
    data_test = trends_data$data_test)
```

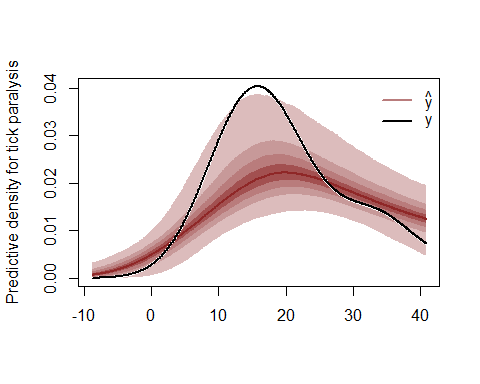

```
plot_mvgam_ppc(trends_mod3, series = 4, type = "mean",
    data_test = trends_data$data_test)
```

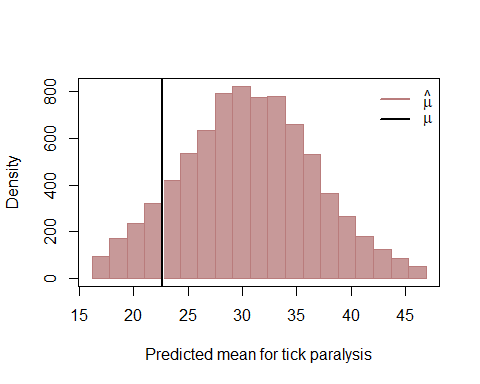

Other next steps could involve devising a more goal-specific set of posterior predictive checks (see [this paper by Gelman et al](https://www.jstor.org/stable/2680852?seq=1#metadata_info_tab_contents) and [relevant works by Betancourt](https://betanalpha.github.io/assets/case_studies/principled_bayesian_workflow.html) for examples) and compare out of sample Discrete Rank Probability Scores for this model and other versions for the latent trends (i.e. AR2, AR3, Random Walk)
