## Appendix S3 for "Dynamic Generalised Additive Models (DGAM) for forecasting discrete ecological time series"

**APPENDIX S3.** Distributed lag dynamic GAMs using the mvgam package for R

Nicholas Clark and Konstans Wells

Here we will use the mvgam package, which fits dynamic GAMs using MCMC sampling via the JAGS software (Note that JAGS 4.3.0 is required; installation links are found [here](https://sourceforge.net/projects/mcmc-jags/files/)), to estimate paramaters of a Bayesian distributed lag model. These models are used to describe simultaneously non-linear and delayed functional relationships between a covariate and a response, and are sometimes referred to as exposure-lag-response models.

Load the Portal rodents capture data, which are available from the portalr package

```
# devtools::install_github('nicholasjclark/mvgam')
library(mvgam)
library(dplyr)
portal_dat <- read.csv("https://raw.githubusercontent.com/nicholasjclark/mvgam/master/NEON_manuscript/Case studies/rodents_data.csv",
    as.is = T)
```

We'll keep data from the year 2004 onwards to make the model quicker to estimate for this simple example

```
portal_dat_all <- portal_dat %>% dplyr::filter(year >=
    2004) %>% dplyr::group_by(year, month) %>%
    dplyr::slice_head(n = 1)
```

Below is an exact reproduction of Simon Wood's lag matrix function (which he uses in his distributed lag example from his book [Generalized Additive Models - An Introduction with R 2nd edition](https://www.taylorfrancis.com/books/mono/10.1201/9781315370279/generalized-additive-models-simon-wood)). Here we supply a vector and specify the maximum lag that we want, and it will return a matrix of dimension length(x) * lag. Note that NAs are used for the missing lag values at the beginning of the matrix. In essence, the matrix objects represent exposure histories, where each row represents the lagged values of the predictor that correspond to each observation in y

```
lagard <- function(x, n.lag = 6) {
    n <- length(x)
    X <- matrix(NA, n, n.lag)
    for (i in 1:n.lag) X[i:n, i] <- x[i:n -
        i + 1]
    X
}
```

Organise all data needed for modelling into a list. We will focus only on the species *Chaetodipus penicillatus* (labelled as PP), which shows reasonable seasonality in its captures over time

```
data_all <- list(lag = matrix(0:5, nrow(portal_dat_all),
    6, byrow = TRUE), y = portal_dat_all$PP,
    season = portal_dat_all$month, year = portal_dat_all$year,
    series = rep(as.factor("series1"), nrow(portal_dat_all)))
data_all$precip <- lagard(portal_dat_all$precipitation)
data_all$mintemp <- lagard(portal_dat_all$mintemp)
```

The exposure history matrix elements of the data list look as follows:

```
head(data_all$lag, 5)
##      [,1] [,2] [,3] [,4] [,5] [,6]
## [1,]    0    1    2    3    4    5
## [2,]    0    1    2    3    4    5
## [3,]    0    1    2    3    4    5
## [4,]    0    1    2    3    4    5
## [5,]    0    1    2    3    4    5
head(data_all$precip, 5)
##      [,1] [,2] [,3] [,4] [,5] [,6]
## [1,] 37.8   NA   NA   NA   NA   NA
## [2,]  8.7 37.8   NA   NA   NA   NA
## [3,] 43.5  8.7 37.8   NA   NA   NA
## [4,] 23.9 43.5  8.7 37.8   NA   NA
## [5,]  0.9 23.9 43.5  8.7 37.8   NA
head(data_all$mintemp, 5)
##        [,1]   [,2]   [,3]   [,4]  [,5] [,6]
## [1,] -9.710     NA     NA     NA    NA   NA
## [2,] -5.924 -9.710     NA     NA    NA   NA
## [3,] -0.220 -5.924 -9.710     NA    NA   NA
## [4,]  1.931 -0.220 -5.924 -9.710    NA   NA
## [5,]  6.568  1.931 -0.220 -5.924 -9.71   NA
```

All other elements of the data list are in the usual vector format

```
head(data_all$y, 5)
## [1]  0  1  2 NA 10
head(data_all$series, 5)
## [1] series1 series1 series1 series1 series1
## Levels: series1
head(data_all$year, 5)
## [1] 2004 2004 2004 2004 2004
```

View the raw series. There is a clear seasonal pattern to the data, and there are missing values scattered throughout

```
plot(ts(data_all$y, frequency = 12), ylab = "Captures for PP",
    xlab = "")
```

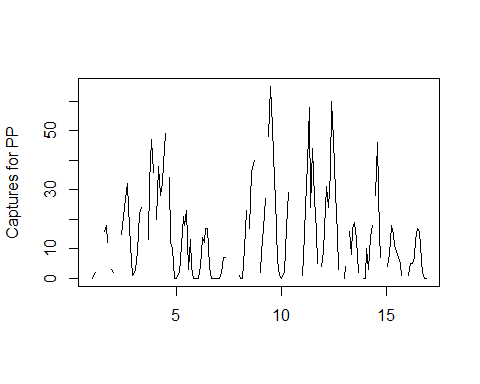

Create training and testing sets; start at observation 7 so that the NA values at the beginning of the covariate lag matrices are not included. Currently there is no option for on-the-fly imputation of missing covariate values in mvgam models, though this can easily be done in JAGS by specifying prior distributions over these missing entries

```
data_train <- list(lag = data_all$lag[7:174,
    ], y = data_all$y[7:174], series = data_all$series[7:174],
    season = data_all$season[7:174], year = data_all$year[7:174],
    precip = data_all$precip[7:174, ], mintemp = data_all$mintemp[7:174,
        ])
data_test <- list(lag = data_all$lag[175:length(data_all$y),
    ], y = data_all$y[175:length(data_all$y)],
    series = data_all$series[175:length(data_all$y)],
    season = data_all$season[175:length(data_all$y)],
    year = data_all$year[175:length(data_all$y)],
    precip = data_all$precip[175:length(data_all$y),
        ], mintemp = data_all$mintemp[175:length(data_all$y),
        ])
```

Now we can fit a dynamic GAM with distributed lag terms for precipitation and minimum temperature and an AR1 process for the unobserved temporal trend. The distributed lags are set up as tensor product smooth functions (see help(te) for an explanation of tensor product smooths) between lag and each covariate. We assume a Poisson observation model for this example

```
test <- mvjagam(formula = y ~ te(mintemp,
    lag, k = c(8, 4)) + te(precip, lag, k = c(8,
    4)), data_train = data_train, data_test = data_test,
    family = "poisson", chains = 4, burnin = 12000,
    trend_model = "AR1")
## Compiling rjags model...
## Starting 4 rjags simulations using a PSOCK cluster with 4 nodes on host
## 'localhost'
## Simulation complete
## Note: Summary statistics were not produced as there are >50 monitored
## variables
## [To override this behaviour see ?add.summary and ?runjags.options]
## FALSEFinished running the simulation
## NOTE: Stopping adaptation
```

The summary of the model provides useful information on convergence for unobserved parameters

```
summary_mvgam(test)
## GAM formula:
## y ~ te(mintemp, lag, k = c(8, 4)) + te(precip, lag, k = c(8,
##     4))
##
## Family:
## Poisson
##
## N series:
## 1
##
## N observations per series:
## 168
##
## GAM smooth term approximate significances:
##                   edf Ref.df  Chi.sq p-value
## te(mintemp,lag) 14.44  30.00 113.471  <2e-16 ***
## te(precip,lag)  12.05  28.00   0.579       1
## ---
## Signif. codes:  0 '***' 0.001 '**' 0.01 '*' 0.05 '.' 0.1 ' ' 1
##
## GAM coefficient (beta) estimates:
##                           2.5%          50%       97.5% Rhat n.eff
## (Intercept)         1.87349502  2.158565082  2.48593018 1.07   162
## te(mintemp,lag).1  -1.72635111 -0.588609269  0.24387727 1.01   555
## te(mintemp,lag).2  -1.50140131 -0.542324409  0.19098724 1.00   835
## te(mintemp,lag).3  -1.72490919 -0.626636522  0.54321850 1.00  1438
## te(mintemp,lag).4  -1.00228143 -0.432498240  0.32527963 1.01  1059
## te(mintemp,lag).5  -0.41920030  0.119827195  0.72837518 1.01   683
## te(mintemp,lag).6  -0.01518754  0.456791007  1.19271129 1.00   930
## te(mintemp,lag).7  -0.02563295  0.570260141  1.44831577 1.00  3410
## te(mintemp,lag).8  -0.74801138 -0.279608109  0.41230751 1.00  2401
## te(mintemp,lag).9  -0.27079232  0.007239820  0.35727574 1.01   620
## te(mintemp,lag).10 -0.03215606  0.241262367  0.65555584 1.00  1334
## te(mintemp,lag).11 -0.13003169  0.461997868  1.09208654 1.00  5879
## te(mintemp,lag).12 -0.40411502  0.085217864  0.80036458 1.00  1449
## te(mintemp,lag).13 -0.26530675  0.139777806  0.61133070 1.01   537
## te(mintemp,lag).14 -0.11597124  0.261435877  0.84599004 1.00   900
## te(mintemp,lag).15 -0.13477207  0.445841288  1.21149393 1.00  3934
## te(mintemp,lag).16 -0.31028784  0.146119096  0.83094234 1.00  2130
## te(mintemp,lag).17 -0.20390784  0.141165783  0.54730700 1.00   705
## te(mintemp,lag).18 -0.17403630  0.175333437  0.65592247 1.00  1234
## te(mintemp,lag).19 -0.26506427  0.315369768  1.00719294 1.00  4577
## te(mintemp,lag).20 -0.24028992  0.228554400  0.93793113 1.00  1756
## te(mintemp,lag).21 -0.15212865  0.205232897  0.63728519 1.00   814
## te(mintemp,lag).22 -0.26649476  0.102779001  0.61444749 1.00  1248
## te(mintemp,lag).23 -0.49475871  0.089651449  0.83123765 1.00  4007
## te(mintemp,lag).24  0.02215563  0.504591166  1.25709474 1.00  1248
## te(mintemp,lag).25 -0.12901993  0.296686141  0.78603436 1.01   661
## te(mintemp,lag).26 -0.44582261 -0.046816394  0.55775470 1.00   997
## te(mintemp,lag).27 -1.05355244 -0.446429867  0.33356717 1.00  3945
## te(mintemp,lag).28 -0.07456074  0.475337925  1.19073526 1.00  2332
## te(mintemp,lag).29 -0.19643819  0.166243922  0.54665782 1.01  1120
## te(mintemp,lag).30 -0.64539861 -0.262747789  0.13356034 1.00  1680
## te(mintemp,lag).31 -1.47578999 -0.788541250 -0.12458561 1.00  3689
## te(precip,lag).1   -0.19586653  0.005496211  0.16150463 1.00   644
## te(precip,lag).2   -0.24256255 -0.051180983  0.12641354 1.00   611
## te(precip,lag).3   -0.29579642 -0.047526554  0.20051491 1.00  1032
## te(precip,lag).4   -0.02714347  0.116698623  0.28852115 1.00  1256
## te(precip,lag).5   -0.15460637 -0.003450895  0.10476689 1.00   804
## te(precip,lag).6   -0.20713938 -0.053545208  0.08511320 1.00   947
## te(precip,lag).7   -0.27701751 -0.060477009  0.14099278 1.00  1245
## te(precip,lag).8   -0.14727954  0.015995435  0.18937856 1.00  2369
## te(precip,lag).9   -0.16333849 -0.026161825  0.09973932 1.00  1882
## te(precip,lag).10  -0.20126022 -0.044626663  0.10269930 1.00  1622
## te(precip,lag).11  -0.28546308 -0.062678453  0.12658747 1.00  1823
## te(precip,lag).12  -0.29023551 -0.089187569  0.11258210 1.00  1560
## te(precip,lag).13  -0.19213591 -0.043076285  0.10115396 1.00   727
## te(precip,lag).14  -0.18440827 -0.032481051  0.12506420 1.00   793
## te(precip,lag).15  -0.26808041 -0.040657930  0.16209547 1.00  1569
## te(precip,lag).16  -0.39516582 -0.166144887  0.06174874 1.00  1194
## te(precip,lag).17  -0.30491606 -0.116175998  0.07178368 1.00   575
## te(precip,lag).18  -0.28993664 -0.083004855  0.12084254 1.00   794
## te(precip,lag).19  -0.33698114 -0.066241158  0.17647656 1.00  1595
## te(precip,lag).20  -0.16129995  0.040723490  0.24211929 1.00  1072
## te(precip,lag).21  -0.17014045 -0.031111913  0.11224393 1.00   486
## te(precip,lag).22  -0.23108616 -0.084559812  0.05652241 1.00   674
## te(precip,lag).23  -0.23379891 -0.050217236  0.13993977 1.00  1181
## te(precip,lag).24  -0.33754506 -0.075523648  0.10581697 1.00  1768
## te(precip,lag).25  -0.92375325 -0.090271371  0.34620540 1.01  1986
## te(precip,lag).26  -0.60604654 -0.046790524  0.27342490 1.01  1640
## te(precip,lag).27  -0.44298899 -0.011000657  0.33722600 1.01  1662
## te(precip,lag).28  -0.53679705 -0.028710241  0.46845157 1.00  2641
##
## GAM smoothing parameter (rho) estimates:
##                         2.5%       50%    97.5% Rhat n.eff
## te(mintemp,lag)   2.17734261 3.4838673 4.521299    1  1920
## te(mintemp,lag)2  0.20249971 2.4631802 4.029051    1  1778
## te(mintemp,lag)3 -1.92855955 0.4501257 3.371396    1   612
## te(precip,lag)    2.66223047 3.9028886 4.856156    1  4260
## te(precip,lag)2   0.05072617 2.4308375 4.146685    1  1252
## te(precip,lag)3   0.29750697 2.8051368 4.327658    1  2336
##
## Latent trend drift (phi) and AR parameter estimates:
##          2.5%       50%     97.5% Rhat n.eff
## phi 0.0000000 0.0000000 0.0000000  NaN     0
## ar1 0.7544379 0.8594178 0.9437071    1  3018
## ar2 0.0000000 0.0000000 0.0000000  NaN     0
## ar3 0.0000000 0.0000000 0.0000000  NaN     0
##
```

As with all other mvgam objects, we can create plots of the estimated forecast distribution and the dynamic latent trend process

```
plot_mvgam_fc(test, series = 1, data_test = data_test,
    ylim = c(0, 100))
```

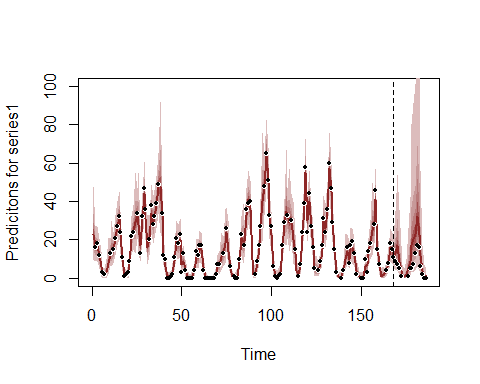

plot_mvgam_trend(test, series = 1, data_test = data_test)

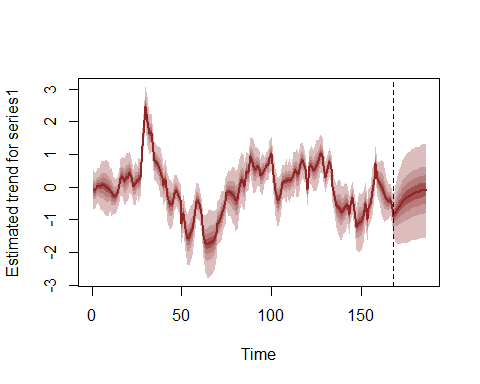

Traceplots of smooth penalties and latent trend parameters indicate good mixing and convergence of the four MCMC chains

plot_mvgam_trace(test, "rho")

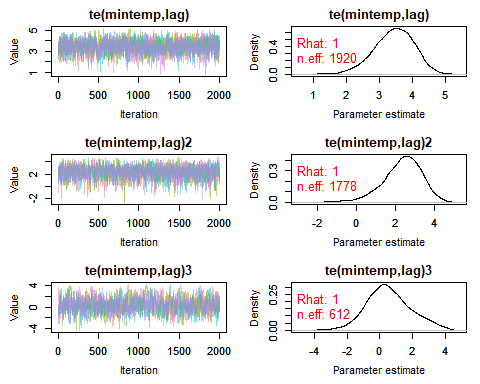

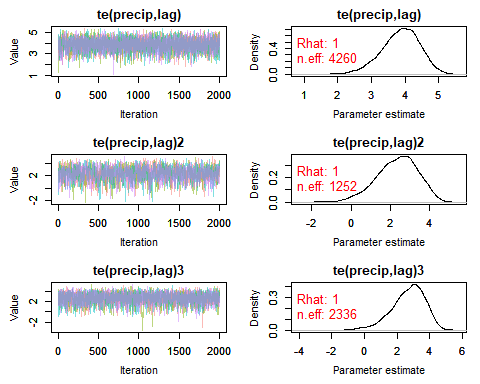

plot_mvgam_trace(test, "trend")

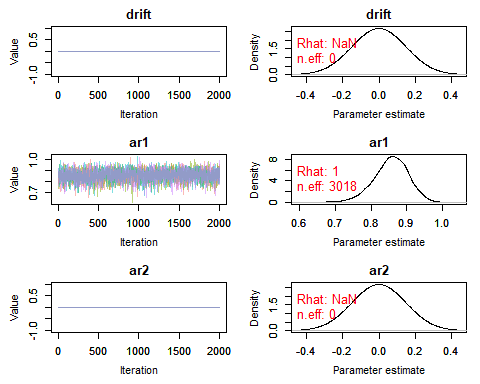

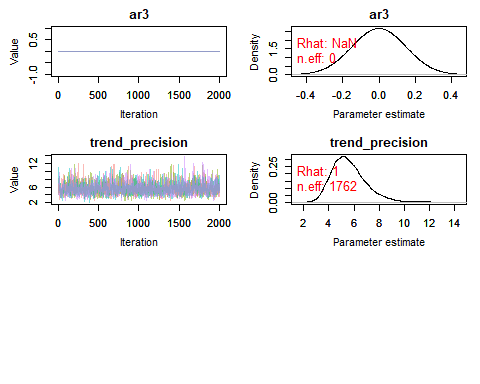

We can also create quick plots of the estimated smooth tensor product interactions for the distributed lag terms, which basically follow mgcv's two-dimensional plotting utility but uses the mvgam's estimated coefficients

plot_mvgam_smooth(test, series = 1, smooth = 1)

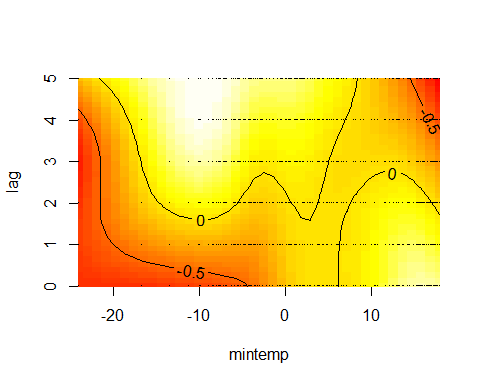

plot_mvgam_smooth(test, series = 1, smooth = 2)

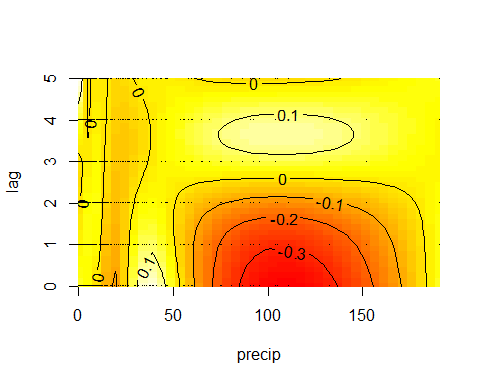

If you are like me then you'll find these plots rather difficult to interpret! The more intense yellow/white colours indicate higher predicted values, with the deeper red colours representing lower predicted values, but actually making sense of how the functional response is expected to change over different lags is not easy from these plots. HOwever, we can use the predict_mvgam function to generate much more interpretable plots. First we will focus on the effect of mintemp and generate a series of predictions to visualise how the estimated function changes over different lags. Set up prediction data by zeroing out all covariates apart from the covariate of interest

```
newdata <- data_test
newdata$year <- rep(0, length(newdata$year))
newdata$season <- rep(0, length(newdata$season))
newdata$precip <- matrix(0, ncol = ncol(newdata$precip),
    nrow = nrow(newdata$precip))
```

Set up viridis plot colours and initiate the plot window to be centred around zero. We will then keep all mintemp values at zero apart from the particular lag being predicted so that we can visualise how the predicted function changes over lags of mintemp. Predictions are generated on the link scale in this case, though you could also use the response scale. Note that we need to first generate predictions with all covariates (including the mintemp covariate) zeroed out to find the 'baseline' prediction so that we can shift by this baseline for generating a zero-centred plot. That way our resulting plot will roughly follow the traditional mgcv partial effect plots

```
cols <- viridis::inferno(6)
plot(1, type = "n", xlab = "Mintemp", ylab = "Predicted response function",
    xlim = c(min(data_train$mintemp), max(data_train$mintemp)),
    ylim = c(-1.6, 1.6))

# Calculate predictions for when mintemp
# is all zeros to find the baseline value
# for centring the plot
newdata$mintemp <- matrix(0, ncol = ncol(newdata$mintemp),
    nrow = nrow(newdata$mintemp))
preds <- predict_mvgam(test, series = 1,
    newdata = newdata, type = "link")
offset <- mean(preds)

for (i in 1:6) {
    # Set up prediction matrix for mintemp
    # with lag i as the prediction sequence;
    # use a sequence of mintemp values across
    # the full range of observed values in
    # the training data
    newdata$mintemp <- matrix(0, ncol = ncol(newdata$precip),
        nrow = nrow(newdata$precip))
    newdata$mintemp[, i] <- seq(min(data_train$mintemp),
        max(data_train$mintemp), length.out = length(newdata$year))

    # Predict on the link scale and shift by
    # the offset so that values are roughly
    # centred at zero
    preds <- predict_mvgam(test, series = 1,
        newdata = newdata, type = "link") -
        offset

    # Calculate empirical prediction
    # quantiles
    probs = c(0.05, 0.2, 0.3, 0.4, 0.5, 0.6,
        0.7, 0.8, 0.95)
    cred <- sapply(1:NCOL(preds), function(n) quantile(preds[,
        n], probs = probs))

    # Plot expected function posterior
    # intervals (40-60%) and medians in
    # varying colours per lag
    pred_upper <- cred[4, ]
    pred_lower <- cred[6, ]
    pred_vals <- seq(min(data_train$mintemp),
        max(data_train$mintemp), length.out = length(newdata$year))
    polygon(c(pred_vals, rev(pred_vals)),
        c(pred_upper, rev(pred_lower)), col = scales::alpha(cols[i],
            0.6), border = scales::alpha(cols[i],
            0.7))
    lines(pred_vals, cred[5, ], col = scales::alpha(cols[i],
        0.8), lwd = 2.5)
}
abline(h = 0, lty = "dashed")
legend("topleft", legend = paste0("lag",
    seq(0, 5)), bg = "white", bty = "n",
    col = cols, lty = 1, lwd = 6)
```

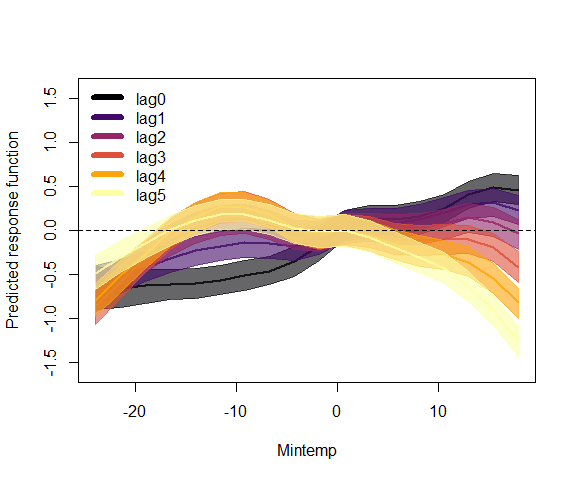

This plot demonstrates how the effect of mintemp is expected to change over different exposure lags, with the 3 - 5 month lags showing more of a cyclic seasonal pattern (catches expected to increase in the summer and autumn, roughly 3 - 5 months following cold minimum winter temperatures) while the recent lags (lags 0 and 1) demonstrate a more linear response function (catches broadly increasing as minimum temperature increases). This is hopefully a useful example for developing a better understanding of how a distributed lag model is attempting to recreate the data generating process. And here is the same plot for precipitation, which demonstrates how a u-shaped functional relationship diminishes toward a flat function at lags 2 - 5 (though this effect is clearly less important in the model than the mintemp * lag effect above)

```
newdata <- data_test
newdata$year <- rep(0, length(newdata$year))
newdata$season <- rep(0, length(newdata$season))
newdata$mintemp <- matrix(0, ncol = ncol(newdata$mintemp),
    nrow = nrow(newdata$mintemp))
newdata$precip <- matrix(0, ncol = ncol(newdata$precip),
    nrow = nrow(newdata$precip))
preds <- predict_mvgam(test, series = 1,
    newdata = newdata, type = "link")
offset <- mean(preds)
plot(1, type = "n", xlab = "Precipitation",
    ylab = "Predicted response function",
    xlim = c(min(data_train$precip), max(data_train$precip)),
    ylim = c(-1.6, 1.6))

for (i in 1:6) {
    newdata$precip <- matrix(0, ncol = ncol(newdata$precip),
        nrow = nrow(newdata$precip))
    newdata$precip[, i] <- seq(min(data_train$precip),
        max(data_train$precip), length.out = length(newdata$year))
    preds <- predict_mvgam(test, series = 1,
        newdata = newdata, type = "link") -
        offset
    probs = c(0.05, 0.2, 0.3, 0.4, 0.5, 0.6,
        0.7, 0.8, 0.95)
    cred <- sapply(1:NCOL(preds), function(n) quantile(preds[,
        n], probs = probs))
    pred_upper <- cred[4, ]
    pred_lower <- cred[6, ]
    pred_vals <- seq(min(data_train$precip),
        max(data_train$precip), length.out = length(newdata$year))
    polygon(c(pred_vals, rev(pred_vals)),
        c(pred_upper, rev(pred_lower)), col = scales::alpha(cols[i],
            0.6), border = scales::alpha(cols[i],
            0.7))
    lines(pred_vals, cred[5, ], col = scales::alpha(cols[i],
        0.8), lwd = 2.5)
}
abline(h = 0, lty = "dashed")
legend("bottomleft", legend = paste0("lag",
    seq(0, 5)), bg = "white", bty = "n",
    col = cols, lty = 1, lwd = 6)
```

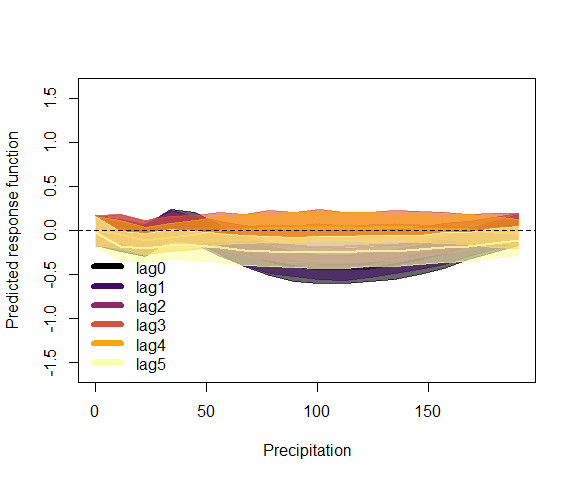

All of the usual functions in mvgam can also be used for list data objects, so long as they contain the necessary fields series, season and year. For example, posterior retrodictive checks for the in-sample training period:

plot_mvgam_ppc(test, series = 1, type = "cdf")

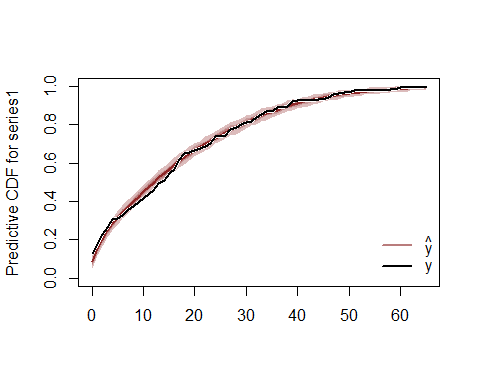

and predictive checks for the out of sample forecast period (which demonstrates how the model overpredicts for the forecast period in this particular example):

```
plot_mvgam_ppc(test, data_test = data_test,
    series = 1, type = "cdf")
```

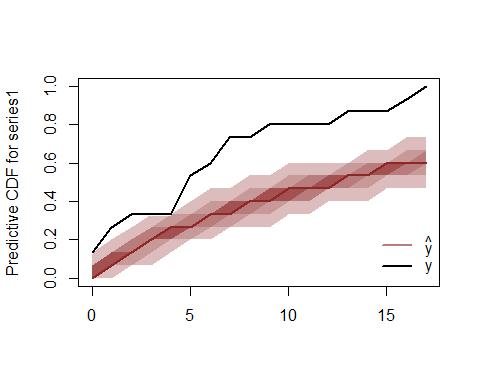

Logical next steps for interrogating this model would be to trial different trend types (random walk, AR2, AR3), replace the distributed lag function for precip with a standard smooth function (that does not include lag interactions, as clearly the model above indicates that these are not supported) and inspect whether different covariates (such as ndvi or maxtemp) might play a role in modulating catches of PP. Finally, once we are satisfied that we have a well-performing model that we can understand and interrogate, we could expand up to a multivariate model by including other species as response variables. This would allow us to capture any possible unobserved dependencies in the catches of multiple co-occurring species in a single unified modelling framework
